# Origin-1: a generative AI platform for *de novo* antibody design against novel epitopes

**DOI:** 10.64898/2026.01.14.699389

**Authors:** Simon Levine, Jonathan Edward King, Jacob Stern, David Grayson, Raymond Wang, Rui Yin, Umberto Lupo, Paulina Kulytė, Ryan Matthew Brand, Tristan Bertin, Robert Pfingsten, Jovan Cejovic, Chelsea Chung, Breanna K. Luton, Andrew Hagemann, Robel Haile, Elliot Medina, Pankaj Panwar, Oleksii Dubrovskyi, Chase LaCombe, Zahra Anderson, Derrik Mildh, Scott Benjamin, Joe Kaiser, Joseph Ferron, Marta Sarrico, Alexandria Kershner, Apurva Mishra, Kai R. Ejan, Emily K. Marsh, Paul Bringas, Phetsamay Vilaychack, Kyra Chapman, Jacob Ripley, Muttappa Gowda, Kathryn M. Collins, Cailen M. McCloskey, Jeremiah S. Joseph, Rylee Ripley, Shaheed A. Abdulhaqq, David Spencer, Tiffany DeVine, Audree Feltner, Michael Guerin, Jeffrey Goby, Jesse Hendricks, Danielle Castillo, Sean McClain, Douglas Ganini, Derek Shpiel, James Mategko, Eder Cruz Garcia, Masoud Zabet-Moghaddam, John M. Sutton, Zheyuan Guo, Sean M. West, Janani S. Iyer, Amir Shanehsazzadeh

**Affiliations:** Absci Corporation; Opilio LLC

## Abstract

Generative artificial intelligence has advanced antibody discovery, yet *de novo* design of therapeutic antibodies against targets with “zero-prior” epitopes remains a fundamental challenge. We define “zero-prior” epitopes as target sites lacking structural data from any reported antibody-antigen or protein-protein complex involving the target. Here we present Origin-1, a generative AI platform that overcomes this by integrating epitope-conditioned all-atom structure generation, paired complementarity determining region sequence design, and a specialized co-folding-based scoring protocol to select antibody designs predicted to be high-confidence, specific binders with favorable developability. We evaluated Origin-1 on a panel of ten targets selected to have no available protein–protein complex structures and minimal homology (≤60% sequence identity) to proteins with known complexes, creating stringent design conditions. In fewer than one hundred design attempts per target, we identified developable, specific antibodies, validated across multiple biophysical and developability assays, for four targets: COL6A3, AZGP1, CHI3L2, and IL36RA, with functional inhibition demonstrated for IL36RA. Cryogenic electron microscopy confirmed the atomic accuracy of our designs, revealing complexes that closely matched the computational models with high structural fidelity (3.0-3.3 Å resolution; 0.83-0.91 DockQ). Furthermore, we employed AI-guided affinity maturation to optimize a *de novo* antibody binder against IL36RA, producing functional antagonists with sub-nanomolar affinities and a top EC_50_ of 12.3 nM. These results demonstrate a framework for targeting epitopes without structural precedent, expanding the programmable therapeutic antibody landscape.

## 1 Introduction

Antibodies serve as the immune system’s primary defense mechanism, recognizing and neutralizing pathogens with exceptional specificity. Their capacity to bind diverse antigens with high affinity has established antibodies as essential therapeutic agents [1, 2, 3]. However, despite their success, traditional antibody discovery methods remain resource-intensive and provide limited control over which epitopes are targeted. This limitation motivates the development of computational approaches for efficient *de novo* antibody design with precise epitope specification.

Antibody-antigen recognition is primarily mediated by six complementarity-determining regions (CDRs), whose high sequence and conformational variability make CDR design the core computational challenge [4, 5]. Therapeutic antibody design typically begins with known framework scaffolds that have favorable developability, stability, and expression properties, and then focuses on designing CDRs that bind a target epitope. A common computational paradigm addresses this in two stages: first generating atomistic antibody-antigen complex structures conditioned on the antigen and epitope, then designing CDR sequences predicted to fold into these structures.

Recent advances in machine learning have enabled progress in protein structure prediction [6, 7] and *de novo* design [8, 9], yet existing approaches exhibit significant limitations for epitope-targeted antibody design. Structure generation methods face several challenges: backbone-only diffusion approaches such as RFdiffusion [9] and RFantibody [10] lack atomic resolution and treat antigens rigidly; general-purpose all-atom design models including P(all-atom) [11] and La-Proteina [12] are not fine-tuned for antibody-specific conditioning or epitope-targeted generation; and hallucination-based approaches such as BoltzDesign [13], BindCraft [14], Germinal [15], and mBER [16] leverage gradient signals from trained structure prediction models for optimization but incur substantial computational expense and often necessitate subsequent sequence redesign or complex loss function composition to avoid degenerate outputs.

Sequence design models also remain limited in their applications to epitope-specific antibody design. Inverse folding models such as ProteinMPNN [17] and antibody-specific variants such as AbMPNN [18] achieve strong performance but treat heavy and light chains independently. Although hybrid approaches that combine structure-based models with protein language models have shown promise [19–22] and paired antibody language models [21] enable heavy-light chain co-modeling, these methods do not deeply integrate paired language representations with geometric structure encoding in a unified multi-modal framework for therapeutic CDR design.

Other antibody modeling approaches that report successful epitope-specific design provide limited detail on their technical methodology; thus, effectively evaluating them for their advances or limitations remains difficult [23–27]. The recently open-sourced BoltzGen [28], developed independently and concurrently with this work, addresses several key limitations—providing all-atom resolution and binding site conditioning through a flexible design specification language—though experimental validation for antibodies is limited to nanobodies, leaving full-length antibody design with experimental validation unaddressed. These gaps motivate approaches that combine epitope-conditioned all-atom structure generation with paired-chain sequence design in addition to transparency in development methodology.

When applying protein structure and sequence design tools to problems in drug discovery, it is important to not only produce accurate designs, but also stringently filter outputs such that a minimal, high-confidence set can be advanced to the laboratory for *in vitro* experimentation [29]. This “design-and-score” paradigm uses generative models to design putative binders to targets of interest and subsequently leverages scoring metrics to filter and rank designs for downstream experimentation. Prior work has shown that the confidence metrics output from folding models such as AlphaFold-Multimer [30] can accurately evaluate the likelihood that a sequence will adopt a particular structural geometry [30–32]. However, other studies have shown that these folding models perform poorly for antibody-antigen complex structures specifically [33, 34], with models failing to recover correct poses when provided with antibody-antigen complex sequences [2]. Therefore, beyond addressing limitations in protein structure and sequence design to achieve accurate epitope-targeted CDR design, further innovation is required in folding model methodology to apply the associated confidence metrics to antibody design scoring.

To these ends, we introduce Origin-1, a design-and-score AI platform that addresses the aforementioned limitations to achieve epitope-specific *de novo* antibody design via a two-stage framework. The first stage, AbsciGen, enables site-specific, conditional CDR design. The second stage, AbsciBind, addresses the limitations of traditional folding-based confidence metrics to accurately evaluate antibody-antigen complex designs and select the best candidates for downstream experimentation. We use this platform to generate developable and functional full-length monoclonal antibodies (mAbs) against four human protein targets (COL6A3, AZGP1, CHI3L2, and IL36RA) in fewer than one hundred attempts per target. Of note, these mAbs were designed against “zero-prior” epitopes – that is, epitopes that were selected as putative functional binding sites without guidance from solved complex structures. These results show that AI-based approaches can succeed not only in designing antibody binders against structurally resolved protein-protein interfaces [23–27], but also user-specified novel interfaces, which may unlock access to disease targets that have historically been difficult to drug or otherwise remain understudied.

## 2 Methods

### 2.1 Overview of Origin-1

Origin-1 is a generative AI platform that designs and scores putative antibody binders to target proteins of interest. To achieve this outcome, Origin-1 performs two key tasks: 1) generates the structures and corresponding amino acid sequences of antibodies that are likely binders to proteins of interest, and 2) scores the resulting candidates to select high-confidence designs to prioritize for *in vitro* experimentation. We refer to the protocols used to perform these tasks as AbsciGen and AbsciBind, respectively.

In the sections that follow, we describe the methodology underlying these strategies and the experimental approaches we used to assess their joint performance in **Figure 1**.

**Figure 1.**
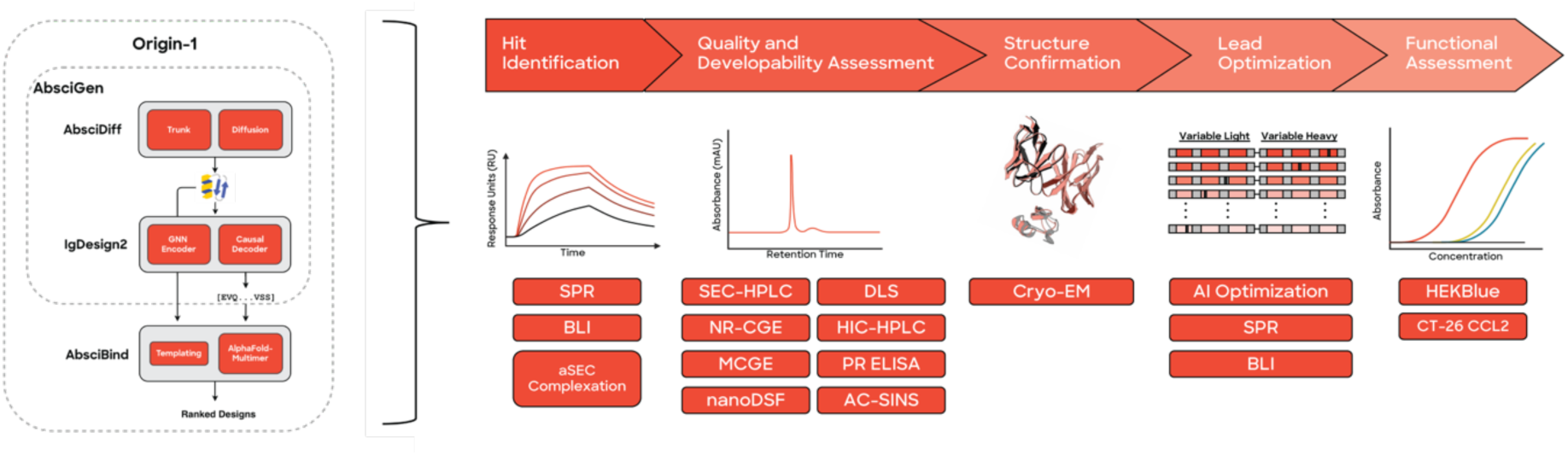
Overview of the Origin-1 platform and experimental validation cascade. (Left) Overview of Origin-1 Platform. AbsciGen generates antibody designs through two stages: AbsciDiff, which produces all-atom antibody-antigen complex structures via diffusion-based generation, and IgDesign2, which designs CDR sequences. AbsciGen designs are evaluated by AbsciBind, which identifies high-confidence binders to reduce experimental screening demands. (Right) Overview of experimental cascade. Antibody designs are expressed as full-length monoclonal antibodies and are evaluated through stringent validation experiments to confirm specificity, selectivity, and epitope-specific binding. Initial hits are identified by SPR and confirmed by BLI and aSEC complexation experiments. Hits are assessed for quality and developability, including purity, aggregation, thermal stability, polyreactivity, and hydrophobicity. Antibody-antigen complexes are formed and analyzed by cryo-EM to confirm agreement between *in silico* designs and experimentally determined structures. Some hits are optimized through synthesis of model-ranked mutational variants (e.g., Origin-1), with affinity confirmed by SPR and BLI. Top parent and variant designs are evaluated in HEKBlue assays to confirm functional activity in human cells and an in-house CT-26 CCL2 assay to confirm functional activity in mouse cells. SPR = Surface Plasmon Resonance; BLI = Biolayer Interferometry; aSEC = Analytical Size Exclusion Chromatography; SEC-HPLC = Size Exclusion Chromatography – High-Performance Liquid Chromatography; NR-CGE = Non-Reduced Microchip Capillary Gel Electrophoresis; MCGE = Microchip Capillary Gel Electrophoresis; nanoDSF = Nanoscale Differential Scanning Fluorimetry; DLS = Dynamic Light Scattering; HIC-HPLC = Hydrophobic Interaction Chromatography - High-Performance Liquid Chromatography; PR ELISA = Polyreactivity Enzyme-Linked Immunosorbent Assay; AC-SINS = Affinity-Capture Self-Interaction Nanoparticle Spectroscopy; Cryo-EM = Cryogenic Electron Microscopy. Created using Biorender.com.

### 2.2 Origin-1 Datasets and Curation

To train AbsciGen we rely on AbData, a protein structure curation workflow and database that we developed specifically for training antibody design models. In brief, the AbData pipeline comprises three stages: 1) Protein Data Bank (PDB) [35] sequence and structure information extraction, in addition to structure cleaning and resolution of missing residues and atoms; 2) antibody-antigen complex or protein-protein dimeric interface metadata extraction; and 3) structure dataset creation, dataset splitting, data deduplication, and final annotation of structure interfaces, complementarity determining regions (CDRs), and frameworks. This pipeline captured data from all bioassemblies and asymmetric units in the PDB, resulting in antibody structures from 10,045 distinct PDBs, including all structures in the Structural Antibody Database (SAbDab) [36] as well as many rare categories of antibodies that are incompletely captured in SAbDab [37, 38]. We also devised an inspection process that automatically flagged over 1400 candidate entries with potential errors arising from spurious bioassemblies, false symmetries, or artifactual chain duplication. When errors were confirmed, individual complexes were corrected by removing erroneous chains or reverting to the asymmetric unit. Our protein-protein interaction dataset largely follows the logic outlined in Townshend et al. 2019 [39], where interfaces are extracted for all contacting chain-pairs across the PDB. Additional details on AbData’s methodology are reported in Supplement §7.1.

### 2.3 Origin-1 Models

Origin-1 generates antibody designs via AbsciGen, comprising AbsciDiff for structure generation (§2.3.1, Supplement §7.2) and IgDesign2 for sequence design (§2.3.2). It scores designs using AbsciBind (§2.3.3).

#### 2.3.1 Structure Design via AbsciDiff

AbsciDiff (**Figure 2**) is a diffusion-based all-atom generative model fine-tuned from Boltz-1 [40] for epitope-conditioned antibody design. Key modifications include antibody- and docking-specific feature masking and conditioning strategies, an intermediate sequence hypothesis module with recycling, integration of optimized CuEquivariance kernels [41], and support for structural templates. The sampling procedure follows Boltz-1, preserving the sample efficiency and stability of the parent model. In the following sections, we describe the diffusion formulation, feature generation strategy, architectural changes, and training protocol.

**Figure 2.**
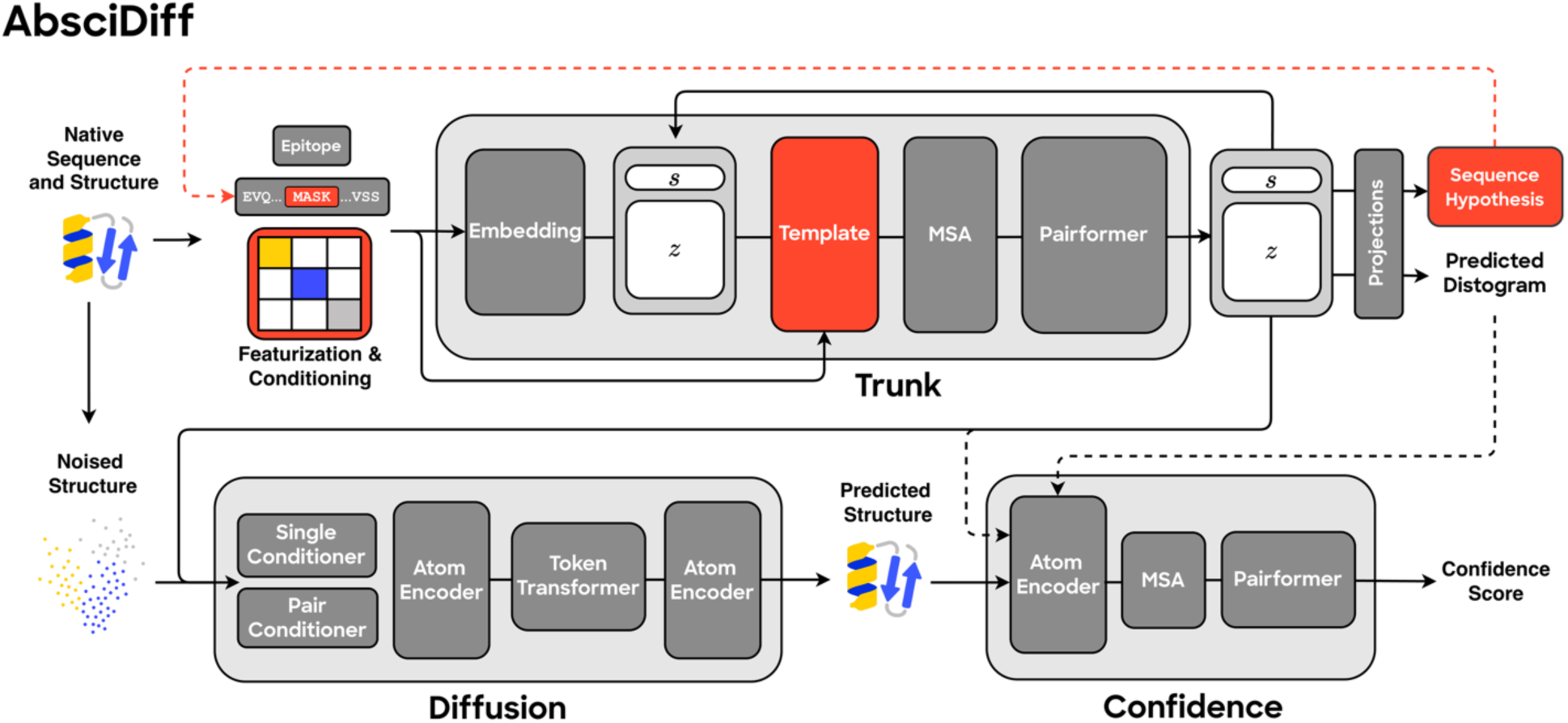
Schematic illustrating the AbsciDiff model. Building on the architecture of Boltz-1, AbsciDiff uses a pairformer-based trunk to provide conditioning information to a diffusion module. Modifications include invariant feature conditioning via an AlphaFold3-like template module [6], as well as a module to predict a sequence hypothesis. The sequence hypothesis is used to update the model’s input features for a second forward pass through the model. Dashed lines indicate a stopped gradient. Red indicates modifications to Boltz-1.

#### AbsciDiff Diffusion Formulation

Following Boltz-1, we model the noise added to the native structure of an antibody-antigen complex *X*_0_ ∈ ℝ^*N* ×3^ through a forward diffusion process,

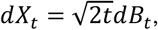

where *dB_t_* is (*N* × 3)-dimensional Brownian motion.

To generate new complexes, we reverse this diffusion process over 𝑇 timesteps (**Figure 3**). Starting from *X_T_* ∼ 𝒩(0, *I*), we iteratively denoise using a model *P_θ_* parametrized by a neural network and conditioned on antigen and framework structure templates *X_a_*, *X_f_*, sequences *S_a_*, *S_f_*, and epitope residues 𝐸*_a_*. We denote the full conditioning information as

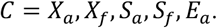

**Figure 3.**
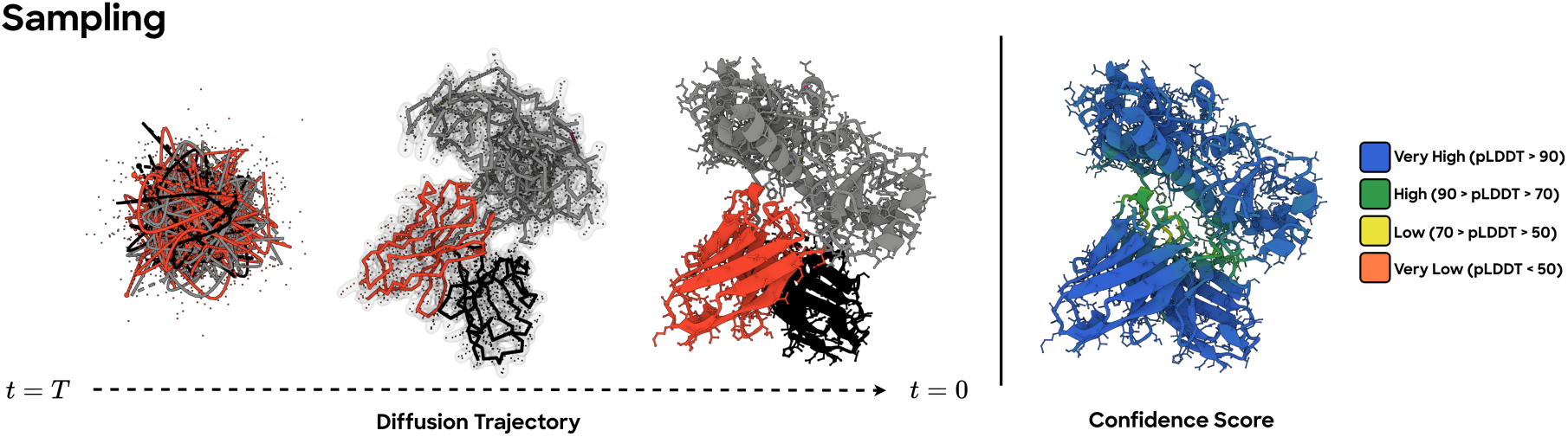
Reverse diffusion trajectory for joint antibody–antigen structure generation. Starting from a fully corrupted state *X_T_* (left), AbsciDiff performs iterative denoising transitions *p*(*X_t_*|*X*_*t*–1_,*t*) (middle), progressively recovering global geometry and local structure until producing a final denoised sample *X*_0_ (center right). Subsequently predicted confidence computed on the final structure is shown on the far right (color scaled by residue pLDDT, where blue represents high confidence and red represents low confidence).

We train this model to approximate the expected coordinates of the entire original complex structure under the defined diffusion process given the current timestep and all available conditioning information:

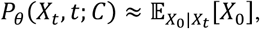

with the standard denoising loss

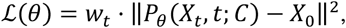

where *w_t_* is a weight proportional to the variance of the noising process at timestep *t*.

#### AbsciDiff Featurization

##### Antigen, Framework, and Epitope Conditioning

AbsciDiff’s most significant departure from Boltz-1 concerns feature masking and conditioning. For our intended use, we assume antigen structures are known, and we aim to design or redesign only the CDR regions while keeping therapeutic framework sequences and structures intact. We additionally condition on an epitope defined during training as the set of antigen residues within 6 Å of the antibody, down-sampled via a geometric distribution (*p* = 0.3) for robustness. We mask sequence and structure features for all CDR positions, as well as all inter-chain pair features. Docking is thus guided only by the token-wise binary epitope vector, implemented analogously to Boltz-1’s pocket-conditioning feature.

##### Design Region Representation

Amino acid identities in the design region are initially set to the unknown (UNK) amino acid token for both sequence input and reference conformer lookup. During structure prediction, the model is trained to perform atom superposition, as in P(all-atom) [11]. For this task, all amino acids in the design region are represented in Atom14 notation, with excess “virtual” atoms placed at the location of their residue’s Cα (**Supplementary Figure 1**). Amino acids outside the design region have no virtual atoms since their identities are known *a priori*. Other than the sequence hypothesis head described below, AbsciDiff performs no sequence decoding on designed amino acids.

##### Template Structures

AbsciDiff also adds support for structural template featurization in two ways: *endogenous* templating (in which pairwise residue distance information is derived from a user-provided structure) and *exogenous* templating (where the same information is sourced from external template databases). In both cases, structural information is encoded and embedded as in AlphaFold3. During development, we found that endogenous templating was practical and effective at providing structural guidance without introducing bias. Thus, the final training runs included only endogenous template information.

##### Multiple Sequence Alignments (MSAs)

AbsciDiff supports MSA featurization as in Boltz-1, though it remains disabled by default. We found that full MSA feature sets performed no better than a one-hot encoding of the input sequence when conditioned on endogenous templates.

#### AbsciDiff Trunk Module and Intermediate Sequence-Informed Design

An auxiliary prediction head in AbsciDiff produces a design region sequence hypothesis from the final single (*s*) representation of the trunk. The predicted logits are decoded into the most likely amino acid tokens and, along with corresponding atom-level reference conformer features, are recycled through the trunk with gradient flow stopped. Sequence prediction is trained using cross-entropy loss against the native sequence. We hypothesize that intermediate sequence prediction and conformer recycling assists the design process by focusing the search space of potential residue identities and local atomic conformations prior to the diffusion model performing final design.

#### AbsciDiff Confidence Module and Design Pre-Ranking

Though the underlying confidence module architecture is the same as Boltz-1, AbsciDiff introduces a new on-model ranking and filtering strategy. Like Boltz-1, the diffusion module emits *M* samples per sampling pass, which are subsequently scored using the confidence module to generate predicted Template Modeling scores (pTM), predicted Local Distance Difference Test scores (pLDDT), predicted Docking Error (pDE), and predicted Aligned Error (pAE). We normalize these scores (noting that pDE and pAE are unbounded and must be transformed and inverted) before averaging to produce a composite ranking score by which the top *k* ≤ *M* candidates are selected. Although the confidence module is efficient — and its scores well-correlated with supervised metrics - we do not rely on it alone for evaluating real-world binding affinity. As such, we employ this on-model scoring strategy for efficient *pre-ranking*, thereby reducing the downstream computational cost of re-folding and scoring designs with AbsciBind.

#### AbsciDiff Training

##### Fine-tuning

We initialize AbsciDiff using the weights from the provided Boltz-1 checkpoint and fine-tune the model on a subset of AbData. As described earlier, redundant dataset entries were filtered based on sequence similarity thresholds, and data were split using a combination of temporal and sequence similarity-based criteria. Training follows Boltz-1 defaults with the number of learning rate steps reduced to 20 and an effective batch size of 64 for 10 epochs.

##### Cropping

Input structures are spatially cropped to a maximum of 512 residues, with larger crop sizes showing minimal improvement on sampled antibody-antigen interface quality. The cropping strategy centers the representation on the antibody-antigen interface and requires that: (1) the variable antibody region (Fv) is always included; (2) antibody constant domains are always excluded; (3) the remaining budget is allocated to antigen residues in order of the smallest distance to the epitope.

#### 2.3.2 Sequence Design via IgDesign2

##### IgDesign2 Problem Formulation

We model the joint probability of the unknown sequence region *R* using an autoregressive factorization:

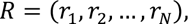

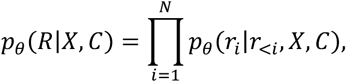

where each amino acid *r_i_* is conditioned on previously generated positions *r*_<*i*_, backbone structure *X*, and known sequence context *C*. Our goal is to find the neural network parameters *θ* that maximize the likelihood of the training data under this scheme.

##### IgDesign2 Model Architecture

IgDesign2 combines a Graph Neural Network (GNN) encoder, a causal transformer decoder, and a protein language model (pLM) refinement module into a structure-conditioned sequence design system (Figure 4). The input to the encoder consists of the four-atom representation of the protein backbone [***N*, *C_a_*, *C*, *0***] residues in which each residue constitutes a node in a graph with *k* edges connecting to its nearest neighbors in Euclidean space. We leverage PiFold’s [42] node and edge featurization scheme and mirror their GNN message passing architecture for encoding the three-dimensional geometry of each residue based on its atomic coordinates. The decoder ingests the structure embeddings and any antigen and antibody framework sequence context for conditioning the autoregressive CDR sequence generation process, which is applied in a randomly shuffled decoding order via temperature-weighted sampling or beam search.

**Figure 4.**
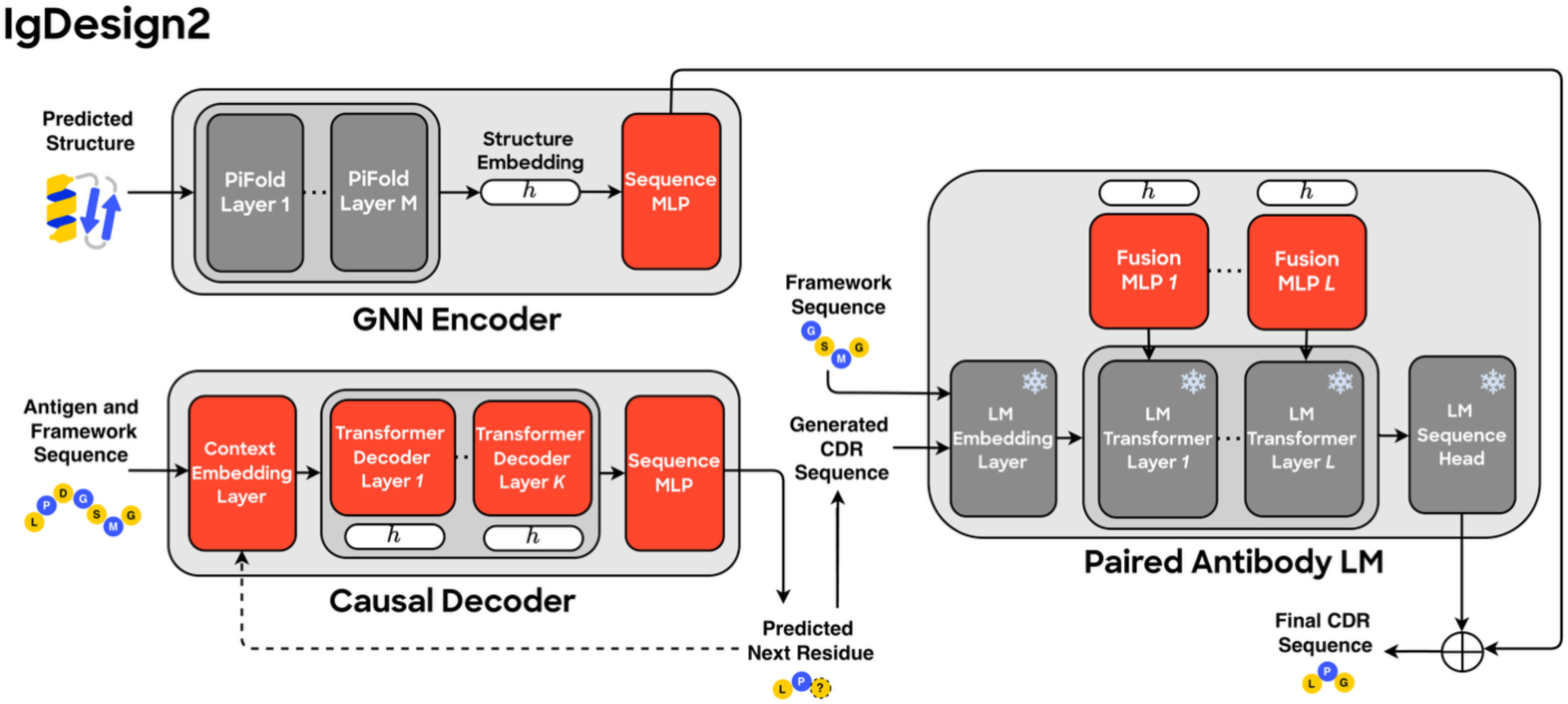
Schematic illustrating IgDesign2 architecture for CDR sequence design. A GNN encoder processes antibody-antigen structure features, followed by a causal transformer decoder for autoregressive sequence generation. The decoder supports both temperature-weighted sampling (for diversity) and beam search (for high-likelihood sampling). Generated sequences are refined by a structure-aware paired antibody language model. Red indicates model components developed specifically for this work. Snowflakes indicate frozen pre-trained modules. Solid arrows show forward information flow. Dotted arrows show recycled information flow.

The heavy and light chain sequences are then provided to the IgBert paired antibody language model [22] for structure-aware refinement. At every layer of the pLM’s transformer, the pLM latent embeddings are fused with the final structure embedding in a shared low-dimensional space before being projected back to the pLM dimension for processing by the subsequent transformer layer [20, 43, 44]. After the transformer has fully processed the fused representation, the final latent representations of both the GNN and pLM are decoded by their respective sequence prediction heads and integrated with a residual connection to produce the final sequence output. This jointly optimized “generate-and-refine” approach combines the strengths of both sequence modeling paradigms to enhance the quality of our conditional CDR designs.

##### IgDesign2 Pre-Training

We pre-train the encoder and decoder on protein-protein interaction examples from AbData, spatially cropped in three dimensions around the interfacial region of the two chains to a maximum of 500 total residues. We use the Adam optimizer with a learning rate of 10^-3^, *β*_1_ = 0.9, and *β*_2_ = 0.999, along with an effective batch size of 32 and a standard cross-entropy loss over 20 possible amino acids. Early stopping is applied when the loss has not improved for 10 consecutive epochs on the held-out validation set. We retain the published hyperparameters of the original PiFold model for the encoder (dimension 128, 10 layers, 4 heads, 30 nearest neighbors in the structure graph), while the causal transformer decoder consists of 10 standard transformer decoder layers with four attention heads and a feedforward dimension of 512. We also apply Gaussian noise with a standard deviation of 0.1 Å to the coordinates of each input structure during training to encourage robustness.

##### IgDesign2 Fine-Tuning

We fine-tune the model with the same hyperparameters, aside from two adjustments: we reduce the effective batch size to 8 and terminate training when the cross-entropy loss over the masked region has not improved for 5 consecutive epochs on the held-out validation set. Antibody-antigen complexes are spatially cropped to a maximum of 600 residues around the interacting residues of the antigen while retaining the full antibody Fv. The pLM weights remain frozen throughout fine-tuning, while a trainable multi-layer perceptron (MLP) is introduced per pLM layer to fuse the incoming projections from PiFold and the pLM. Each MLP consists of three standard linear layers of dimension 128, with ReLU activation functions. The incoming projections from PiFold and the pLM reduce their respective embeddings to 64 dimensions prior to concatenation, while the final projection maps the 128-dimensional MLP output back to the pLM embedding dimension. We compute the cross-entropy loss over both the sequence produced by the causal decoder and the sequence refined by the pLM, summing them to compute the total loss.

#### 2.3.3 Design Filtering via AbsciBind

We create a protocol, AbsciBind, as a scoring method for antibody-antigen complexes, leveraging the advantages of existing co-folding-based scoring approaches and identifying workarounds to their limitations [6, 7, 33, 34]. We provide extended details on the methodology underlying AbsciBind in Supplement §7.5. In brief, AbsciBind is a derivation of AF_Unmasked [45], where the associated interface-predicted template modeling (ipTM) score is computed with greater awareness of the relative arrangements of antibody heavy and light chains in addition to antigen chains. We benchmark the resulting AbsciBind ipTM Score against several existing comparable ipTM scores [27] via a binder vs. non-binder discrimination task in an experimental set of eight antibody-antigen systems [21, 46]. Results show that AbsciBind achieves the strongest average binder vs. non-binder discrimination in seven of eight targets (**Figure 5**) when the maximum ipTM score across all five AlphaFold-Multimer (AFM) v2.3 model checkpoints is used for assessment.

**Figure 5.**
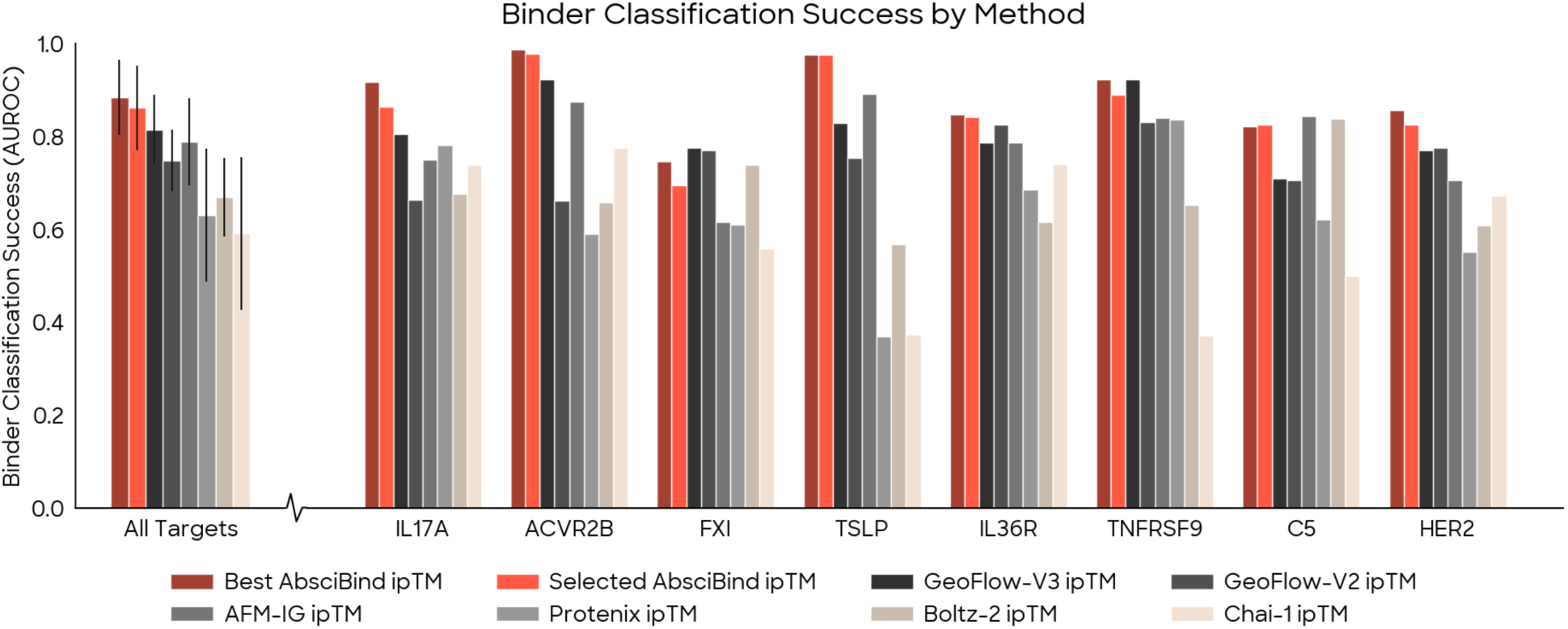
The AbsciBind protocol ipTM score outperforms comparable alternatives for evaluating antibody-antigen complexes. Evaluation was performed on an experimental set of eight antibody–antigen systems [21, 46]. All AbsciBind approaches were evaluated using ipTM scores to ensure comparability with previously reported results. AUROC values for GeoFlow-V3, GeoFlow-V2, AFM-IG, Protenix, Boltz-2 and Chai-1 were sourced from BioGeometry et al. 2025 [27]. Best AbsciBind ipTM is the maximum ipTM score across all five AFM v2.3 model checkpoints for a given design. Selected AbsciBind ipTM uses the ipTM score from AFM v2.3 Model 2 only (`model_2_multimer_v3` checkpoint).

Importantly, using only Model 2 yields an approximately 80% reduction in runtime compared with evaluating designs with all five models. Given this substantial runtime reduction and the strong classification performance, Model 2 is used to support the AbsciBind protocol for the remainder of the present effort.

We define a final AbsciBind Score as the mean of the default AbsciBind protocol ipTM score (computed over all antibody and antigen chains and interfaces) and the Antibody-Aligned ipTM score (Supplement §7.5). We design this metric to integrate a global interface score with an antibody-aligned, consistently normalized assessment of the antibody-antigen interface’s quality. We use this score to evaluate AbsciGen designs for *de novo* library design (§2.4.3) and to select mutant variants for lead optimization efforts (§2.4.4).

#### 2.3.4 *In silico* Benchmarking of Origin-1

We benchmark AbsciGen against RFantibody [10], a diffusion-based antibody design method selected for its experimentally-validated design pipeline. To ensure a fair comparison, we modify RFantibody’s input pre-processing to initialize idealized coordinates at the origin for all CDR residues (Supplement §7.3). All other RFantibody settings remain at their default values. For AbsciGen, we disable the beam search feature of IgDesign2 and instead use autoregressive sampling to encourage greater sequence diversity. All other design choices remain as described below and in the Methods.

Our objective was to evaluate AbsciGen and RFantibody in their respective abilities to design antibodies against epitopes that lacked associated target structures in complex with antibody or protein binders (“zero-prior” epitopes). To this end, we applied both approaches to design CDR regions using multiple frameworks, CDR lengths, and putative epitopes. We then assessed the quality of generated designs with unsupervised metrics. We used AbsciBind Score (§2.3.3) to compare AbsciGen and RFantibody design performance. Of note, AbsciGen was developed without the explicit goal of maximizing AbsciBind Score. We also computed Observed Antibody Space identity search (OASis [47]) scores associated with AbsciGen vs. RFantibody sequences to compare the humanness achieved via both approaches.

##### Test Set

We test AbsciGen and RFantibody against ten targets with no antibody-antigen complex structures in our training data (**Table 1**). The epitope selection process is described in §2.4.2.

**Table 1.**
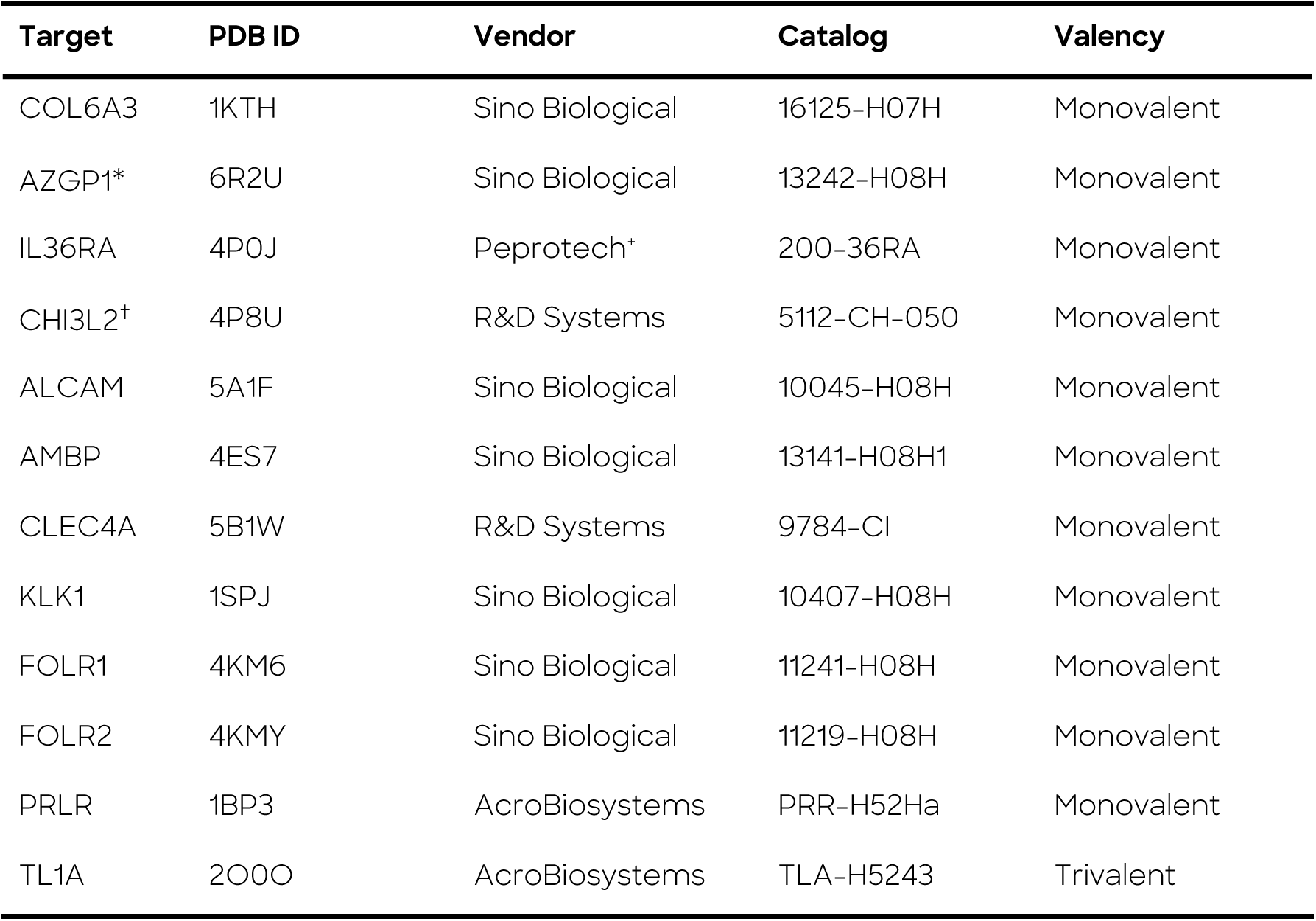
List of targets used for Origin-1 experimental validation. *There is one PDB entry of AZGP1 in complex with a non-antibody protein (PDB:3ES6) that was missed during selection due to an error in the PDB bioassembly; thus, for AZGP1 we required the model to identify epitopes that were distinct from the solved interface in this PDB entry. ^+^Peprotech is now owned and operated by Thermo Fisher Scientific. ^†^We additionally produced CHI3L2 internally for downstream validation experiments as described in §7.8.8.

##### Frameworks

For each target, we employ three well-characterized therapeutic antibody frameworks: trastuzumab, relatlimab, and sotrovimab. This standard design strategy mitigates the risk of memorizing native antibody-antigen pairs and ensures robust starting scaffolds with established developability profiles.

##### Design Specification

We generate diverse design specifications by varying HCDR3 length (8–26 residues), LCDR3 length (8–10 residues), and epitope subsampling (retaining 45–90% of identified epitope residues). Epitope subsampling reflects uncertainty in epitope identification and encourages exploration of diverse binding modes within the target region. This procedure yielded 102 distinct design specifications per target. For each design specification, we generate one structure and eight sequences.

### 2.4 Experimental Validation of Origin-1

To experimentally test Origin-1’s ability to design full-length monoclonal antibodies in a low-throughput setting, we selected fewer than one hundred designs per target and ordered them as monoclonal antibodies for low-throughput *in vitro* screening. Designs that demonstrated successful binding in primary surface plasmon resonance (SPR) screens were further validated using biolayer interferometry (BLI), complexation analyses, developability assessments, and functional assays. Where appropriate, cryogenic electron microscopy (Cryo-EM) was used for structure confirmation (**Figure 1**). Together, these approaches allowed us to assess AbsciGen’s ability to design high quality binders to novel binding locations, and AbsciBind’s ability to select winning candidates from amongst these designs.

#### 2.4.1 Target Selection

We selected targets to test Origin-1’s performance by prioritizing entries with high structural resolution (<3.5 Å) and few missing/unresolved residues, in addition to targets for which antigen reagents were commercially procurable (**Table 1**). To extend beyond what has been demonstrated in recent reports of successful *de novo* antibody design [23–27], we additionally confirmed that the PDB did not contain structures of antibodies or other proteins in complex with these targets, requiring Origin-1 to identify novel binding interfaces (“zero-prior epitopes”), rather than providing Origin-1 with known binding interfaces to increase the likelihood of successful binder design. We further increased the complexity of the challenge by selecting targets that maintained limited sequence overlap (≤60% identity by MMseqs2 [48]) with any protein for which a protein-protein complex structure existed in the PDB, ensuring that the selected targets were understudied and required novel epitope identification. This approach allowed us to evaluate Origin-1’s generalizability. We experimentally validated Origin-1 on ten targets meeting these criteria (**Table 1**).

IL36RA was of particular interest given its role as an anti-inflammatory cytokine that inhibits binding of the proinflammatory cytokines IL36α, IL36β, and IL36γ to the IL36 receptor (IL36R). This axis plays a complex role in some cancers where IL36β and IL36γ are believed to promote inflammation and therefore promote anti-tumor immune response, which would be dampened by IL36RA [49]. Inhibition of IL36RA may enhance IL-36R signaling, and could promote immune infiltration into previously “immune-cold” tumors.

#### 2.4.2 Epitope selection, framework selection, and CDR length determination

##### Epitope selection

To select epitopes per target, we surveyed the literature to assess each target’s biological role, surface geometry, chemical properties, and any reported interactions with other proteins, peptides, or small molecule ligands. We prioritized solvent-accessible surface regions maintaining curvature (i.e., knobs or holes), hydrophobic core patches surrounded by hydrophilic/nucleophilic residues, and structured regions as opposed to disordered/loop regions (**Figure 6**). For example, for IL36RA we identified a putative biologically relevant epitope to enable downstream functional characterization of any binders. In particular, we used the complex structure of A-552, a small molecule antagonist of IL36γ, [50] from PDB:6P9E to convert a functional binding pocket into a putative functional epitope of IL36γ, a homolog of IL36RA. We then mapped this putative epitope onto IL36RA via structural homology.

**Figure 6.**
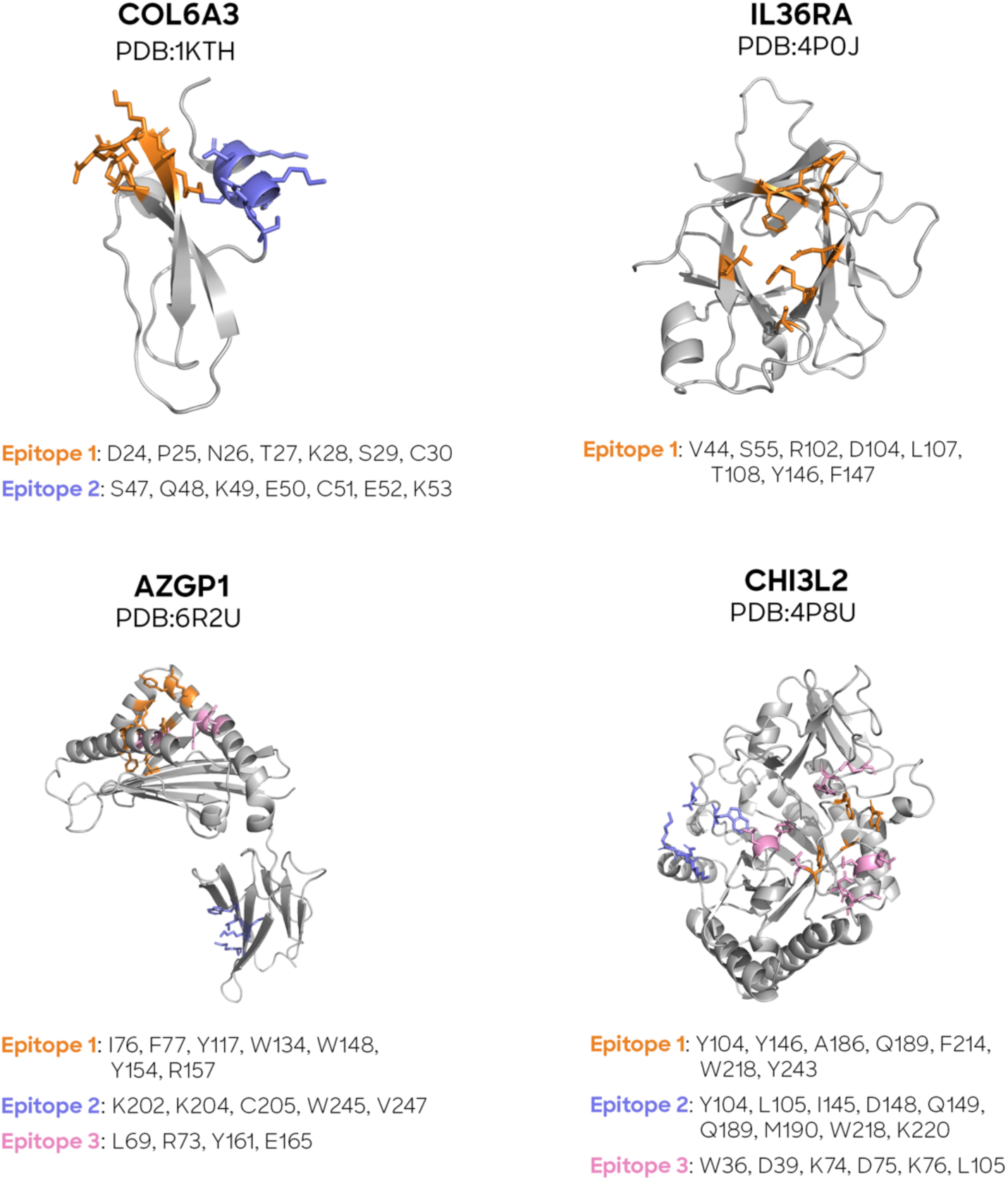
List of residues selected per epitope per target for which Origin-1 binders were confirmed.

During inference, the epitope feature was subsampled from the full epitope as follows. For a single input sample, the number of epitope feature residues was sampled uniformly between 0.45 and 0.9 times the number of full epitope residues, with a minimum of four epitope feature residues. The epitope feature positions were then sampled without replacement from the full epitope until this number was reached.

##### Framework Selection

We selected four antibody frameworks (FWRs) from clinically approved therapeutic antibodies with diverse germlines, Kappa light chains, and for which high resolution (<3.5 Å) experimental structures were available (**Table 2**). An equal number of design specifications were allocated to each framework.

**Table 2.** Summary of frameworks (FWRs) used as inputs for Origin-1 *in vitro* performance evaluation. PDB = Protein Data Bank.

| Framework | PDB ID | Heavy Chain Germline | Light Chain Germline |
| --- | --- | --- | --- |
| Trastuzumab | 1N8Z | IGHV3-66 | IGKV1-39 |
| Relatlimab | 7UM3 | IGHV4-34 | IGKV3-11 |
| Dupilumab | 6WG8 | IGHV3-23 | IGKV2D-28 |
| Sotrovimab | 6WS6 | IGHV1-18 | IGKV3-20 |

##### CDR Length Determination

HCDR1, HCDR2, and LCDR2 lengths were fixed to the germline CDR length, as determined by the selected FWRs. HCDR3, LCDR1, and LCDR3 lengths were sampled independently of one another from the empirical distributions observed within AbData (Supplementary Table 5). The distribution was truncated to prevent overly short or overly long CDRs. We used the Kabat [51] definition of LCDR2 and the IMGT [52] definition of HCDR1, HCDR2, HCDR3, LCDR1, and LCDR3. We used Kabat for LCDR2 to extend the typically very short (3 amino acid) LCDR2 by IMGT notation.

##### Design Specifications

For each target, 3360 “design specifications” (configurations of target, epitope subsample, FWR, and CDR lengths) were sampled to provide diverse model input feature sets within the constraints of the design problem.

#### 2.4.3 Antibody library design

Given a set of sampled input specifications, Origin-1 was applied through a series of *in silico* search and evaluation steps to iteratively design and score candidate binders to the intended targets at the desired epitopes. These stages comprised Wide Structure Search, Deep Structure Search, and Sequence Search (**Supplementary Figure 16**). Different metrics were used to evaluate designs at each of these stages depending on the type of output being assessed (**Supplementary Table 6**). The motivation for spreading design and GPU compute allocation across three stages was to concentrate resources on the most promising candidates in a particular search effort. During Wide Structure Search, backbone structures were designed for a large number of design specifications, and design specifications corresponding to the top structures were selected. During Deep Structure Search, additional structures were generated based on these design specifications, and top structures were selected. During Sequence Search, additional sequences were designed for the top structures, culminating in selecting the top sequence for each of the top 95 structures. This scaling approach enabled stepwise improvements in AbsciBind Score among top designs with each search stage (**Supplementary Figure 17**). We describe the strategies underlying Wide Structure Search, Deep Structure Search, and Sequence Search in Supplement §7.6.1 and include detailed library design methodology in Supplement §7.6.

#### 2.4.4 Lead Optimization

We optimized the top designs against all four targets, as well as a second design against AZGP1, by modifying Hie et al.’s Efficient Evolution strategy [53, 54] to generate a library of single-mutant variants relative to the parent designs for a first round of optimization. Specifically, we integrated the AbsciBind protocol into this approach and used AbsciBind Score to support selection of variants for downstream experiments. A collection of protein language models (ESM-1b [55], ESM-1v [56], ESM2-650M [57], and AbLang2 [58]) was used in addition to the AbsciBind Score. Data generated from this first round were used to create a library of higher-order mutants for a second round of optimization. Scoring, selection, and optimized variant library creation are described in Supplement §7.7.

#### 2.4.5 *In vitro* Assessment of Computational Designs

*In vitro* experimental methods used to assess binding, developability, structural fidelity, and function are reported in Supplement §7.8.

## 3 Results

### 3.1 *In silico* Benchmarking of Origin-1 Reveals Superior Antibody-Antigen Complex Design Relative to Field Standard

Results from AbsciGen vs. RFantibody benchmarking experiments revealed that AbsciGen outperforms RFantibody when designing antibody sequences and structures for targets without known binders according to *in silico* unsupervised metrics (**Table 3**, **Figure 7**). Using the AbsciBind Score, internally validated to have high discrimination power for binding prediction, AbsciGen produces 23.50% of designs with AbsciBind Score ≥ 0.5, compared to 0.80% for RFantibody, constituting a nearly 30-fold improvement. AbsciGen also achieves a higher mean AbsciBind Score than RFantibody, further demonstrating its superior performance for antibody-antigen binding prediction. Furthermore, AbsciBind Score distributions across ten targets showed that AbsciGen frequently generates designs with significantly higher scores, indicating improved overall quality. Additional results from AbsciGen benchmarking experiments, including per-target statistics and a comparison between the original RFantibody and our modified version, can be found in §7.4.

**Figure 7.**
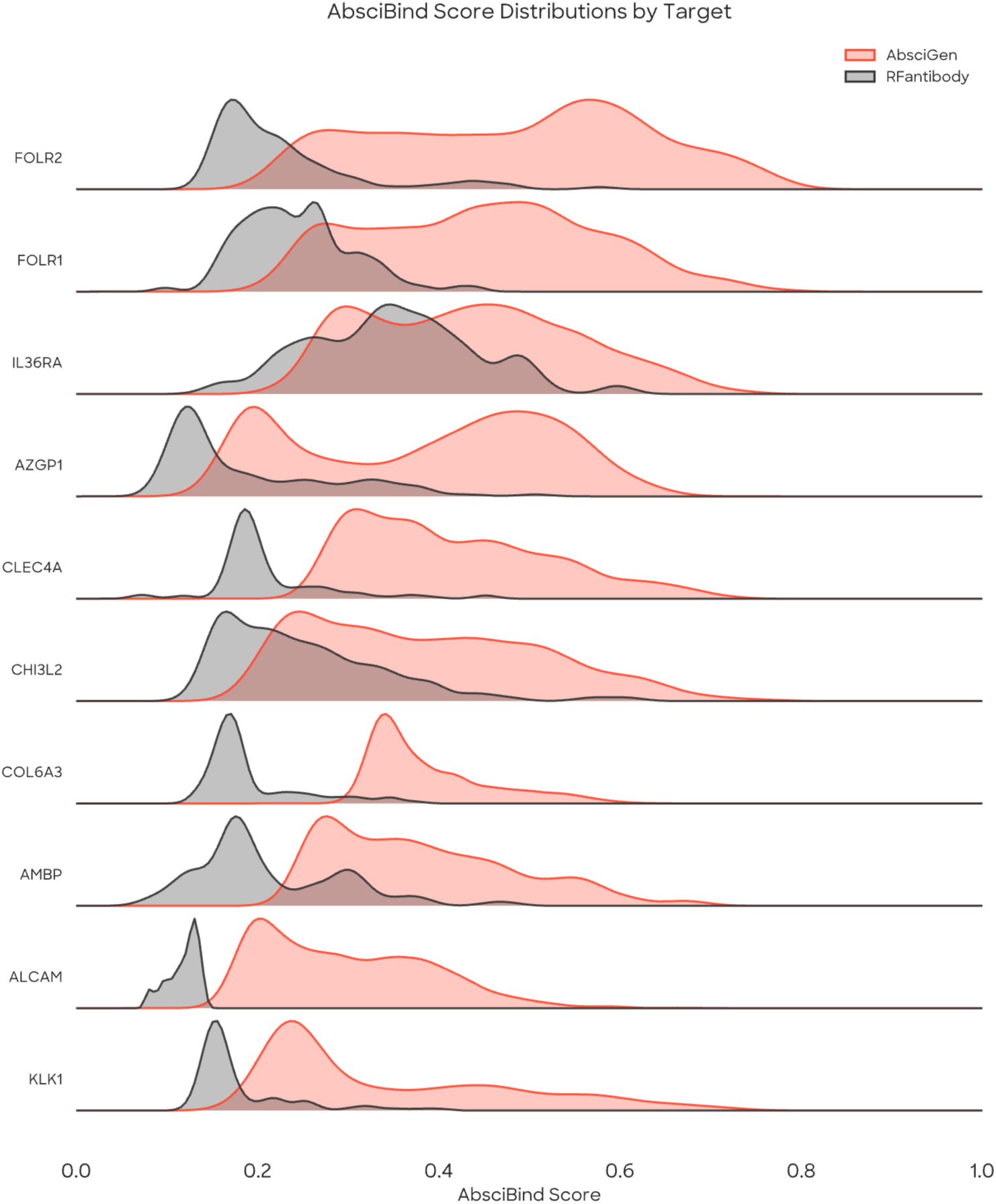
Distribution of AbsciBind Scores for AbsciGen vs. RFantibody across ten targets. AbsciGen generates designs with higher average AbsciBind Scores than RFantibody for all targets assessed.

**Table 3.** Summary of results from AbsciGen vs. RFantibody benchmarking experiments. The AbsciBind Score is computed as the average of ipTM and Antibody-Aligned ipTM scores from the AbsciBind protocol. Metrics are computed across all targets assessed. Results are reported as mean +/- standard deviation. * indicates *p* < 0.001 by Mann-Whitney U. Best overall is marked in bold.

| Method | Mean AbsciBind Score | % Designs with AbsciBind Score $\geq 0.5$ |
| --- | --- | --- |
| RFantibody | 0.218 ± 0.092 | 0.80% |
| AbsciGen | **0.398 ± 0.129\*** | **23.50%** |

OASis score distributions for sequences generated by AbsciGen vs. RFantibody for all targets suggest that AbsciGen achieves significantly higher humanness scores than RFantibody (**Figure 8**). We suspect this is due to IgDesign2’s fine-tuning on antibody-antigen complexes, which ProteinMPNN lacks.

**Figure 8.**
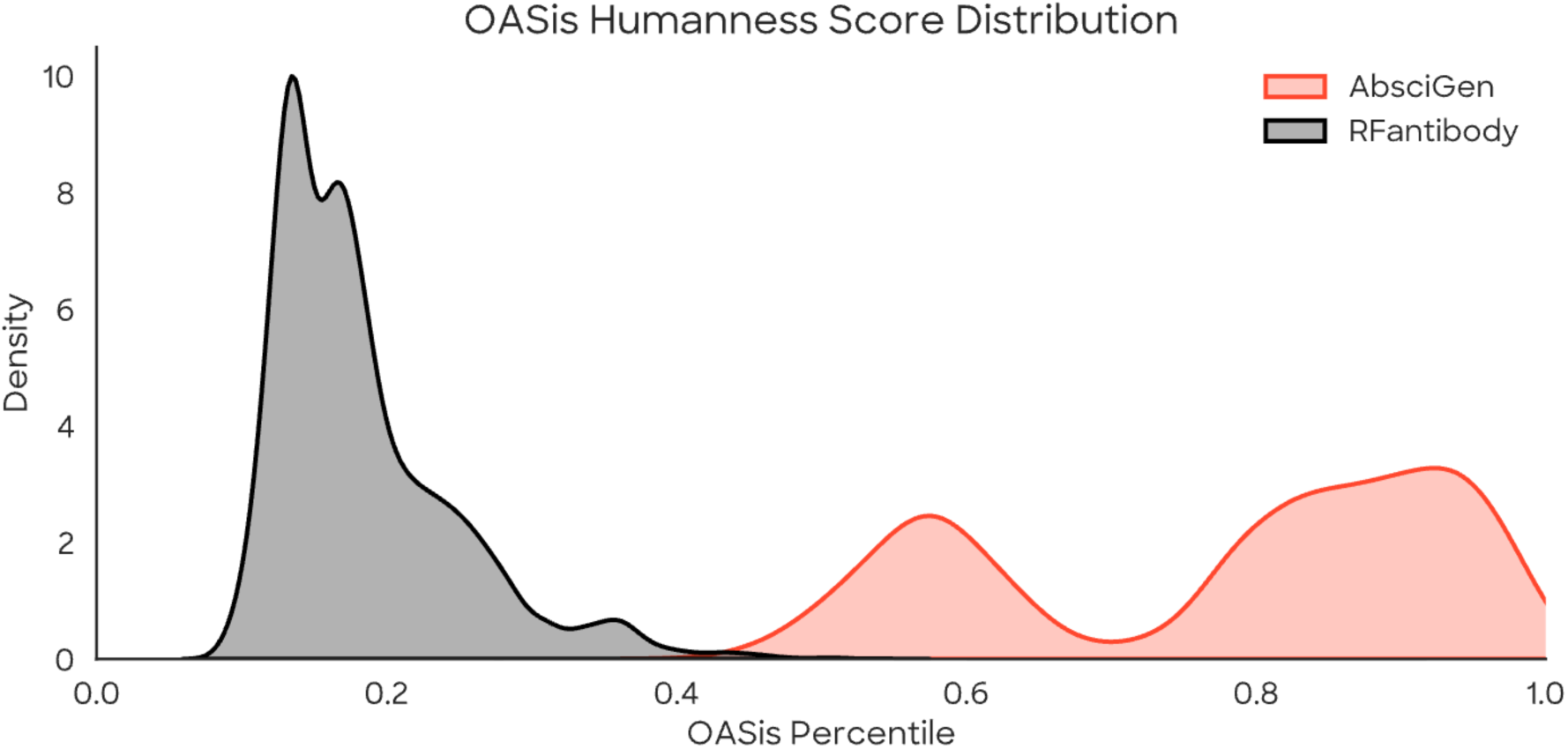
Distribution of OASis percentile scores for sequences generated by both models across four targets. Higher percentiles suggest sequences are more likely to exhibit human-like antibody qualities.

### 3.2 SPR Identifies Multiple Hits

SPR identified three Origin-1 hits against COL6A3, four against AZGP1, one against CHI3L2, and one against IL36RA (**Figures 9-12**). No hits were identified against FOLR1, FOLR2, CLEC4A, AMBP, ALCAM, or KLK1. As noted in the Methods, we screened all designs against two non-antigen commercial protein targets (TL1A and PRLR) to assess polyspecificity, and all antigens against unintended designs to assess target stickiness. All of the aforementioned hits demonstrated binding to their designed antigen, did not bind to unintended designs, and did not bind to the off-target proteins based on binding specificity criteria described in Supplement §7.8.2. This result inspires confidence that Origin-1 can *de novo* design antibody binders to novel epitopes. Additional binding results and measured binding affinities are reported in Supplement §7.8.2.

**Figure 9.**
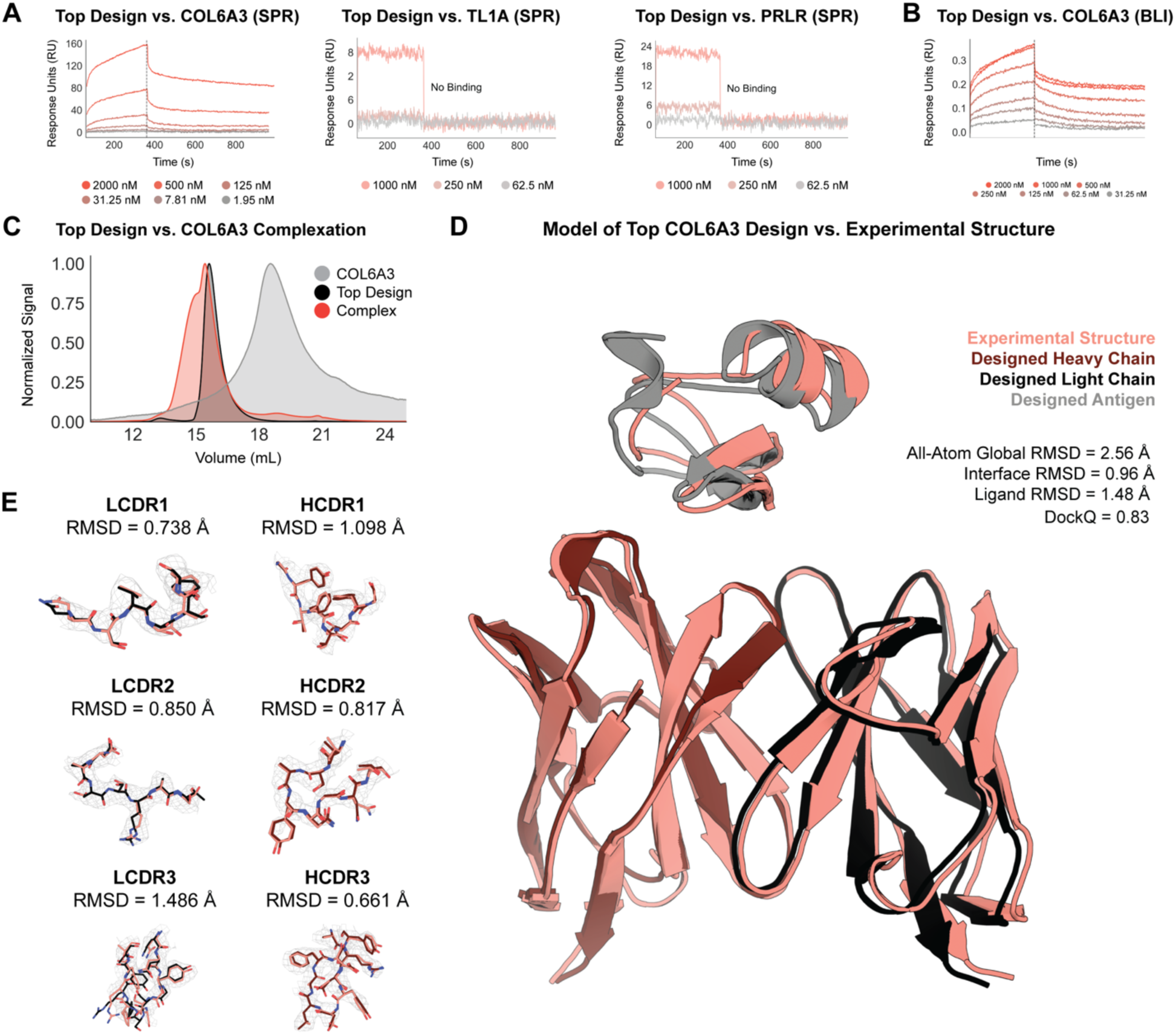
*In vitro* experimentation confirms that Origin-1 generated a *de novo* binder against a novel epitope on COL6A3. (A) SPR demonstrates that the top design binds to COL6A3 in mAb format and does not bind to either of two unintended targets (TL1A and PRLR) (B) BLI confirms that the top design binds to COL6A3 in Fab format. (C) Complexation experiment confirms that the top design, in Fab format, binds to COL6A3 in solution. (D) Cryo-EM of top design complexed with COL6A3 confirms epitope-specificity and atomic accuracy of the Origin-1 computational model. (E) CDR-specific analysis of model vs. solved complex structure confirms atomic accuracy.

**Figure 10.**
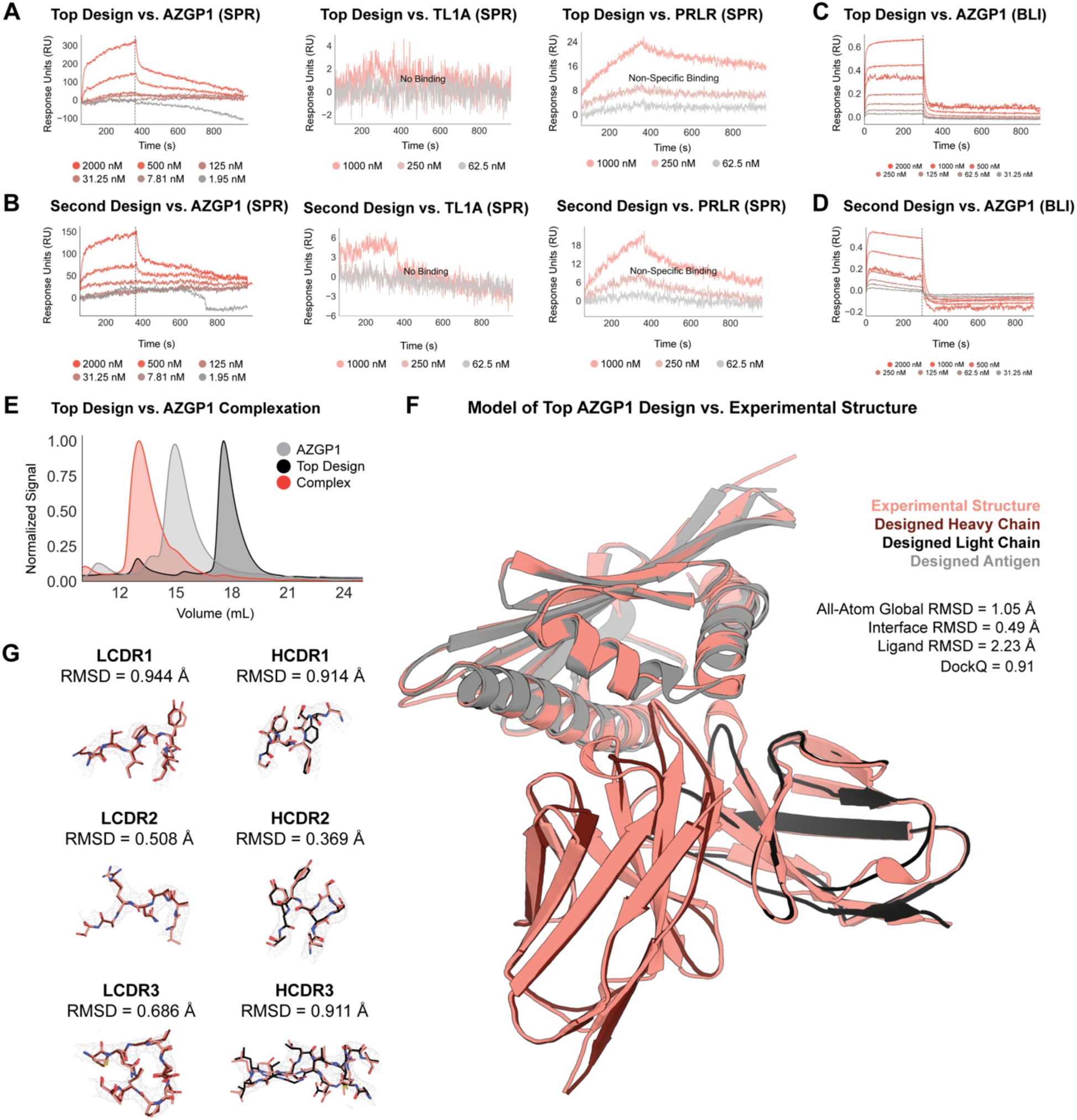
*In vitro* experimentation confirms that Origin-1 generated a *de novo* binder against a novel epitope on AZGP1. (A) SPR demonstrates that the top design binds to AZGP1 in mAb format and does not bind specifically to either of two unintended targets (TL1A and PRLR). (B) BLI confirms that the top design binds to AZGP1 in Fab format. (C) SPR demonstrates that the second design binds to AZGP1 in mAb format and does not bind specifically to either of two unintended targets (TL1A and PRLR). (D) BLI confirms that the second design binds to AZGP1 in Fab format. (E) Complexation experiment confirms that the top design, in Fab format, binds to AZGP1 in solution. (F) Cryo-EM of top design complexed with AZGP1 confirms epitope specificity and atomic accuracy of the Origin-1 computational model. (G) CDR-specific analysis of model vs. solved complex structure confirms atomic accuracy.

**Figure 11.**
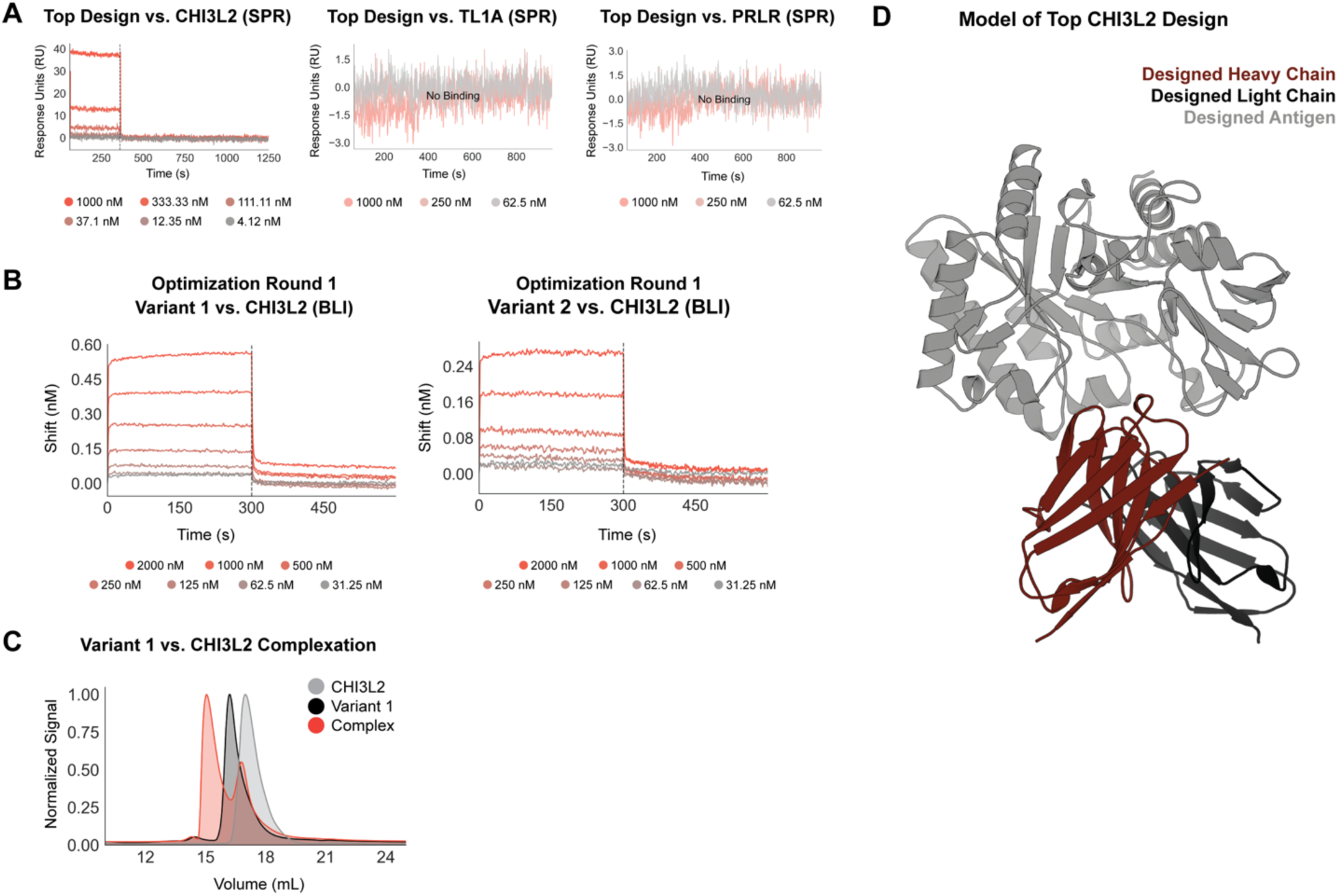
*In vitro* experimentation confirms that Origin-1 generated a *de novo* binder against a novel epitope on CHI3L2. (A) SPR demonstrates that the top design binds to CHI3L2 in mAb format and does not bind to either of two unintended targets (TL1A and PRLR). (B) BLI confirms that the two optimized variants bind to CHI3L2 in Fab format. (C) Complexation experiment confirms that optimized variant 1, in Fab format, binds to CHI3L2 in solution. (D) Origin-1 computational model of top design in complex with CHI3L2 is shown.

**Figure 12.**
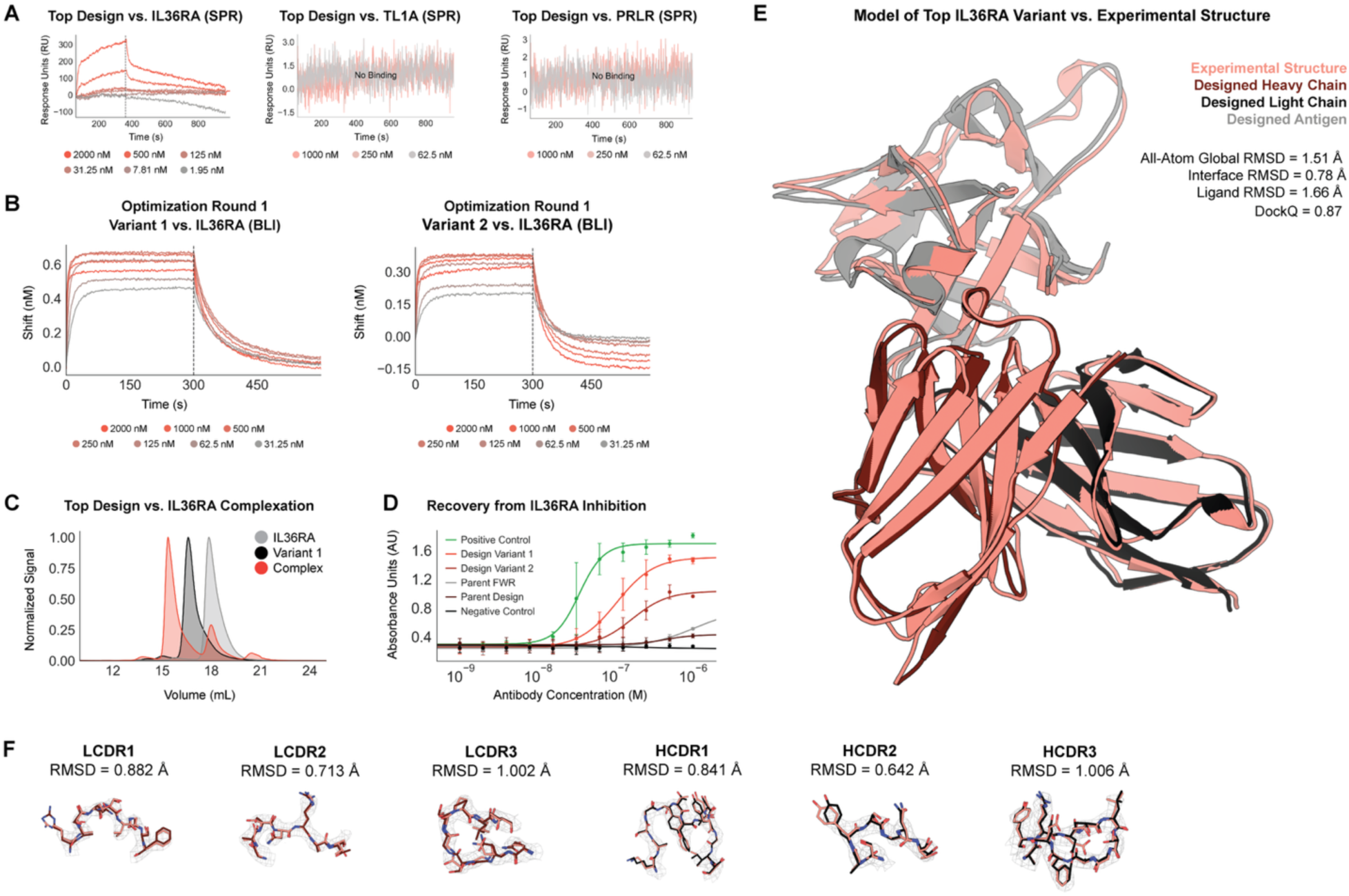
*In vitro* experimentation confirms that Origin-1 generated a *de novo* binder against a novel epitope on IL36RA. (A) SPR demonstrates that the top design binds to IL36RA in mAb format and does not bind to either of two unintended targets (TL1A and PRLR). (B) BLI confirms that the two optimized variants bind to IL36RA. (C) Complexation experiment confirms that optimized variant 1, in Fab format, binds to IL36RA in solution. (D) HEKBlue functional assays demonstrate that the top two optimized variants antagonize IL36RA-mediated inhibition. (E) Cryo-EM of top variant from first round of lead optimization complexed with IL36RA confirms epitope specificity and atomic accuracy of the Origin-1 computational model. (F) CDR-specific analysis of model vs. solved complex structure confirms atomic accuracy.

### 3.3 BLI and Complexation via aSEC Confirm Binding

To further validate hits identified via SPR, we performed both on-target and off-target BLI experiments in mAb format. Hits in mAb format were assessed against their intended target; BLI confirmed one hit against COL6A3 (**Figure 9**), and two hits against AZGP1 (**Figure 10**). Limited mAb BLI binding was observed for the hits against CHI3L2 and IL36RA (data not shown). All hits lacked binding against a negative control antigen.

In preparation for downstream complexation experiments, BLI-confirmed hits were reformatted as Fabs and were re-assessed for binding in BLI. Both AZGP1 hits were confirmed to bind in both assay orientations (**Figure 10**, antigen-immobilized data not shown). The COL6A3 hit bound in the original orientation of the assay (antibody-immobilized, **Figure 9**), but failed to bind in the flipped orientation (antigen-immobilized, data not shown). We hypothesize that this hit bound in only one orientation due to the small size of the antigen and potential tag interference impacting the binding interface when the antigen was immobilized on the probe.

The most promising hits against COL6A3 and AZGP1 were then complexed with their respective targets in Fab format. Complexes were successfully formed for both design-target pairs, confirming binding (**Figures 9** and **10**; Supplement §7.8.4).

### 3.4 Optimization of Zero-Shot Designs Produces Sub-Nanomolar Binders and Confirms Origin-1’s Success in Binder Identification

As mentioned above, SPR-identified hits for CHI3L2 and IL36RA could not subsequently be confirmed by BLI. We reasoned the lack of BLI confirmation was due to low affinity (K_D_ > 2 µM) and chose not to fully characterize them due to the large amount of antigen that would be required.

Instead, we used our lead optimization platform to affinity-mature not only the SPR-identified hits against CHI3L2 and IL36RA, but also the hits against COL6A3 and AZGP1 that were confirmed in BLI. For each of these hits, we designed a first round of optimization libraries composed of single-mutants relative to the parental hit sequences. This approach resulted in affinity improvements across all hits, with the greatest improvement (68-fold) observed for IL36RA (**Figure 12**). We report binding affinities for top designs (referred to as parent designs) and optimized variants in Supplementary Figure 18.

We repeated SPR and BLI experiments using the top affinity-matured designs against CHI3L2 and IL36RA, which demonstrated approximately 4X and 68X increases in affinity relative to their parent designs, respectively. We also validated binding by complexing CHI3L2 and IL36RA with their respective top affinity-matured design variants (**Figures 11** and **12**; Supplement §7.8.4). This result confirms that Origin-1 was able to generate binders against CHI3L2 and IL36RA and further demonstrates the utility of our platform to identify and rescue even very weak binders.

For IL36RA and COL6A3, we designed a second round of optimization libraries by combining beneficial single-mutants into higher-order mutants as described in §7.7. These libraries produced fourteen (14) sub-nanomolar binders against IL36RA, several of which are highlighted in **Figure 13**, and three (3) sub-nanomolar binders against COL6A3 (**Figure 14**). Additionally, we confirmed that the binders designed against human IL36RA bound both human and mouse IL36RA and that the optimization process produced single digit nanomolar binders against mIL36RA. We hypothesize that this is a result of selecting an epitope with substantial homology (>90%) across the orthologs, highlighting an advantage of combining *de novo* design with rational epitope selection.

**Figure 13.**
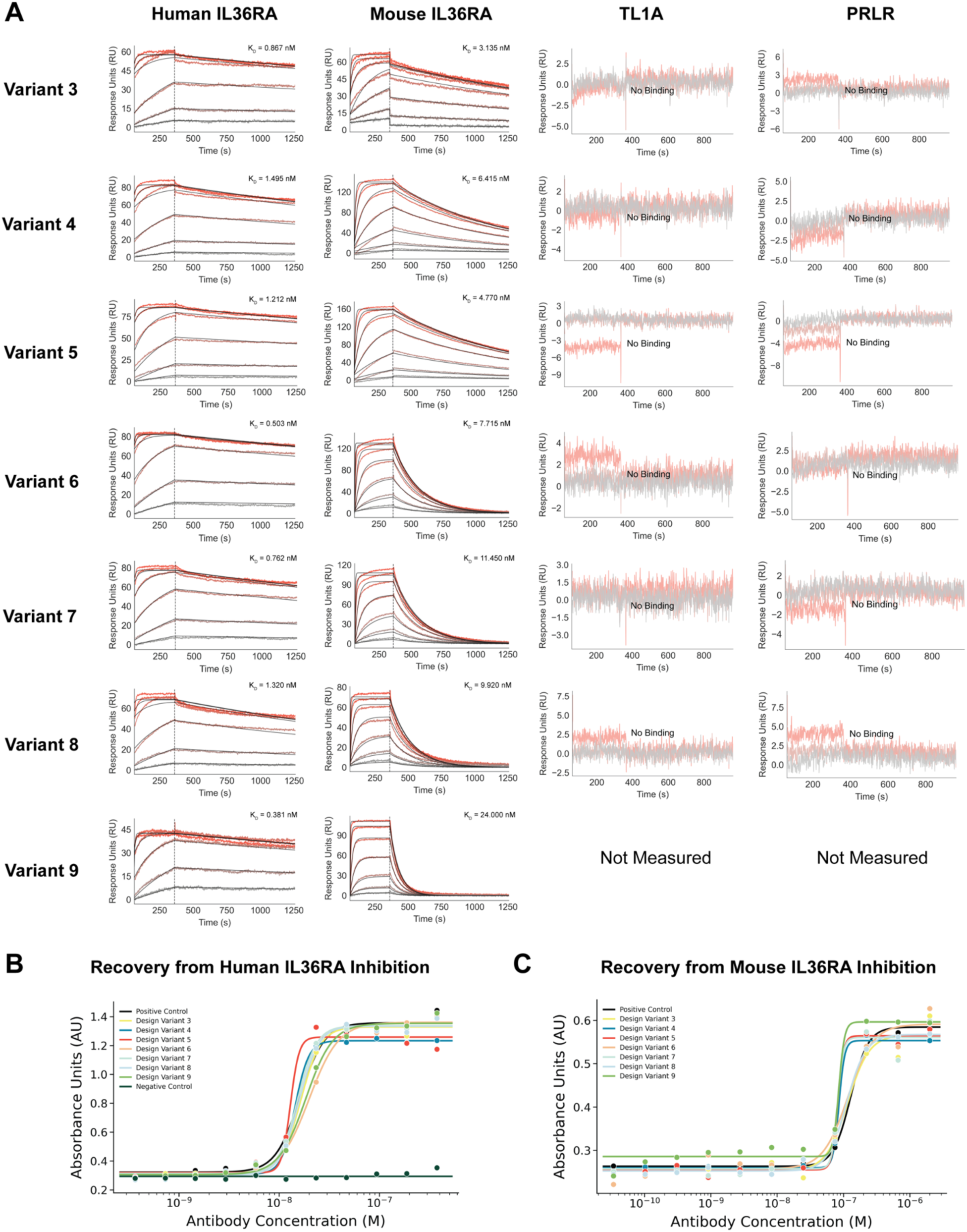
Lead optimization produced sub-nanomolar, cross-reactive binders with high potency against IL36RA. (A) Sensorgrams and affinities of selected designs against human IL36RA, mouse IL36RA, and two off-targets (TL1A and PRLR). Designs are high-affinity and cross-reactive against human and mouse IL36RA while also being specific. (B) HEKBlue functional assay confirms high potency against human cells with top EC50 of 12.3 nM. (C) CT-26 CCL2 functional assay confirms functional antagonism against mouse cells.

**Figure 14.**
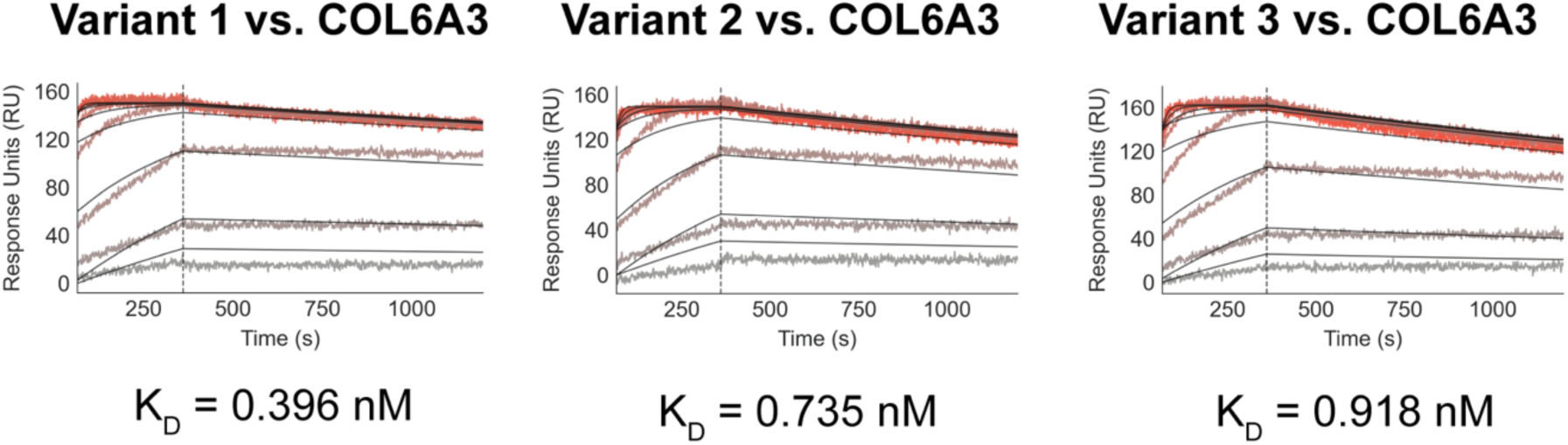
Sensorgrams and affinities for sub-nanomolar binders against COL6A3 that were produced by lead optimization.

The full set of binding affinities can be found [here](https://github.com/AbSciBio/origin-1/tree/main).

### 3.5 Developability Assessments

All zero-shot Origin-1 binders were evaluated across a panel of developability and material quality properties, including polyreactivity, polydispersity, self-association, hydrophobicity, melting temperature, and purity. COL6A3, CHI3L2, and IL36RA binders met therapeutically acceptable criteria for these developability properties, aside from one hydrophobicity flag associated with the IL36RA binder. For AZGP1, our top binder was flagged for self-association, hydrophobicity, and polyreactivity (**Table 4**). A subset of six variants from the second round of IL36RA lead optimization were selected for developability assessment (**Table 5**). Most variants had comparable developability to their parent *de novo* binder and we did not observe substantial increases in polyreactivity despite increasing affinity by >1000-fold. Methods for developability assessments are reported in Supplement §7.8.5.

**Table 4.** Summary of developability results from top binders across COL6A3, AZGP1, IL36RA, and CHI3L2 as well as controls. Trastuzumab (negative), Cetuximab (negative), Briakinumab (positive), and Bococizumab (positive) were used as controls for polyreactivity. Trastuzumab (negative), Briakinumab (positive), and Infliximab (positive) were used as controls for AC-SINS. Trastuzumab was used a control for HIC, DSF, and DLS. RNS = Range Normalized Sum; HIC = Hydrophobic Interaction Chromatography, RRT = Relative Retention Time (relative to Trastuzumab); AC-SINS = Affinity-Capture Self-Interaction Nanoparticle Spectroscopy; DSF = Differential Scanning Fluorimetry; DLS = Dynamic Light Scattering; PDI = Polydispersity Index; N.D. = Not Detected. N.M. = Not Measured.

| Target | Polyreactivity<br>(Anti-Insulin,<br>RNS) | Polyreactivity<br>(Anti-DNA,<br>RNS) | HIC<br>(RRT) | AC-SINS<br>(nM) | DSF,<br>$T_{m1}$<br>(°C) | DLS<br>(Cumulant<br>PDI) |
| --- | --- | --- | --- | --- | --- | --- |
| COL6A3 | 3.5 | 3.8 | 1.03 | 1.6 | 68.00 | 0.00 |
| AZGP1 | 15.8 | 8.5 | N.D. | 25.7 | 77.86 | 0.08 |
| IL36RA | 3.0 | 3.6 | 1.19 | 0.6 | 70.60 | 0.00 |
| CHI3L2 | 3.6 | 4.0 | 0.91 | -0.1 | 70.20 | 0.01 |
| **Control** |  |  |  |  |  |  |
| Trastuzumab | 4.0 | 4.1 | 1.00 | 1.6 | 71.1 | 0.01 |
| Cetuximab | 5.5 | 4.6 | N.M. | N.M. | N.M. | N.M. |
| Briakinumab | 39.9 | 35.5 | N.M. | 28.9 | N.M. | N.M. |
| Bococizumab | 43.6 | 36.9 | N.M. | N.M. | N.M. | N.M. |
| Infliximab | N.M. | N.M. | N.M. | 31.2 | N.M. | N.M. |

**Table 5.**
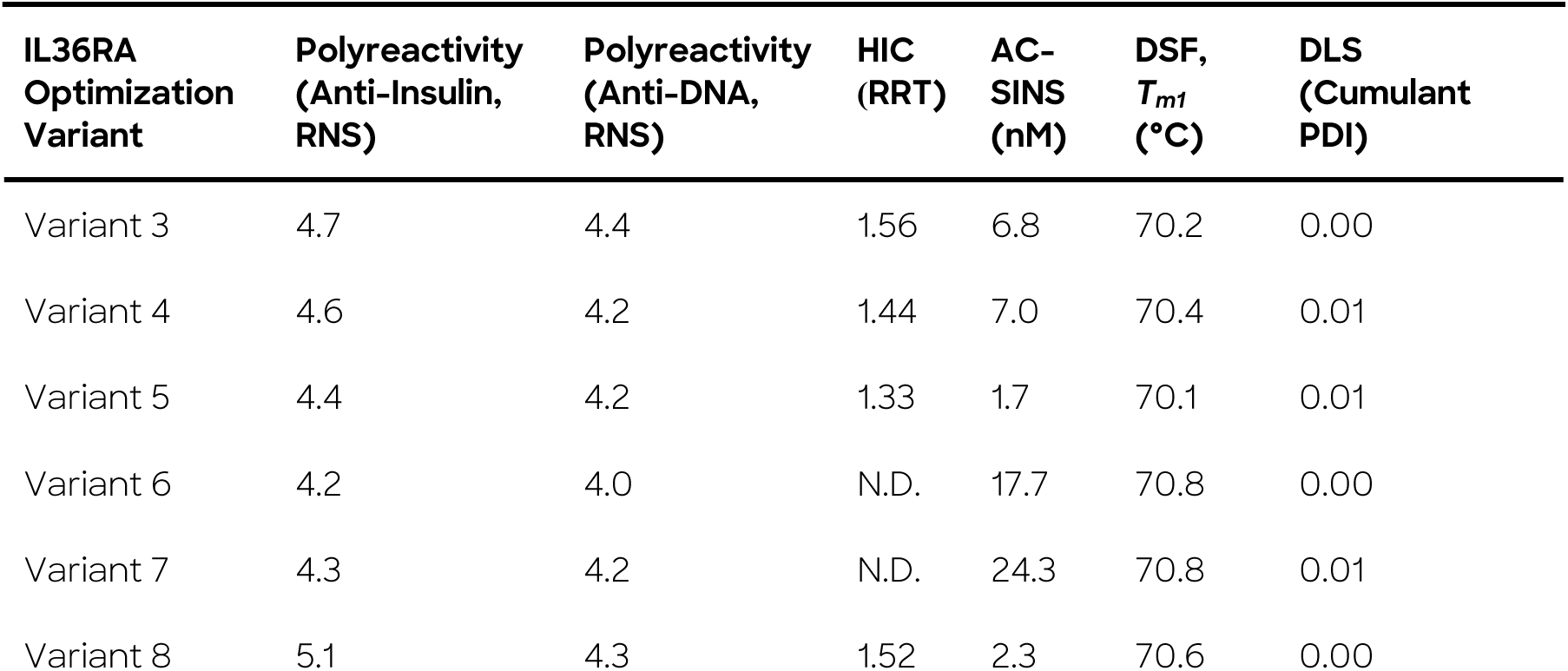
Summary of developability results from top binders against IL36RA from second round of lead optimization. Results for control antibodies are reported in **Table 4**. R2 = Round 2 Lead Optimization; RNS = Range Normalized Sum; HIC = Hydrophobic Interaction Chromatography, RRT = Relative Retention Time (relative to Trastuzumab); AC-SINS = Affinity-Capture Self-Interaction Nanoparticle Spectroscopy; DSF = Differential Scanning Fluorimetry; DLS = Dynamic Light Scattering; PDI = Polydispersity Index; N.D. = Not Detected. N.M. = Not Measured.

| IL36RA<br>Optimization<br>Variant | Polyreactivity<br>(Anti-Insulin,<br>RNS) | Polyreactivity<br>(Anti-DNA,<br>RNS) | HIC<br>(RRT) | AC-SINS<br>(nM) | DSF,<br>$T_{m1}$<br>(°C) | DLS<br>(Cumulant<br>PDI) |
| --- | --- | --- | --- | --- | --- | --- |
| Variant 3 | 4.7 | 4.4 | 1.56 | 6.8 | 70.2 | 0.00 |
| Variant 4 | 4.6 | 4.2 | 1.44 | 7.0 | 70.4 | 0.01 |
| Variant 5 | 4.4 | 4.2 | 1.33 | 1.7 | 70.1 | 0.01 |
| Variant 6 | 4.2 | 4.0 | N.D. | 17.7 | 70.8 | 0.00 |
| Variant 7 | 4.3 | 4.2 | N.D. | 24.3 | 70.8 | 0.01 |
| Variant 8 | 5.1 | 4.3 | 1.52 | 2.3 | 70.6 | 0.00 |

### 3.6 Cryo-EM Confirms Atomic Accuracy of Designs Against COL6A3, AZGP1, and IL36RA

To assess structural fidelity, we used cryo-EM to solve the structures of the top designs against COL6A3 and AZGP1 as well as the top first round lead optimization variant against IL36RA. The experimental structures were resolved with 3.0, 3.3, and 3.1 Å resolution, respectively, and show high concordance with the designed structures generated by Origin-1, confirming both epitope-specificity and atomic accuracy (**Figures 9, 10**, and **12**).

The computational model of the COL6A3 design complex and the corresponding experimental complex structure have an all-atom global RMSD of 2.56 Å, interface RMSD of 0.96 Å, ligand RMSD of 1.48 Å, and a DockQ [59] of 0.83. When overlaid, the CDRs have all-atom RMSDs of 0.738 Å, 0.850 Å, 1.486 Å, 1.098 Å, 0.817 Å, and 0.661 Å for LCDR1, LCDR2, LCDR3, HCDR1, HCDR2, and HCDR3, respectively.

The computational model of the AZGP1 design complex and the corresponding experimental complex structure have an all-atom global RMSD of 1.05 Å, interface RMSD of 0.49 Å, ligand RMSD of 2.23 Å, and a DockQ of 0.91. When overlaid, the CDRs have all-atom RMSDs of 0.944 Å, 0.508 Å, 0.686 Å, 0.914 Å, 0.369 Å, and 0.911 Å for LCDR1, LCDR2, LCDR3, HCDR1, HCDR2, and HCDR3, respectively.

The computational model of the IL36RA design complex and the corresponding experimental complex structure have an all-atom global RMSD of 1.51 Å, interface RMSD of 0.78 Å, ligand RMSD of 1.66 Å, and a DockQ of 0.87. When overlaid, the CDRs have all-atom RMSDs of 0.882 Å, 0.713 Å, 1.002 Å, 0.841 Å, 0.642 Å, and 1.006 Å for LCDR1, LCDR2, LCDR3, HCDR1, HCDR2, and HCDR3, respectively.

### 3.7 Optimized IL36RA Designs are Functional Antagonists

The top two affinity-matured variants from the first round of lead optimization along with the parent IL36RA design were assessed for function in a HEKBlue assay (Supplement §7.8.7). While the parent was not functional, the affinity-matured variants displayed antagonism that correlated with soluble protein affinity (**Figure 12**), suggesting a clear path to a more potent molecule. The highest affinity variant achieved an EC_50_ of 104 nM.

Indeed, by combining single mutants into a library of higher-order mutants for a second round of lead optimization, we identified multiple sub-nanomolar affinity binders. A subset of these were tested again in the HEKBlue assay and the highest potency achieved was an EC_50_ of 12.3 nM (**Figure 13**). After confirming cross-reactivity of our designs with mouse IL36RA, we also tested for and confirmed functional antagonism on mouse cells using an internally developed CT-26 + CCL2 assay (**Figure 13**, Supplement §7.8.7).

## 4 Discussion

Here we develop and experimentally validate Origin-1, an AI platform for *de novo* antibody design against “zero-prior” epitopes. We describe the methodology underlying Origin-1’s component AI models: AbsciDiff and IgDesign2 (together, AbsciGen) for generative structure and sequence design, respectively, and AbsciBind for design scoring and selection for downstream experimental validation. Our *in silico* benchmarking of AbsciGen and AbsciBind confirms that the respective strategies compete with and outperform existing protocols for antibody-antigen complex design and evaluation (**Table 3**, **Figures 7** and **8**), and our *in vitro* assessments confirm that in fewer than one hundred attempts, Origin-1 can generate designs that bind with specificity and are developable (**Figures 9-12**, **Table 4**). We show that our modeled antibody designs are consistent with their experimental structures with atomic accuracy (**Figures 9**, **10**, and **12**), and we demonstrate our ability to use AbsciBind Scores to generate optimized variants of our zero-shot designs that achieve sub-nanomolar affinity and are functional antagonists in both human and mouse cellular environments (**Figures 12** and **13**). Together, these results motivate further investigation into Origin-1’s potential to generate antibody therapeutic candidates against novel disease targets.

One limitation of the current effort is that the hit rates obtained by Origin-1 in a zero-shot manner were lower (at most four from approximately 100 designs per target) than hit rates that have recently been reported by others in the field [23–27]. In interpreting this, one should consider both the design challenge undertaken and the extent of experimental validation conducted to support a reported hit rate. Here, to our knowledge, we extend the complexity of our design challenge beyond what others have reported by designing antibodies against “zero-prior” epitopes for which there are no publicly available complex structures that directly provide epitope residue inputs to guide antibody design. In other words, Origin-1 had to identify antigen residues that were viable candidates for antibody binding and sample correctly from these residues to guide design toward these regions, in addition to correctly scoring designs and selecting winning candidates from among those proposed. We increased the task complexity in this way to test Origin-1 in a setting that more closely resembles a challenging drug design effort, where often the epitope itself is unknown and must be determined *de novo*. Beyond increasing the complexity of the design challenge, we also set strict criteria for labeling a design a binder by requiring designs to meet criteria for binding across multiple orthogonal assays as well as confirming designs bind only to their intended targets and that antigens bind only to their intended designs. Our results indicate that Origin-1 is capable of designing antibodies that bind to targets of interest with limited input guiding design localization, and with stringent hit definition criteria, inspiring confidence in the reproducibility, reliability, and translatability of our results. We provide the comprehensive set of *in vitro* data generated in the course of this effort and encourage our colleagues in the field to leverage these datasets to benchmark their own efforts in antibody design.

Future development should prioritize improving AI antibody design protocols such that they directly address the most significant challenges impacting traditional antibody discovery campaigns today: target difficulty, costs, and timelines. Improvements in these areas will ensure that our technology continues to progress toward addressing unmet needs that ultimately benefit patients requiring innovative, cost-effective solutions to treat their ailments.

## 5 Acknowledgements

The authors thank Matthew Saunders, Aayush Arora, Samuel Tsang, Joshua Bennett, Dani Johnson, Namita Varudkar, and Hyun Park for contributing to discussions surrounding the work described here. We thank Simon V. Mathis for contributions to an earlier version of AbData. We thank Dan Rabinovitsj, Mene Pangalos, Daniele Biasci, Wen Sha, and Zach Jonasson for reviewing and providing feedback on earlier versions of this manuscript.

## 7 Supplementary Material

### 7.1 AbData Inclusion Criteria

We applied both temporal and sequence homology filters to AbData-curated complexes to eliminate overlap between our training, validation, and test sets. Temporal filters were applied first, whereby all complexes from PDB entries released before 2024-01-01 were included in training, between 2024-01-01 and 2024-09-30 in validation, and between 2024-09-01 and 2025-07-10 in test. We applied specific sequence homology filters per split, where homology filters defined using the training set were applied to the validation and test set, and filters defined using the validation set were applied to the test set. For antibody-antigen complexes specifically, we removed all entries containing an antigen with greater than 40% sequence similarity to any antigen in the relevant reference set. For protein-protein complexes, we removed all entries for which either chain had greater than 40% sequence similarity to any chain in the relevant reference set.

Finally, we filtered for redundancy by clustering the antigen and Fv sequences using 40% and 100% sequence identity, respectively. Within each cluster-pair, we selected only a single example containing optimal resolution, proportion of resolved residues, and interface contacts. For protein-protein interactions, we used a similar strategy, adjusted by clustering interface chains at 95% sequence identity and selecting within cluster pairs. The final quantities of antibody-antigen entries in the training / validation / test set splits were 3242 / 63 / 84, respectively, covering 2848 / 62 / 84 respective unique HCDR3 sequences. The final quantities of protein-protein complexes were 29835 / 356 / 322, respectively.

We provide a comprehensive list of our antibody-antigen inclusion criteria for AbData below:

- No Fvs without a heavy chain
- No unbound Fvs
- No Fvs bound to short antigens, defined as antigens containing fewer than 15 residues
- No Fvs with small epitopes, defined as epitopes containing fewer than 5 residues
- No Fvs with any missing CDR residues
- No Fvs with buried surface area less than 500 Å^2^ at the antibody-antigen interface
- No Fvs where greater than 50% of CDR contacts occurred with non-protein molecules
- Resolution must be less than 9 Å

Protein-protein interactions were additionally filtered as follows:

- Only dimeric interfaces (i.e. no additional chains at the interface)
- No interfaces with buried surface area < 500 Å^2^
- No complexes with > 2500 residues
- No complexes with < 50 residues

For antibody-antigen validation and test sets, we applied the following filters in addition to the above:

- No Fvs with cysteines in CDRs
- Resolution must be less than 3.5 Å

### 7.2 Structure Design with AbsciDiff

#### Design Region Atom Representation

AbsciDiff enables open-ended generation of amino acid sidechains by representing all design region residues in Atom14 notation, with excess “virtual” atoms placed at each residue’s Cα position (**Supplementary Figure 1**). This superposition technique allows the model to generate structures without committing to specific residue identities during diffusion. The sequence can subsequently be inferred from the generated backbone structure using methods such as IgDesign2.

**Supplementary Figure 1:**
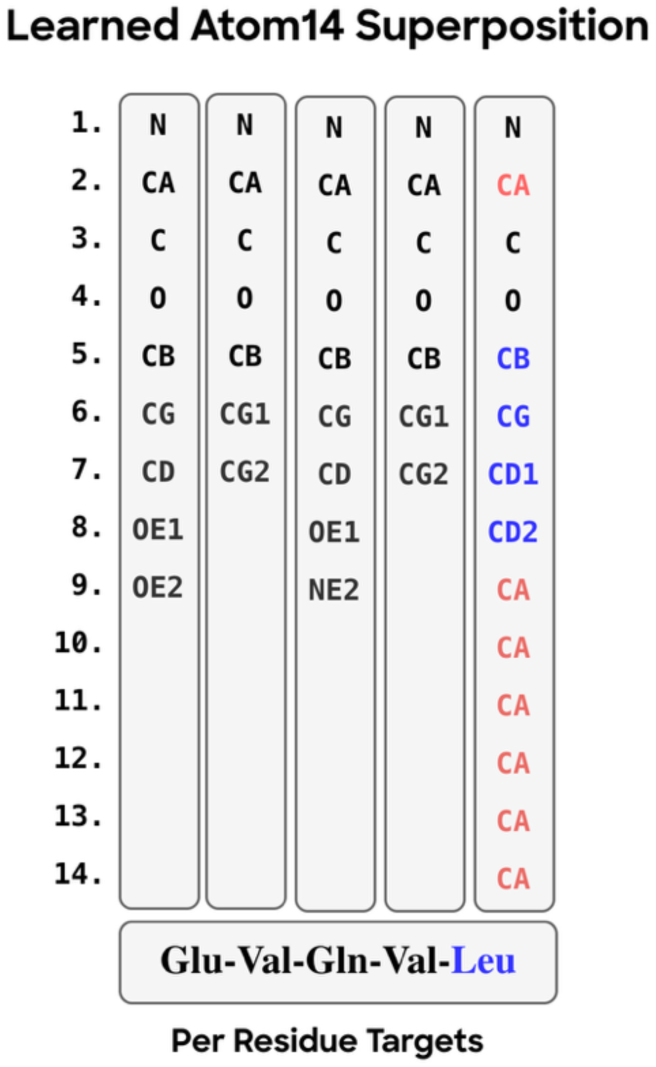
An example of AbsciDiff’s atom superposition technique. Non-design region amino acids are represented with their true atomic structure since their identities are fixed *a priori*. Design region residues (see Leu in blue) use the Atom14 representation, where excess virtual atoms beyond the residue’s actual atom count are placed at the Cα position.

#### Partial Diffusion

In addition to the modifications described in §2.3.1, AbsciDiff’s diffusion module also includes key modifications for partial diffusion. In partial diffusion, sampled datapoints are initialized from a partially noised version of the input rather than pure noise. When enabled, the diffusion process begins at noise level:

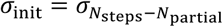

The noised initialization is given by:

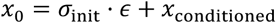

where *ε* ∼ 𝒩(0, *I*), and *x*_conditioned_corresponds to the conditioned coordinates (antigen and framework). This mechanism enables the model to focus the sampling process when full, from-scratch generation is not required, allowing refinement of existing favorable designs or diversification of fixed binding complexes.

### 7.3 Coordinate Initialization for RFantibody Benchmarking

*De novo* antibody design using RFantibody required careful attention to coordinate initialization to achieve reasonable outputs. Without modification, we observed reduced performance compared to the supervised benchmarking setting, including chain breaks in predicted loop structures and glycine-dominated sequences from ProteinMPNN.

**Supplementary Figure 2:**
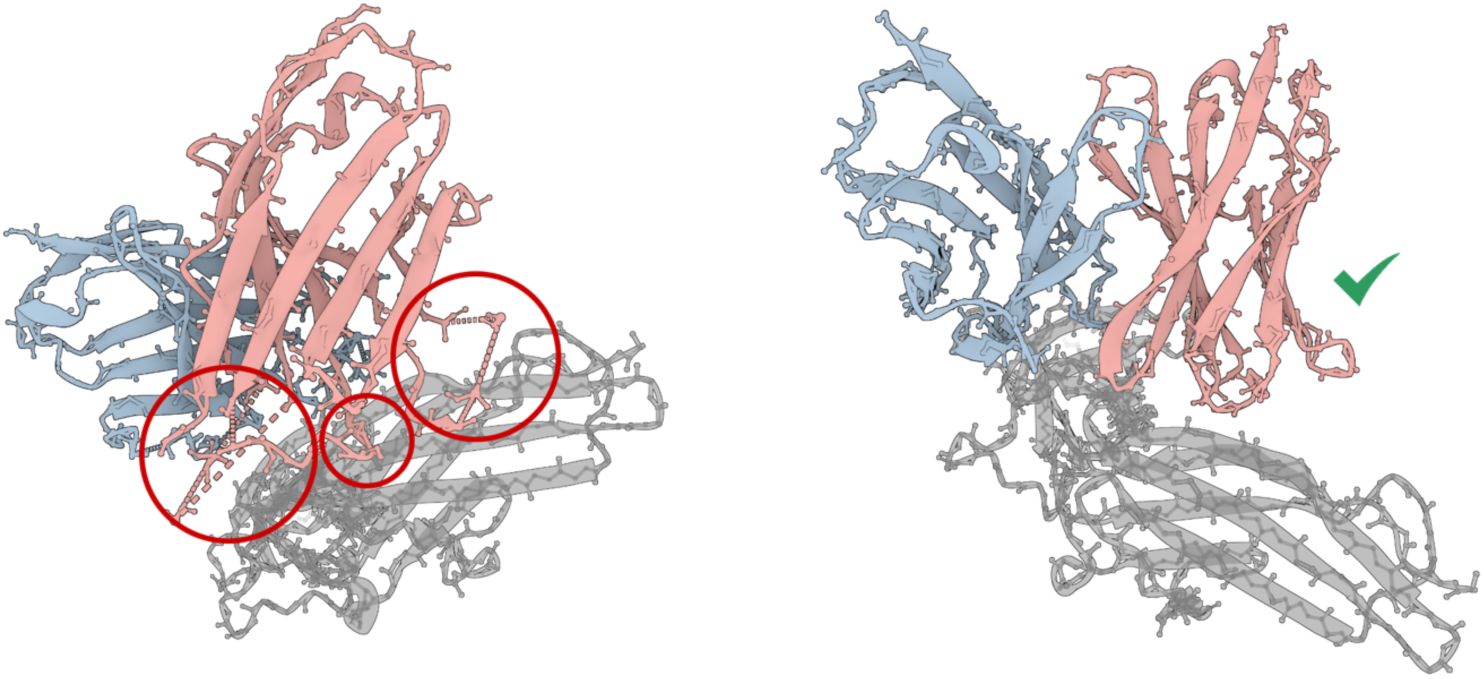
RFantibody loop quality depends on coordinate initialization: chain breaks appear when loop coordinates are placed at the origin (left), but not when idealized coordinates are provided (right). Provided framework PDB files all include ground-truth loop coordinates, motivating our pre-processing changes for a fair *de novo* benchmark.

To avoid potential information leakage under the *de novo* setting, our pipeline preprocesses structures such that CDR loop coordinates are placed at the origin for the designed number of residues, and CDR sequences are populated with glycines. While RFantibody fully masks features for designed loops, we found that initial loop coordinates influence the diffusion process. Specifically, RFantibody’s reverse diffusion implementation during inference parameterizes the initial noise distribution on the input Cα coordinates at timestep *t* = *T*, (rather than sampling from an unconditional prior) causing spatial biases in input coordinates to persist through noising.

By default, when a designed loop length differs from the input “framework” PDB, RFantibody initializes coordinates using idealized backbones with random noise (via “adjust_loop_lengths”). However, when the loop length already matches—as in our pipeline, where we prepare PDBs for each desired length with origin-initialized CDR coordinates — the model uses the provided coordinates directly. What’s more, provided examples in the public RFantibody release include actual loop coordinates from the RCSB PDB entry. As such, to ensure a fair benchmark on examples lacking provided loop coordinates while also limiting changes to the RFantibody codebase, we applied an idealized initialization based on loop “dilation” logic to all CDR coordinates.

Specifically,

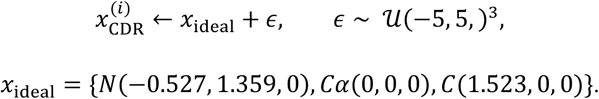

This approach does not fully decouple designs from initialization—loop length and random seed still influence the result—but it respects the intention of the existing loop length dilation methodology without introducing more substantial modifications.

### 7.4 Additional AbsciGen Results

#### 7.4.1 Supervised Task Design and Results

##### Experimental Setup

We evaluate structure and sequence generation against targets with known antibody binders. We do so by taking a wild-type complex, masking CDR and inter-chain distance features, and providing the remaining data features, along with native epitope indices and CDR lengths, to each method. Both methods execute their full pipelines: AbsciGen (AbsciDiff followed by IgDesign2) versus RFantibody (RFdiffusion [9] followed by ProteinMPNN [17]). For sequence design, IgDesign2 autoregressively designs all six CDR sequences conditioning on structural context, while ProteinMPNN uses default settings; both produce eight sequences per structure.

##### Metrics

We use DockQ [59] and antigen-aligned HCDR3 Root Mean Square Deviation (RMSD) to assess structural fidelity, and HCDR3 amino acid recovery (AAR) to assess sequence recovery. While design models are inherently open-ended, we reason that successful models should recover the wild-type structure and sequence given sufficient sampling and training objective.

##### Dataset

We select 44 high-quality antibody-antigen complexes from AbData, balancing target diversity, complex size, and training date cutoff. All complexes are cropped to 512 residues using our spatial cropping method (§2.3.1).

##### Sampling

For each complex, we generate *n* = 100 final structures and *m* = 8 sequences per structure. AbsciDiff produces *M* = 24 structure samples per generation process, which are ranked (§2.3.1) by the confidence module and reduced to *K* = 3 final outputs. For this analysis, we uniformly randomly sample *K* = 1 of these to match the number of designs between pipelines.

#### Supervised Task Results

**Supplementary Table 1** demonstrates that AbsciGen substantially outperforms RFantibody across all metrics. Lower HCDR3 RMSD indicates better structural agreement with the native binding pose, while higher DockQ scores (threshold ≥ 0.23 for acceptable quality) indicate more accurate antibody-antigen interface geometry. For sequence recovery, AbsciGen achieves substantially higher maximum HCDR3 amino acid recovery (AAR) averaged across targets (0.647 vs. 0.371), demonstrating IgDesign2’s effectiveness. We note that sequence recovery is coupled to structure quality - IgDesign2 benefits from higher-fidelity structures produced by AbsciDiff.

**Supplementary Table 1:** Supervised benchmarking results. Metrics are computed for each structure or sequence generation, aggregated per-target using min, max, or median, then averaged across targets. DockQ Success reports the fraction of targets with aggregated DockQ > 0.23. HCDR3 RMSD is computed after aligning on the target structure. * indicates *p* < 0.001 by Mann-Whitney U. Best overall is marked in bold.

|  | HCDR3 RMSD (Å) ↓ |  | DockQ ↑ |  | DockQ Success ↑ |  | HCDR3 AAR ↑ |  |
| --- | --- | --- | --- | --- | --- | --- | --- | --- |
|  | Min | Median | Max | Median | Max | Median | Max | Median |
| **RFantibody** | 6.432 | 13.632 | 0.367 | 0.128 | 77.3% | 6.8% | 0.371 | 0.117 |
| **AbsciGen** | **3.256\*** | **8.166\*** | **0.509\*** | **0.235\*** | **95.5%** | **47.7%** | **0.647\*** | **0.401\*** |

**Supplementary Figure 3** shows per-target results, where AbsciGen achieves acceptable DockQ scores (≥ 0.23) on 95.5% of targets, compared to 77.3% for RFantibody.

**Supplementary Figure 3.**
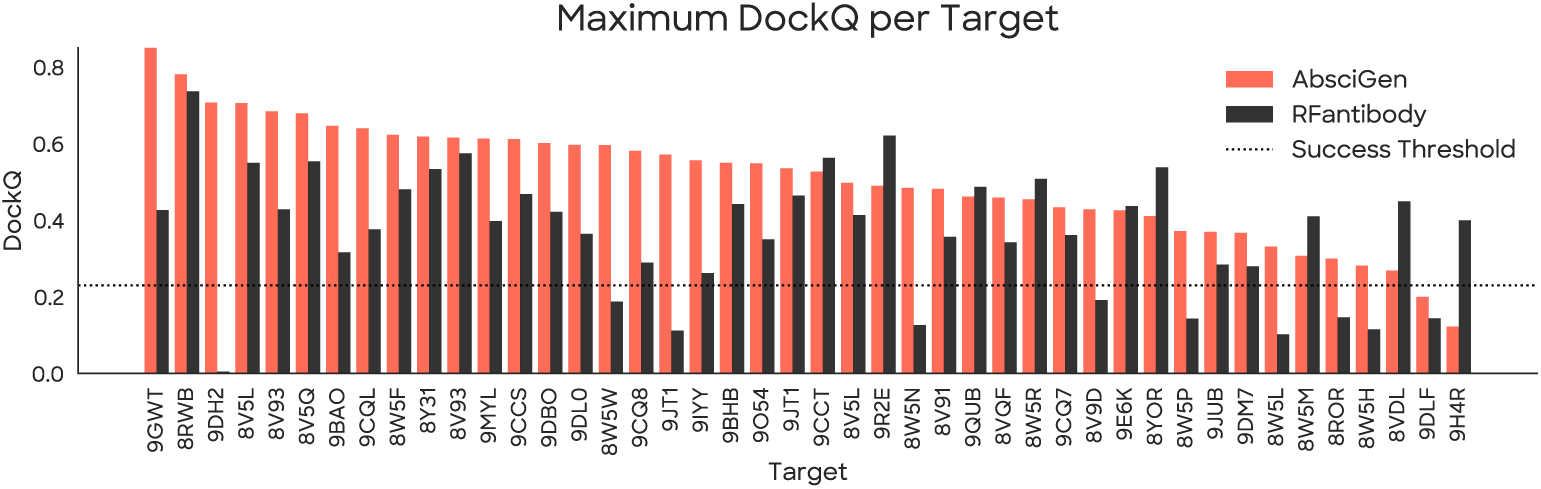
Per-target comparison of maximum DockQ scores for AbsciGen and RFantibody. The acceptable threshold (DockQ = 0.23) is indicated by the dashed line.

**Supplementary Figure 4.**
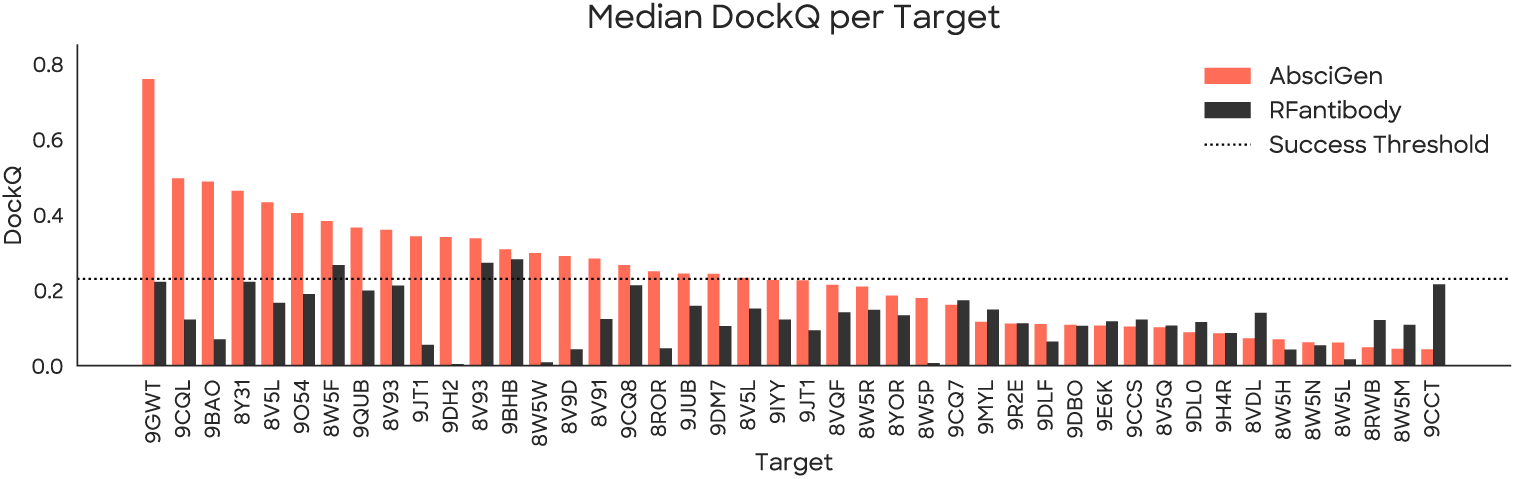
Per-target comparison of median DockQ scores for AbsciGen and RFantibody across all targets in the supervised benchmarking task. The acceptable threshold (DockQ = 0.23) is indicated by the dashed line.

**Supplementary Figure 5.**
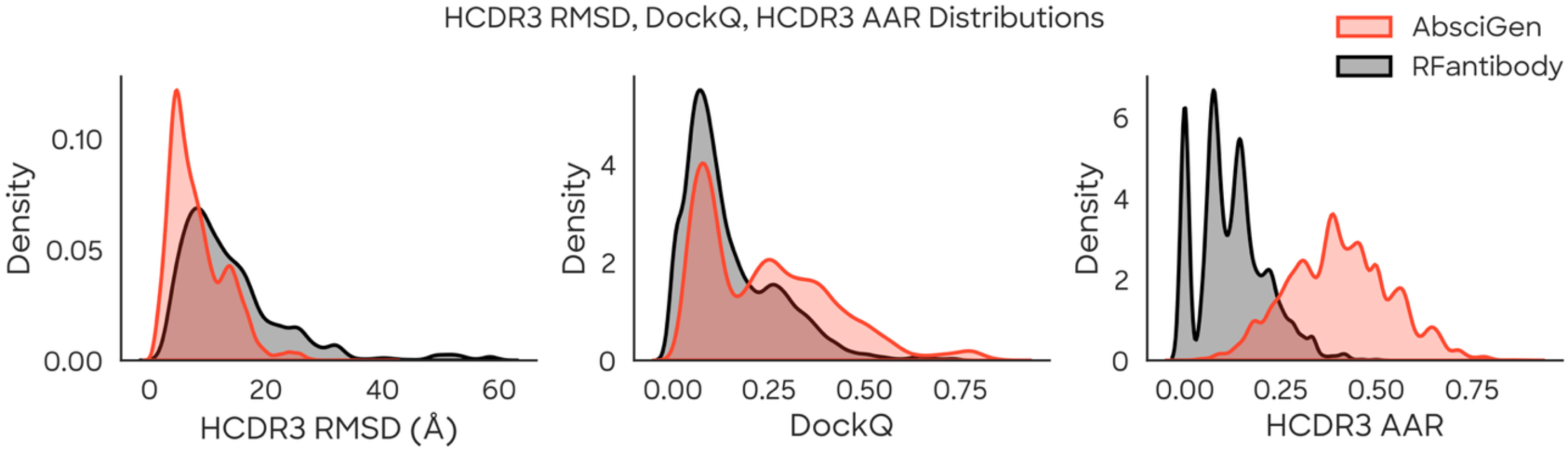
Distribution of supervised benchmark metrics for AbsciGen and RFantibody across all generated samples. Lower HCDR3 RMSD indicates better structural agreement with the native binding pose.

#### 7.4.2 Additional Unsupervised Task Results

**Supplementary Figure 6.**
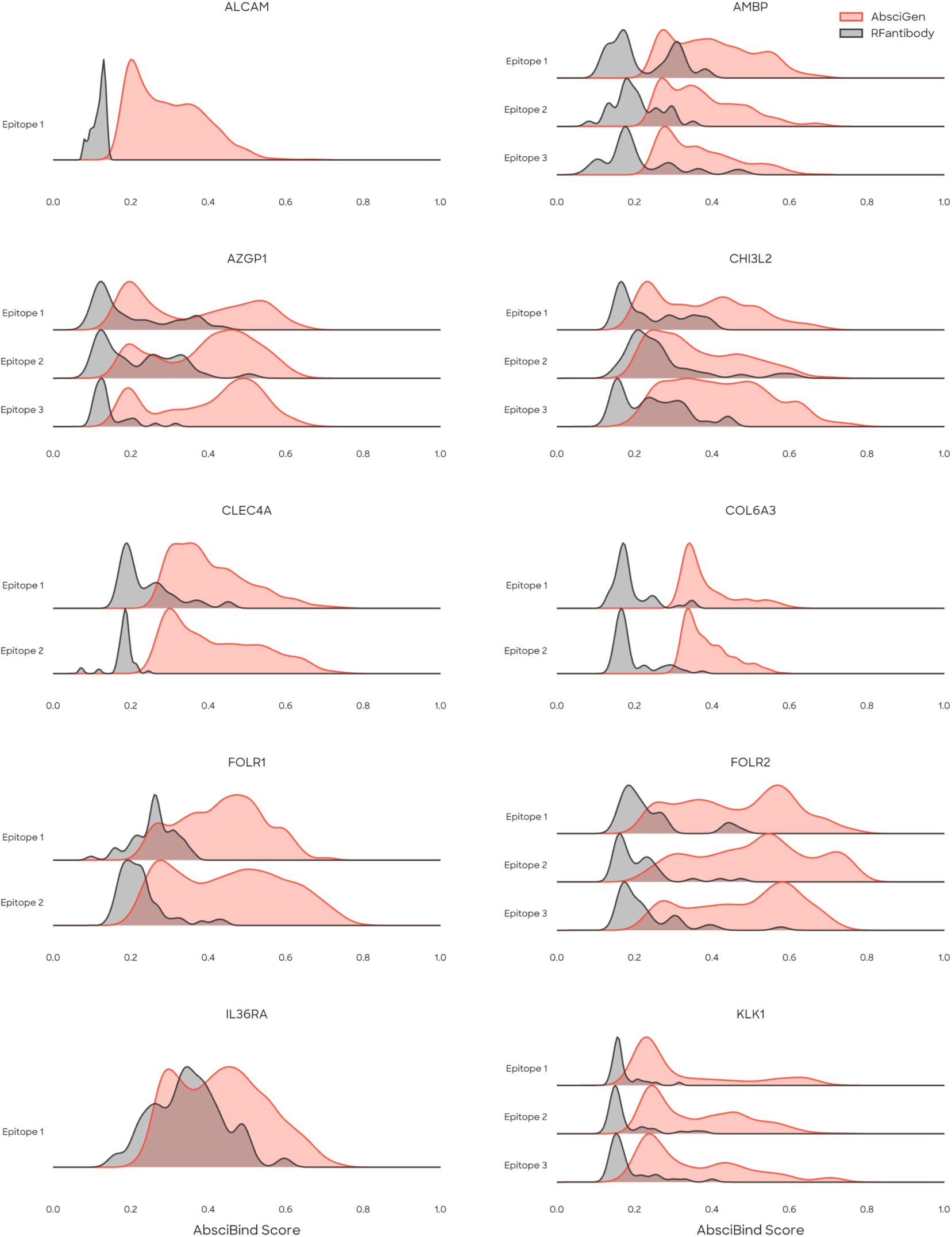
Mean AbsciBind Scores across targets and targeted epitopes for AbsciGen and RFantibody on the unsupervised design task.

**Supplementary Figure 7.**
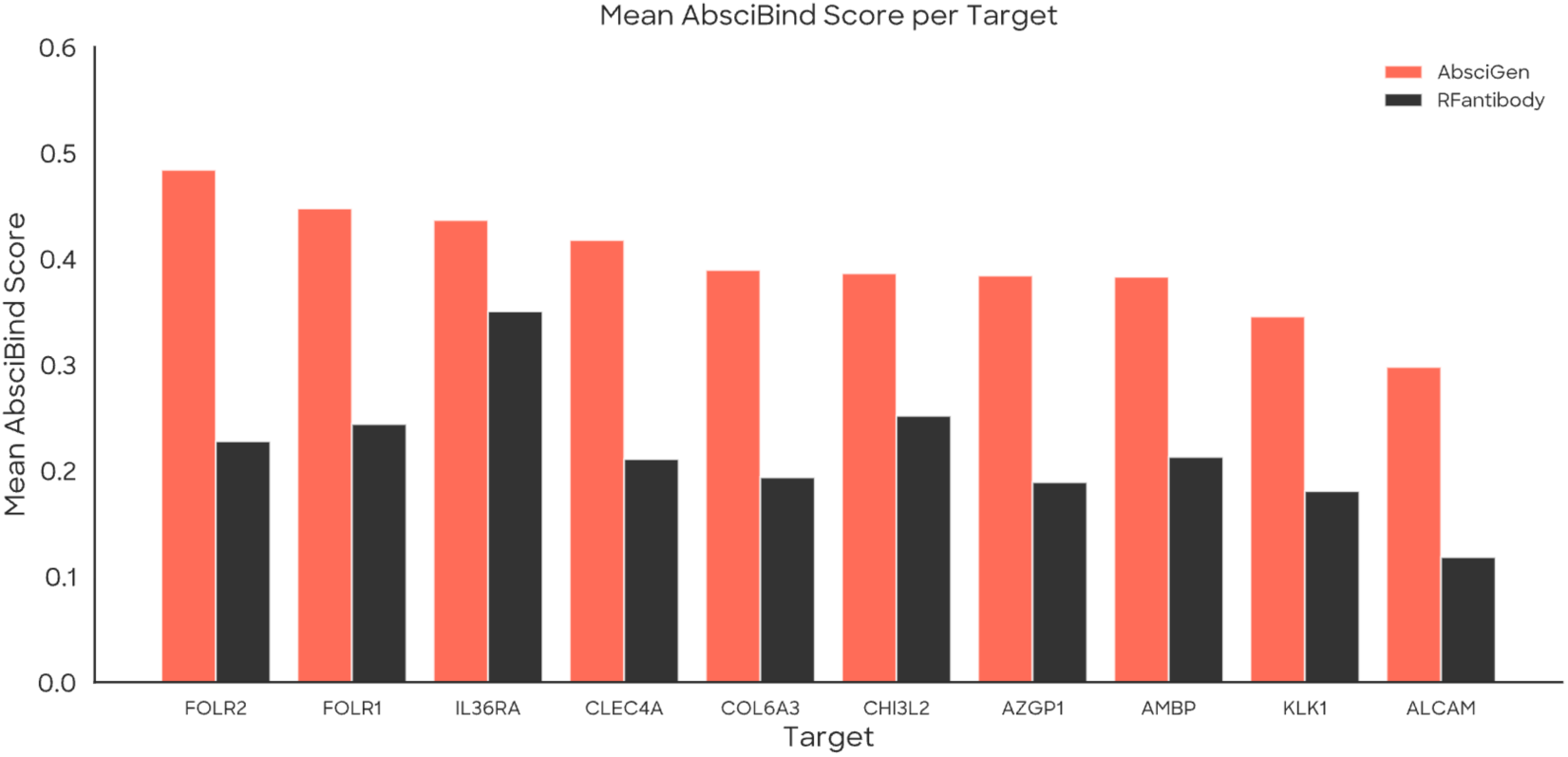
Per-target comparison of mean AbsciBind Scores for AbsciGen and RFantibody on the unsupervised design task.

**Supplementary Figure 8.**
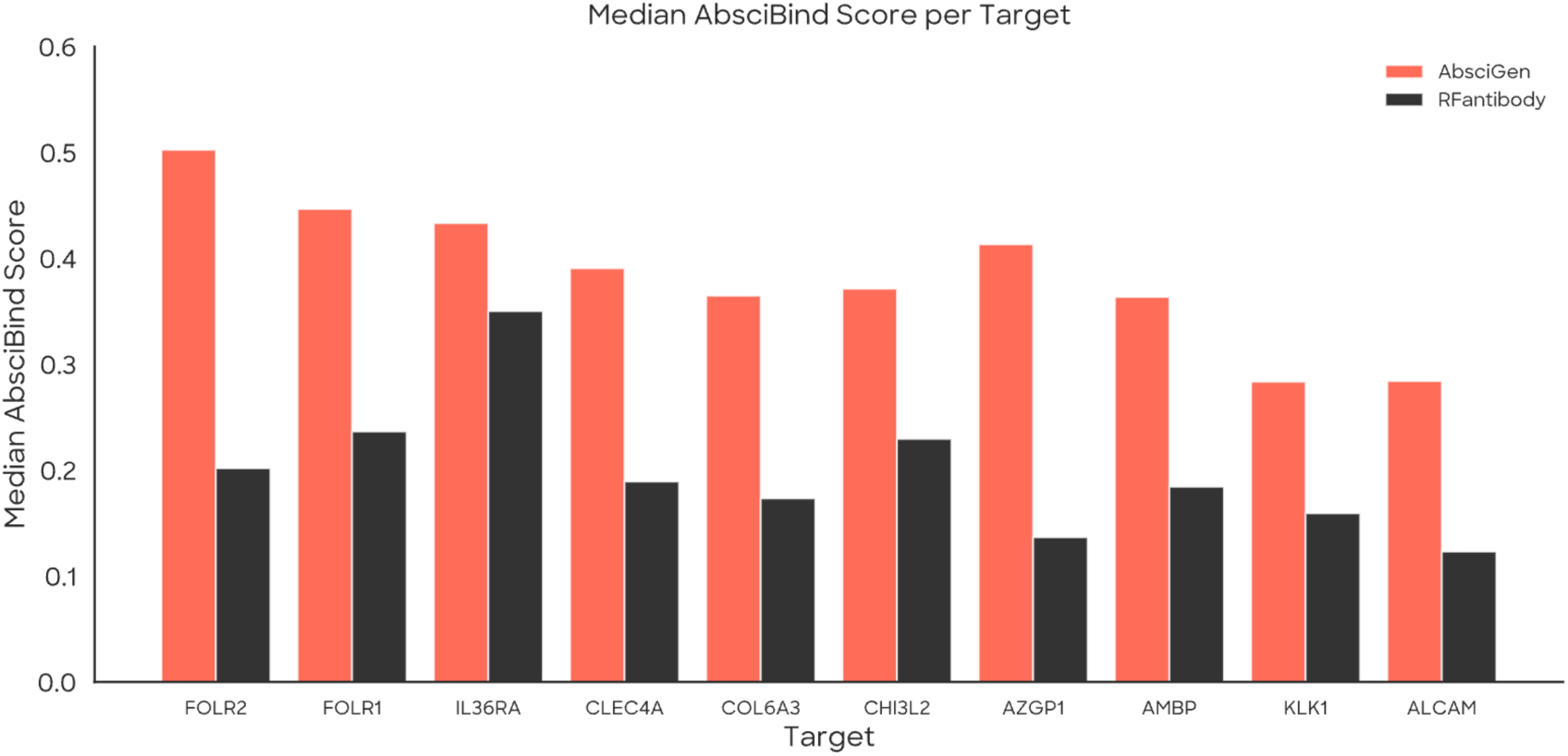
Per-target comparison of median AbsciBind Score scores for AbsciGen and RFantibody on the unsupervised design task. Across all targets, AbsciGen consistently achieves higher median AbsciBind Scores than RFantibody, indicating improved interface quality in its generated designs.

**Supplementary Figure 9.**
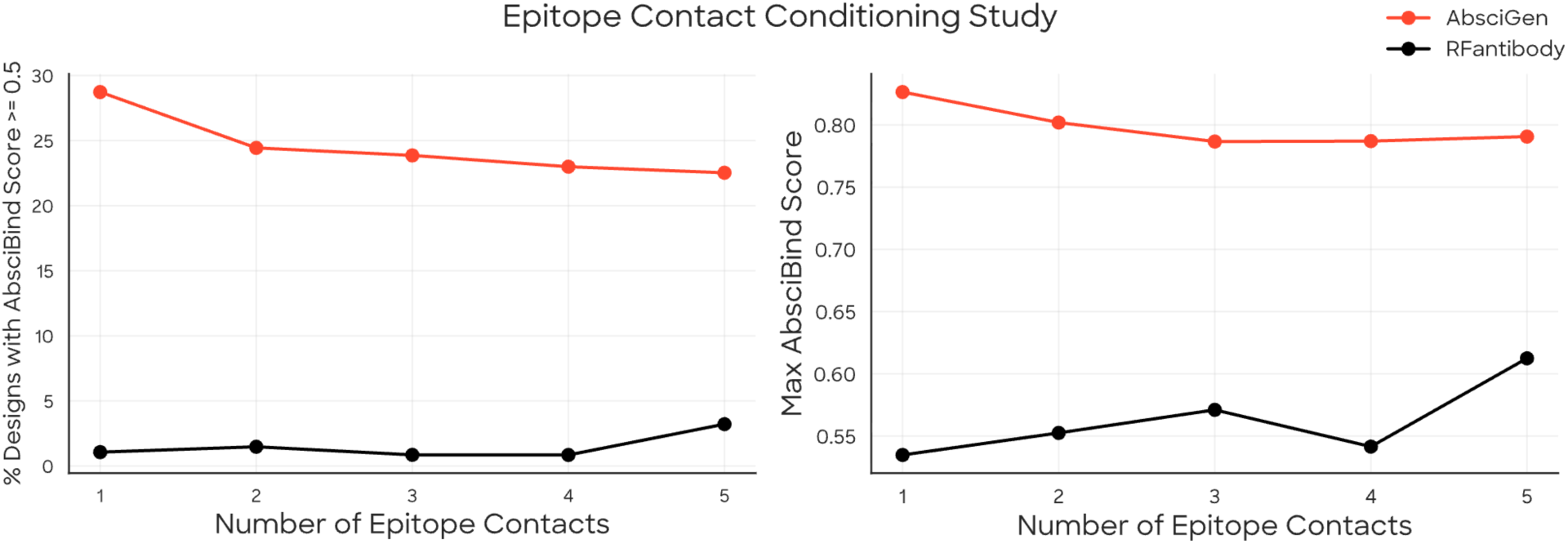
Epitope contact conditioning study. We evaluate how varying the number of provided epitope contact residues impacts the binding, as measured by AbsciBind Score. No significant improvement in the fraction of designs achieving AbsciBind Score ≥ 0.5 is observed with increasing contact information. Notably, the maximum AbsciBind Score for AbsciGen tends to decrease as more epitope contacts are specified, whereas RFantibody shows an increasing trend. We hypothesize that adding contact constraints may restrict the pose diversity for AbsciGen, while RFantibody may benefit from enhanced conditioning, leading to a higher baseline performance. Left: Percentage of designs with AbsciBind Score ≥ 0.5. Right: Maximum AbsciBind Score observed across generated samples for each method.

#### 7.4.3 Sequence Diversity

We assess the diversity of generated HCDR3 sequences by evaluating the fraction of unique sequences and pairwise sequence similarity (PSS) at two aggregation levels.

##### Pairwise Sequence Similarity (PSS)

For variable-length CDR sequences *s*_0_ and *s*_B_, we compute the normalized Levenshtein distance as

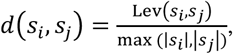

where Lev denotes Levenshtein distance. Pairwise Sequence Similarity is then defined as PSS(*s_i_*, *s_j_*) = 1 − *d*(*s_i_*, *s_j_*), with lower values indicating greater sequence diversity.

**Supplementary Figure 10.**
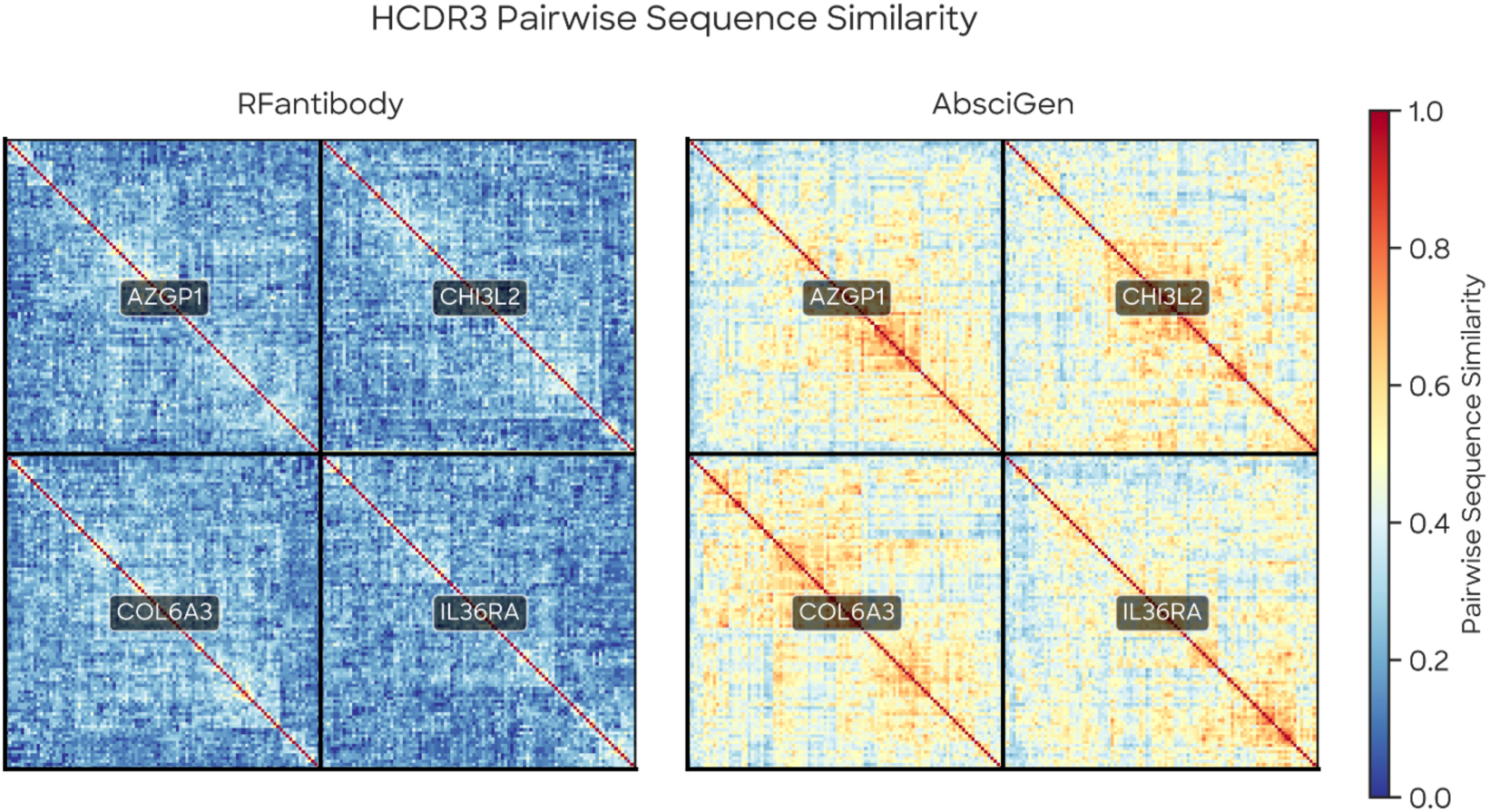
Pairwise HCDR3 sequence similarity matrices, grouped by target. Each pixel shows the PSS score between two sequences generated for the same target (red = high similarity/low diversity, blue = low similarity/high diversity) from our *in silico* benchmark analysis. A single generated sequence was randomly selected for every generated structure, resulting in 102×102 sequence comparisons per target. Sequences are hierarchically clustered within each target block. RFantibody (left) shows more diverse outputs, while AbsciGen (right) generates more similar sequences within each structure. However, both methods achieve high diversity at the target level due to varied backbone generation, as summarized in **Supplementary Table 2**.

**Supplementary Table 2** reveals that AbsciGen exhibits low sequence diversity for a given structure: only 49% of generated HCDR3 sequences are unique (compared to 90% for RFantibody), with a PSS of 0.92 indicating near-identical outputs within each structure. This suggests that AbsciGen’s sequence design module produces limited variation when conditioned on a particular backbone conformation. At the target level, however, both methods achieve 100% unique sequences, indicating that AbsciGen’s generated structures provide sufficient diversity in the context of the full pipeline. In this setting, RFantibody maintains lower PSS (0.21 vs. 0.46), indicating greater sequence diversity.

**Supplementary Table 2.**
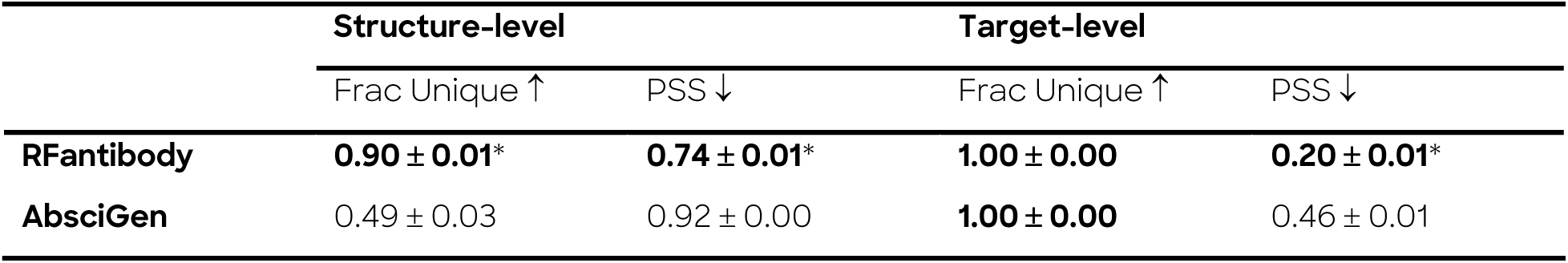
HCDR3 sequence diversity metrics. Fraction of unique sequences and pairwise sequence similarity (PSS) computed at structure-level (within each structure) and target-level (pooled across all structures per target). Metrics computed across 4 targets, reported as mean ± standard deviation. Best values for each metric are highlighted in bold. * indicates *p* < 0.05 by Mann-Whitney U.

**Supplementary Figure 11.**
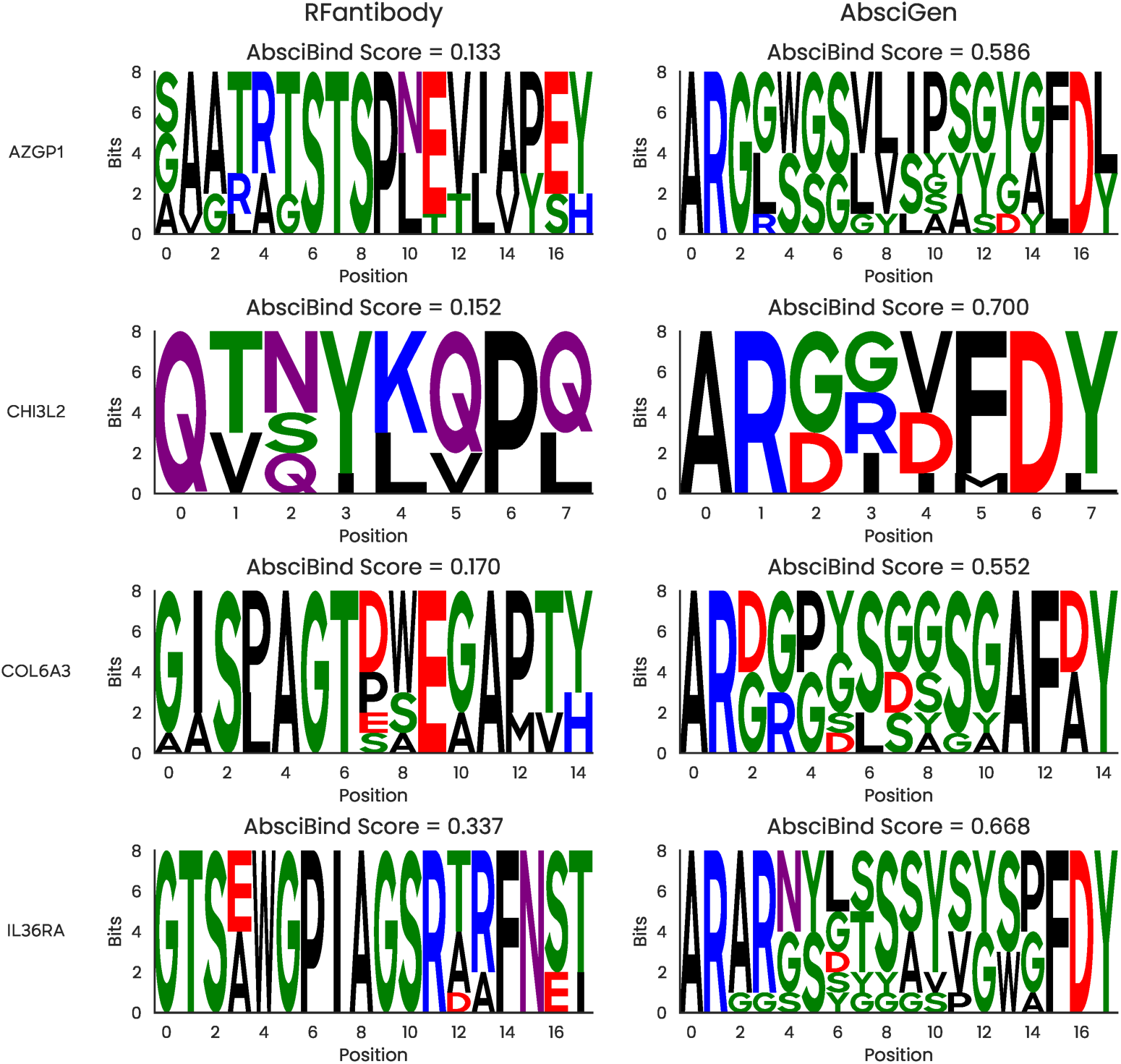
Top generated HCDR3 sequences (*n* = 8) across target, epitope, and CDR-length design specifications per AbsciGen. Configurations were selected based on the highest median ipTM score for each pipeline. Each row shows sequences for the design configuration that achieved the best performance for each method. Configurations producing high scoring samples by AbsciGen are in many cases not found to score highly under RFantibody. Top samples ranked by RFantibody are shown in Supplementary Figure 12.

**Supplementary Figure 12.**
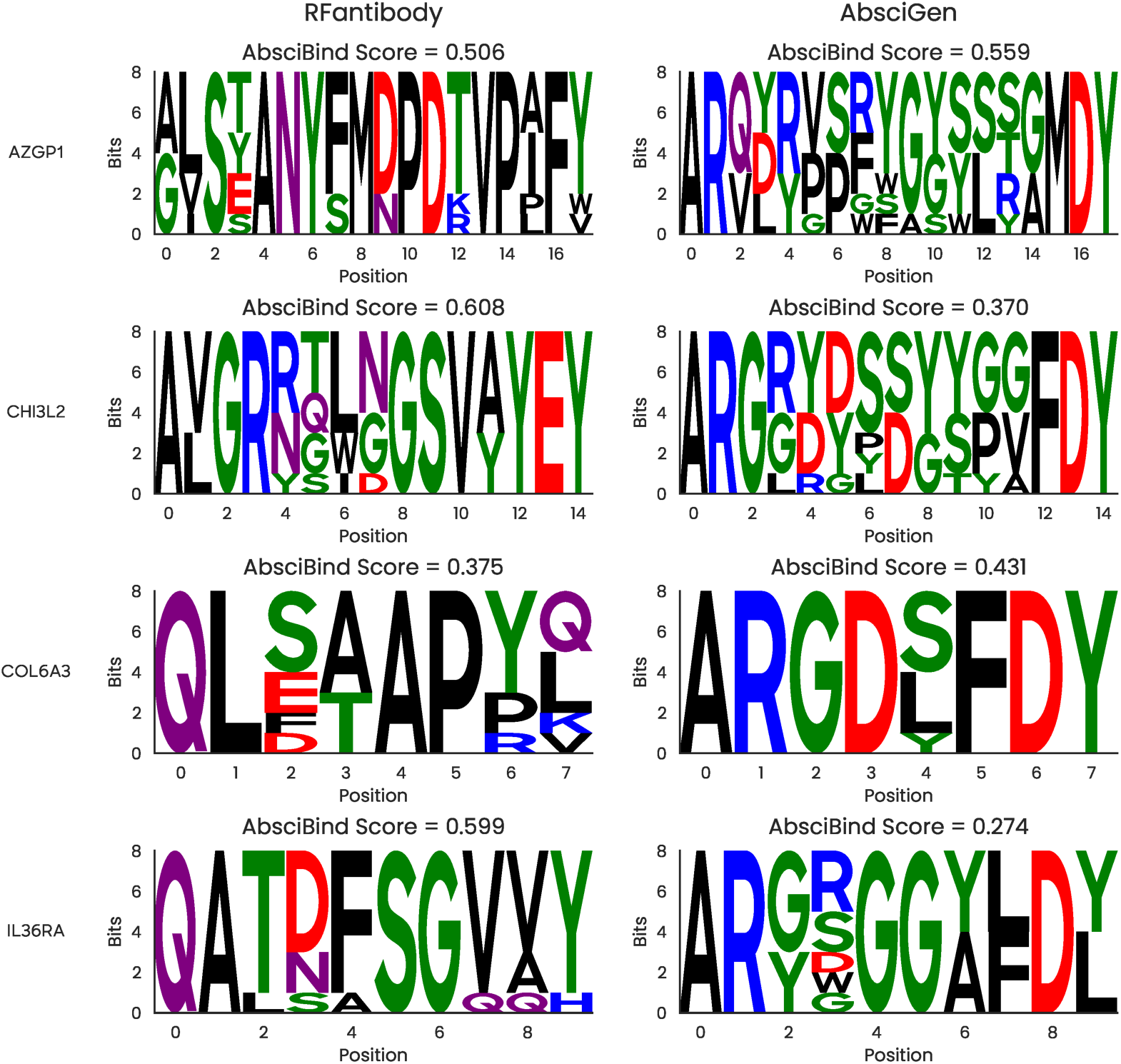
Top generated HCDR3 samples (*n* = 8) of target, CDR-length, and epitope design configurations by median ipTM as ranked by RFantibody. Despite ranking by samples of top configurations under RFantibody, AbsciGen often scores competitively.

#### 7.4.4 Structural Diversity

To quantify structural diversity, we compute pairwise backbone RMSD (*N*, *Cα*, *C*, *0* atoms) within groups of comparable design configurations that share the same antigen target, CDR lengths, and framework lengths. In our benchmarking, we identified over 500 groups of comparable designs for both pipelines, with each group containing up to 48 members with identical topology. We report two complementary metrics, averaging over all valid groups and pairs of group members.

##### Self-Aligned RMSD

As reported in **Supplementary Table 3**, this metric measures conformational diversity by optimally superimposing the target region (HCDR3, LCDR3, or full antibody Fv) onto itself using the Kabsch algorithm [60] before computing RMSD. This captures the range of loop conformations sampled by each method, independent of global orientation.

**Supplementary Table 3.** Self-aligned structural diversity. Pairwise backbone RMSD computed after aligning each region onto itself, measuring conformational diversity independent of global orientation. Metrics are computed across all four unsupervised targets, reported as mean ± standard deviation. Best values in bold. * indicates *p* < 0.001 by Mann-Whitney U.

|  | Antibody RMSD (Å) ↑ | HCDR3 RMSD (Å) ↑ | LCDR3 RMSD (Å) ↑ |
| --- | --- | --- | --- |
| **AbSciGen** | 2.72 ± 1.22 | 1.09 ± 0.22 | 0.78 ± 0.31 |
| **RFantibody** | **4.83 ± 0.69\*** | **1.57 ± 0.23** | **1.26 ± 0.15\*** |

##### Antigen-Aligned RMSD

As reported in **Supplementary Table 4**, this metric measures binding pose diversity by first superimposing structures using the antigen backbone, then computing region RMSD without additional alignment. This captures how diversely each method positions CDR loops relative to the epitope—a metric more relevant to functional diversity in antigen recognition. RFantibody generates greater self-aligned and antigen aligned structural diversity than AbsciGen across all measured regions, although the differences are only statistically significant for self-aligned antibody and LCDR3 RMSD.

**Supplementary Table 4.** Antigen-aligned structural diversity. Pairwise backbone RMSD was computed after superimposing structures on the antigen backbone, measuring binding pose diversity in CDR loop positioning relative to the epitope. Metrics computed across four targets, reported as mean ± standard deviation. Best values highlighted in bold. No differences were determined to be significant by Mann-Whitney U.

|  | Antibody RMSD (Å) ↑ | HCDR3 RMSD (Å) ↑ | LCDR3 RMSD (Å) ↑ |
| --- | --- | --- | --- |
| **AbSciGen** | 6.66 ± 2.21 | 4.18 ± 1.70 | 4.51 ± 1.66 |
| **RFantibody** | **8.16 ± 1.77** | **5.06 ± 1.67** | **5.75 ± 1.60** |

**Supplementary Figure 13.**
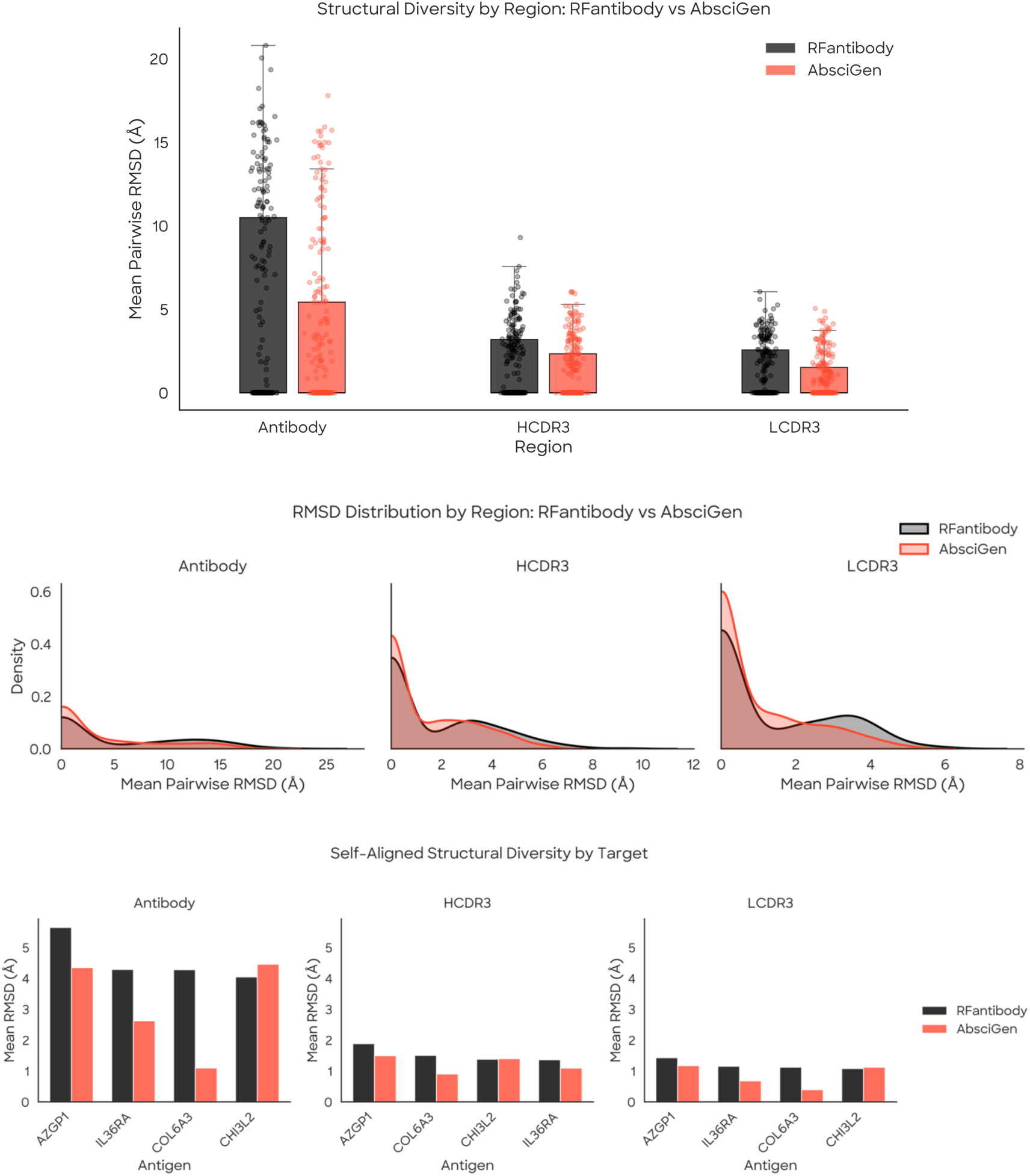
Self-aligned structural diversity comparison. Pairwise backbone RMSD (*N*, *Cα*, *C*, *0* atoms) was computed within groups of comparable antibody structures sharing the same antigen target, CDR lengths, and framework lengths. Each region was optimally superimposed onto itself using the Kabsch algorithm before computing RMSD, measuring conformational diversity independent of global orientation. (Top) Boxplots show the distribution of mean pairwise RMSD across comparable groups for antibody Fv, HCDR3, and LCDR3 regions. Individual points represent comparable groups (*n* = 512 per method). (Middle) Kernel density estimates showing the full distribution of RMSD values. (Bottom) Mean RMSD per antigen target, showing consistent trends across diverse antigens. RFantibody (black) exhibits greater structural diversity than AbsciGen (red) across all regions and targets.

### 7.5 AbsciBind Extended Methodology

Effectively scoring and ranking designs is fundamental to the success and scalability of an AI pipeline for antibody design. While folding models have shown strong performance for scoring and filtering general protein binder designs, their potential to translate for antibodies remains unclear, given reports of limited accuracy for antibody-antigen complex prediction [33, 34].

We curate a protocol, AbsciBind, as a scoring method for antibody-antigen complexes, leveraging the advantages of existing folding approaches and identifying workarounds to their limitations. Folding algorithms such as AlphaFold (AF) [7] and AlphaFold-Multimer (AFM) [30] take protein sequences, structural templates from homologs, and multiple sequence alignments (MSAs) as inputs. For AF and AFM, templates are encoded by amino-acid identity, Cβ distance matrices, and backbone torsion angles. Prior work demonstrated that AFM confidence metrics can be repurposed to score candidate (“decoy”) structures by either “disguising” the decoy as the model’s output from a previous (fictitious) recycling iteration and feeding it back through the network, in the style of AlphaFold-Multimer Initial Guess (AFM-IG) [61], or supplying the decoy as a template, using empty MSAs, masking side chains except for the Cβ atom, adding “virtual” Cβ atoms for glycine residues, and replacing template residues with gap symbols. The latter approach, known as AF2Rank [62], achieves strong decoy ranking performance on single-chain proteins. Although AFM masks inter-chain Cβ distances during training and inference, exposing these distances via minor inference-time modifications enables the model to leverage cross-chain template information without retraining, as demonstrated by the AF_Unmasked method [45]. Motivated by this result, we attempted to apply AF2Rank with cross-chain templates to antibody–antigen complexes, but our results showed that this approach yielded high false-negative rates (**Supplementary Figure 14**), with correct docked poses often not preserved in AF2Rank outputs. We attribute this failure to masking template amino-acid tokens, which causes the model to ignore critical information from antibody–antigen complex templates.

This observation inspired us to implement several adjustments to the native AFM protocol: we provide the amino acid sequence of an antibody-antigen complex as the input; we supply the designed or decoy structure as a multimer template; we retain template amino-acid tokens instead of replacing them with gap symbols; we mask all template side chains except for the Cβ atoms; we disable AFM’s default masking of inter-chain template distances; and we use single-sequence mode. To ensure deterministic inference, we disable dropout and Evoformer residue masking. We call this updated protocol AbsciBind.

To assess the impact of implementing these changes prior to utilizing the AbsciBind protocol to select AbsciGen designs as part of the Origin-1 pipeline, we tested the AbsciBind protocol, AFM-IG, and AF2Rank’s abilities to recover native antibody-antigen poses by deploying these models on a set of antibody-antigen complexes released after the training cutoff for AFM v2.3. We found that while AFM-IG and AF2Rank (with cross-chain templates) often failed to recover the native poses, our AbsciBind protocol succeeded (**Supplementary Figure 14**). We identified one AFM v2.3 model checkpoint (`model_2_multimer_v3`) that best recapitulated experimental antibody-antigen complexes in the PDB when used in our AbsciBind pipeline (**Supplementary Figure 15**). These results motivated us to proceed with AbsciBind as our primary protocol for establishing an antibody design scoring method.

#### Filtering

Consistent with the logic underlying computational filtering methods that rely on folding methodologies for protein design quality assessments, we assumed that AbsciGen designs should be discarded if the 3D structure predicted by the AbsciBind protocol deviated substantially in tertiary and/or quaternary structure from the AbsciGen structure provided as a template input. To this end, we computed structural consistency scores that quantified the similarity between the designed and predicted complexes and used these scores to filter our designs. Specifically, we first merged the heavy and light antibody chains into a single Chain A, and all antigen chains into a separate Chain B. We then computed the ligand root mean square deviation (L-RMSD) as follows: (1) the ligand and receptor were respectively defined as the shorter and longer of Chains A vs. B; (2) all non-backbone atoms were removed from both the designed and predicted complexes; (3) the rigid-body transformation that minimized the RMSD between the receptor chains was computed; and (4) the L-RMSD was defined as the RMSD between the ligand chains after applying this transformation. Any prediction with an L-RMSD greater than 5 Å relative to the designed complex was discarded.

#### AbsciBind Protocol ipTM Score

AFM’s interface-predicted TM (ipTM) score [30] measures the model’s confidence on the global packing of a protein complex. In applying the ipTM score to the antibody-antigen problem, the value of this score is also influenced by the model’s confidence in the relative arrangement between heavy and light antibody chains, as well as between antigen chains if several are present. To reduce the weight of contributions to ipTM coming from intra-antibody or intra-antigen residue pairs, we first merged the antibody heavy and light chains into a single chain Ab of length L_LM_, and all antigen chains into a single chain Ag of length L_LN_. Then, ipTM for this fictitious dimer can be expressed as

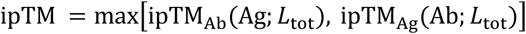

where *L*_tot_ = *L*_Ab_ + *L*_Ag_ is the total number of amino acids in the complex and, for any two chains *A*, *B* in a complex and any *L* > 0, we define

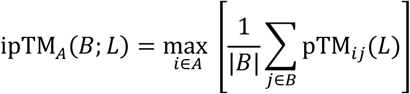

where

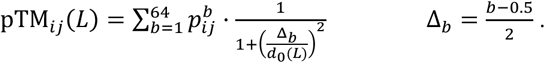

The probabilities 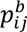 are outputted by AFM and satisfy 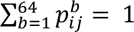. The normalization factor is defined as,

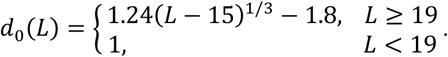

Because the normalization factor *d*_0_(*L*_tot_) depends on the antibody length in the original formulation of ipTM, the relative ranking of designs targeting the same antigen but with different CDR lengths can be systematically affected. To mitigate this effect, we introduce an *Antibody-Aligned* ipTM, defined as:

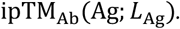

Related one-way alignment scores and similar modifications to the normalization factor d_0_ were previously introduced in [63].

#### ***In silico*** Benchmarking of the AbsciBind Protocol

To assess the AbsciBind protocol’s appropriateness for scoring and selecting AbsciGen designs, we tested AbsciBind’s ability to discriminate true binders from non-binders in an experimental set of eight antibody-antigen systems [21, 46]. We compared the AbsciBind protocol’s performance with the performance of six established reference approaches (GeoFlow-V3, Geoflow-V2, AFM-IG, Protenix, Boltz-2, and Chai-1) [27].

Results showed that AbsciBind achieves the strongest average binder-non-binder discrimination performance as measured by AUROC (**Figure 5**), using the maximum ipTM score across all five AFM v2.3 model checkpoints. Consistent with the prior observation that `model_2_multimer_v3` (model 2) checkpoint accurately recapitulates ground-truth PDB structures, we find that this “Selected AbsciBind ipTM” score (model 2 only) closely tracks “Best AbsciBind ipTM” score (best from all 5 models) across targets. In several cases, including ACVR2B, TSLP, IL36R, and C5, the Selected score achieved nearly identical, and in one case higher, AUROC values compared with the Best ipTM score. Importantly, using only model 2 yields an approximately 80% reduction in runtime compared with evaluating designs with all five models. Given this substantial runtime reduction and the strong classification performance, model 2 was selected to support the AbsciBind protocol for the remainder of the present effort.

We defined a final AbsciBind Score as the arithmetic mean of the default AbsciBind protocol ipTM score (computed over all antibody and antigen chains and interfaces) and the Antibody-Aligned ipTM score. We design this metric to integrate a global interface score with an antibody-aligned, consistently normalized assessment of the antibody-antigen interface’s quality and use this score to evaluate AbsciGen designs for *de novo* library design (§2.4.3) and to select mutant variants for lead optimization efforts (§2.4.4).

**Supplementary Figure 14.**
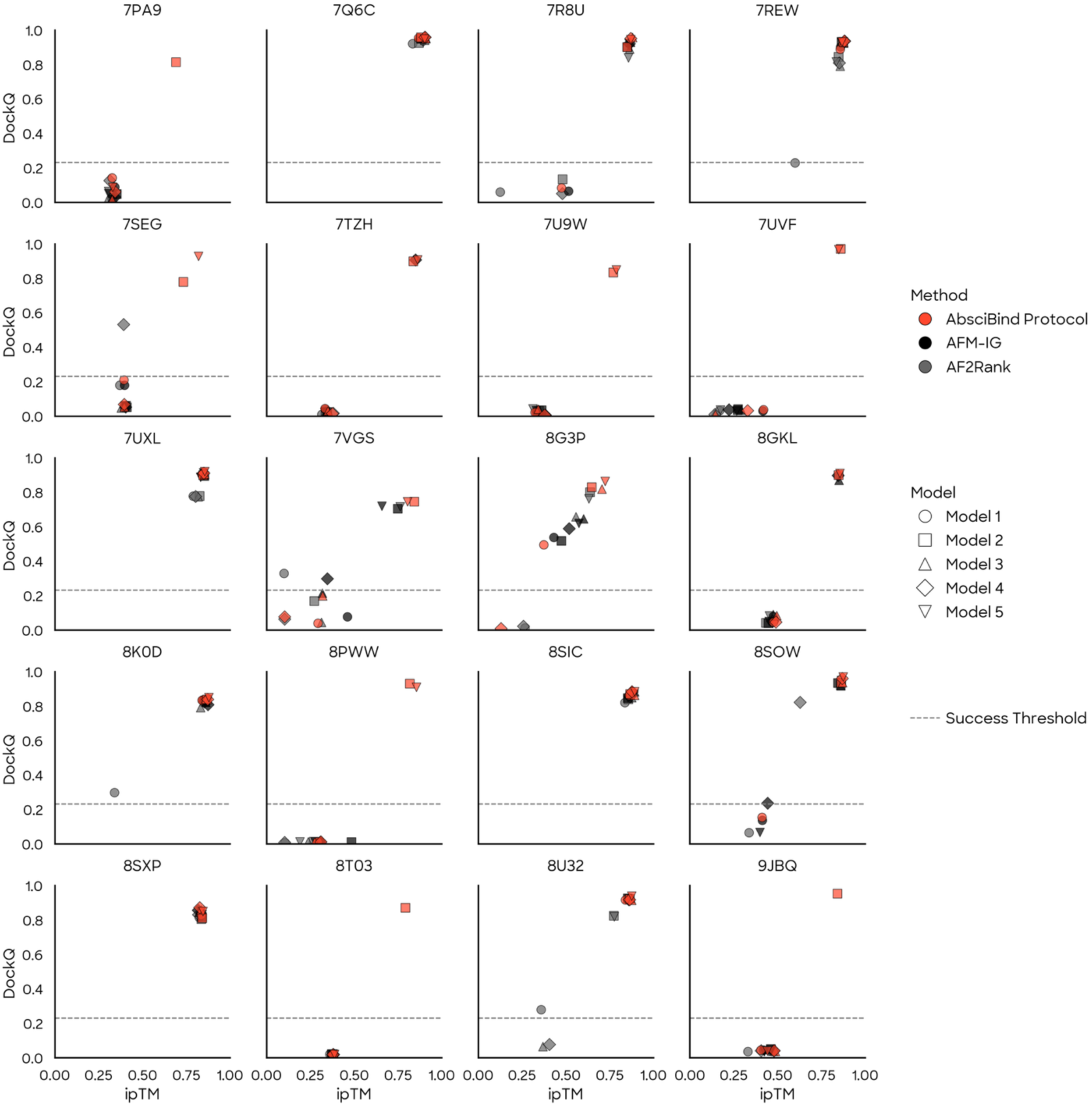
Distribution of DockQ and ipTM scores for AlphaFold-Multimer (AFM) predictions generated using the initial-guess (AFM-IG), AF2Rank, and AbsciBind protocols. Colors and markers indicate the protocol used. The gray dashed line at y = 0.23 marks the CAPRI threshold separating incorrect from acceptable predictions based on DockQ. DockQ scores are computed by comparing the merged antibody heavy–light chains against the antigen chain. All complexes were released after the AFM v2.3 training cutoff date (2021-09-30) and are non-redundant with respect to antigen sequences at a 40% sequence similarity threshold.

**Supplementary Figure 15.**
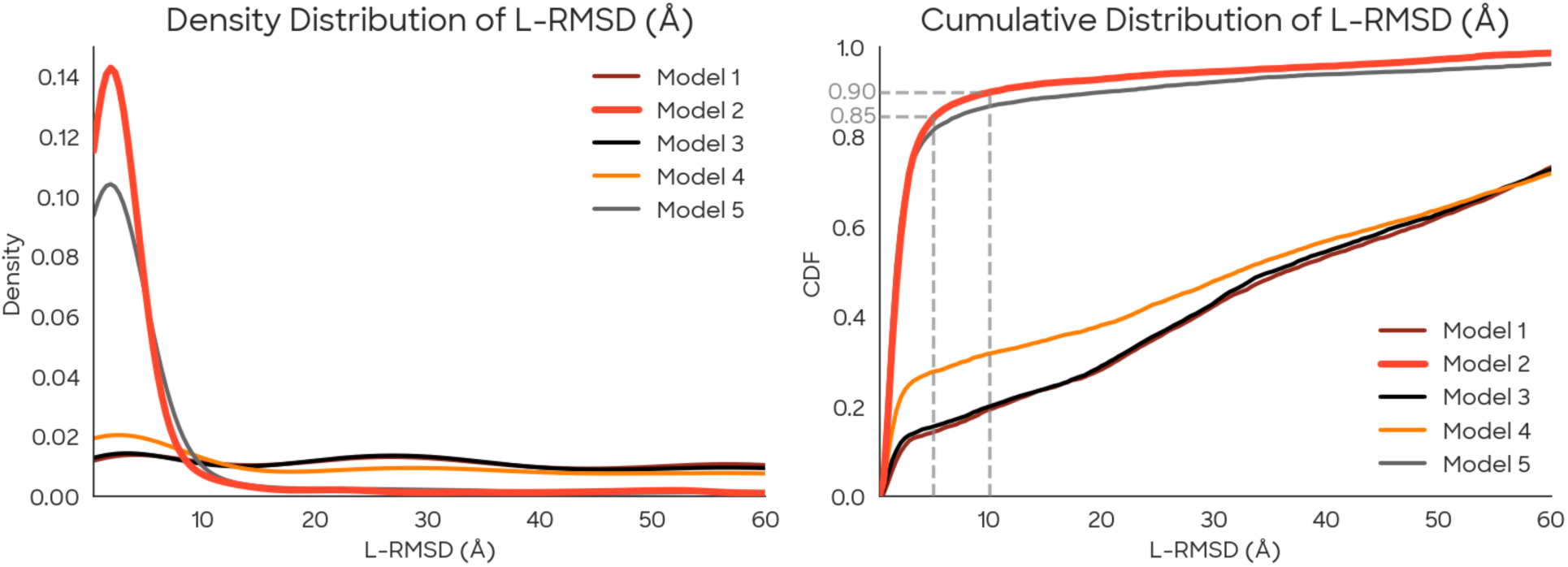
Distribution of L-RMSD values for SAbDAb-generated structures. The left panel shows the histogram of scores, while the right panel shows the cumulative distribution function (CDF), indicating the fraction of structures below a given L-RMSD threshold. Notably, 85% of generated structures fall below an L-RMSD threshold of 5 Å, while 90% fall below 10 Å.

### 7.6 Library Design

#### 7.6.1 Structure and Sequence Search Strategy

**Supplementary Figure 16.**
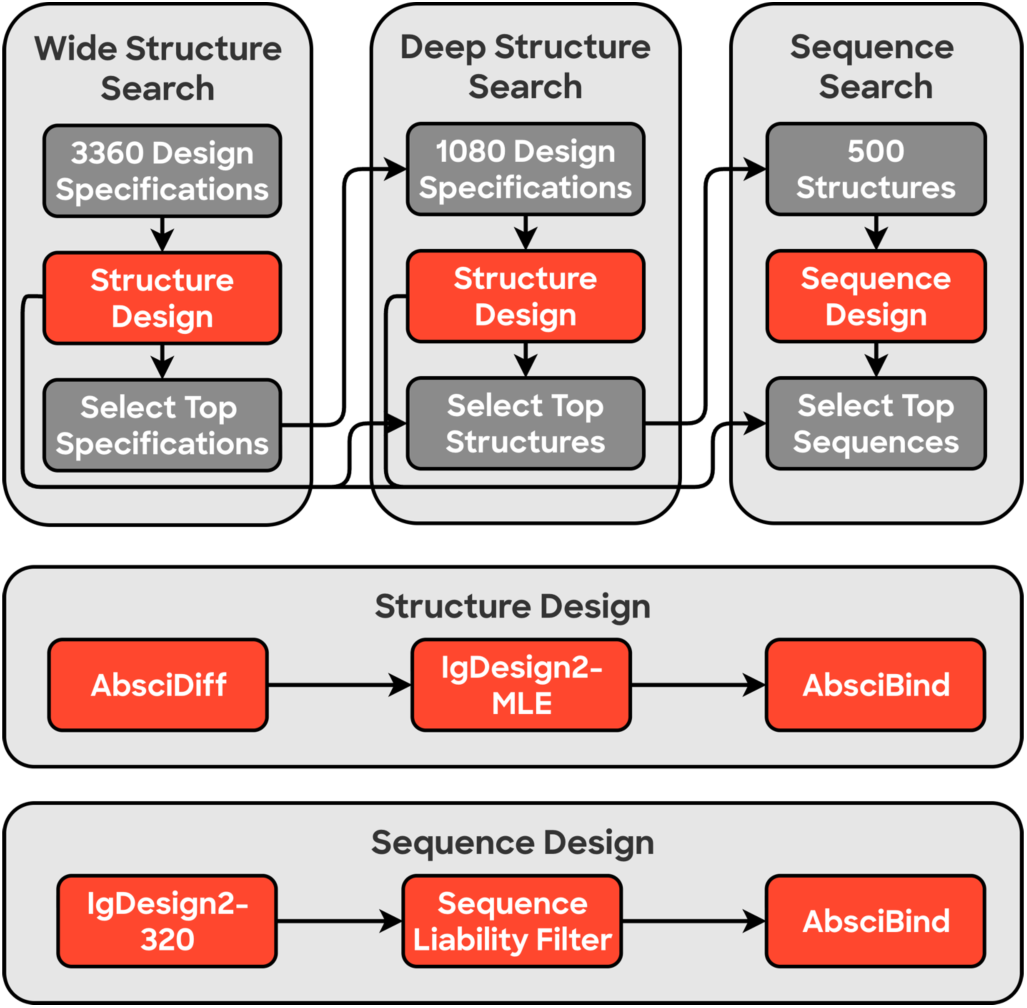
Schematic illustrating inference-time search strategy involving progression through Wide Structure Search, to identify optimal design specifications, Deep Structure Search, to select top backbone structures, and Sequence Search, to select top sequences per backbone structure for experimental validation.

**Supplementary Figure 17.**
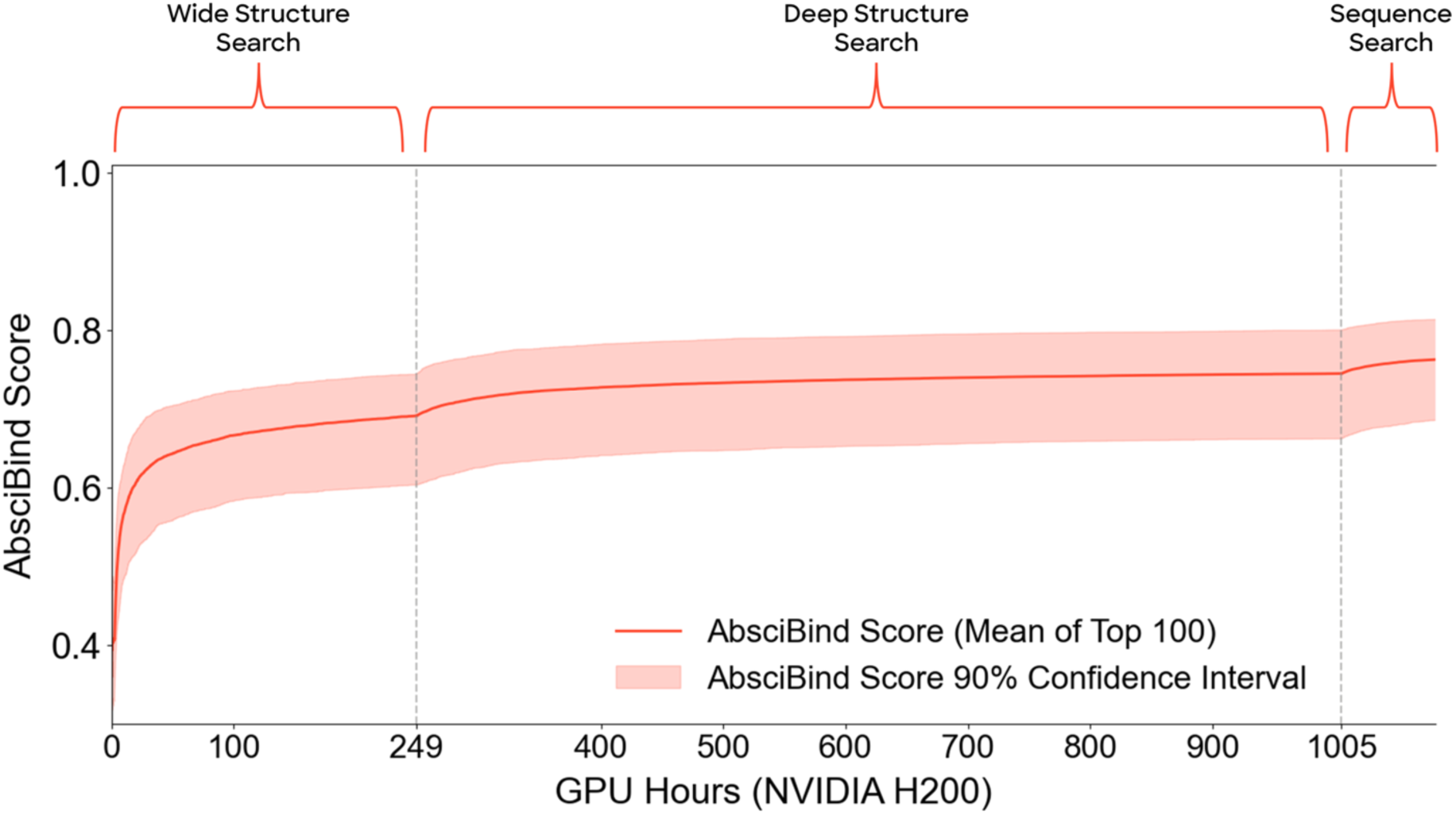
Concentrating compute resources on high-confidence designs within defined search windows improves quality of top designs as assessed by AbsciBind Score. Solid line represents the mean (across multiple targets) of the top score within each target. Shaded areas represent 90% confidence interval.

##### Wide Structure Search

The first structure design step aimed to identify the design specifications (formatted as tuples of FWRs, CDR lengths, and epitope residue samples) that yielded structure-sequence pairs with the highest Antibody-Aligned ipTM Scores. 3360 combinations of design inputs were sampled and provided as inputs to AbsciDiff. AbsciDiff was run with *M* = 24 diffusion samples per trunk sample and *K* = 3 selected samples out of *M* diffusion samples, resulting in 3360 · *M* = 80640 generated structure samples and 3360 · *K* = 10080 selected structure samples. IgDesign2 generated one top sequence per structure, and each structure-sequence pair was scored using AbsciBind.

Of these 10080 structures, we selected the top 5% of design strategies by maximum Antibody-Aligned ipTM Score (across the three structures) and the top 5% by median Antibody-Aligned ipTM Score (across the three structures) for a total of 1008 design strategies (with some redundancy, where the max and median overlap). These design strategies (config files) were then oversampled at a rate of 10X, providing a total of 10080 design specifications for Deep Structure Search, with either ten or twenty repeats of each design specification. We evaluated this approach against alternative filtering strategies of using the top 4%, 7%, or 10% of configs by the maximum + median strategy described above and found that the current method performed favorably for maximizing Antibody-Aligned ipTM Score.

##### Deep Structure Search

Following Wide Structure Search, we executed a second structure design step, Deep Structure Search, to exploit the favorable configurations identified through Wide Structure Search and comprehensively sample the accessible structure space within each design specification.

During Deep Structure Search, AbsciDiff, IgDesign2, and AbsciBind were run with parameters identical to those used during Wide Structure Search, resulting in 10080 · *M* = 241920 generated structure samples and 30240 selected/scored structure samples with one sequence selected per structure.

All 30240 structure-sequence pairs produced during Deep Structure Search were pooled with the 10080 structure-sequence pairs from Wide Structure Search for a total of 40320 structures. Designs were then filtered to remove those for which the L-RMSD (**Supplementary Table 6**) between the AbsciBind protocol-predicted structure and the AbsciDiff-designed structure exceeded 5 Å. Out of the top 10000 remaining structures by Antibody-Aligned ipTM Score, 500 structures were advanced to Sequence Search (see below). The first 25% of structures were selected by highest Antibody-Aligned ipTM Score. The remaining 75% of structures were selected by minimum Intersection Score (**Supplementary Table 6**). The rationale behind leveraging multiple selection strategies was to ensure inclusion of designs that prioritize both recall, by selecting structures scored highly by Antibody-Aligned ipTM Score (first 25% selection), and precision, by selecting structures scored highly by both Antibody-Aligned ipTM Score and AbsciBind ipTM score (last 75% selection). To ensure structural diversity among this selection, structures were clustered via agglomerative clustering using the antigen-aligned-conserved-residue-RMSD metric (**Supplementary Table 6**) with a distance threshold of 10 Å. The percent of structures selected from a single structural cluster was limited to 30% during implementation of the 25%/75% selection strategy.

##### Sequence Search

Sequence Search searches for top-scoring sequences for the top backbone structures identified through Wide Structure Search and Deep Structure Search. 320 sequences were sampled from IgDesign2 for each of the 500 selected structures. These sequences were filtered to remove sequences with “critical” liabilities (**Supplementary Table 7**) and deduplicated. The top twenty sequences per structure, as ranked by IgDesign2 pseudo-perplexity (**Supplementary Table 6**), were scored with the AbsciBind protocol (using their reference structure as a template) and considered for final selection.

For final selection, sequences from Wide and Deep Structure Search were pooled with designs from Sequence Search. Any sequence with an L-RMSD between the design structure vs. AbsciBind-folded structure of greater than 5 Å was removed. For each structure, the top sequence was selected by Intersection Score. The top 95 structure/sequence pairs were then selected by AbsciBind Score (**Supplementary Table 6**), restricting the number of “Other” sequence liabilities permissible (**Supplementary Table 7**). Additional criteria were imposed to promote design diversity: the permitted number of replicates of each unique HCDR3 sequence was restricted to three; the number of replicates of each unique (HCDR3 sequence, `cdr_lengths`) pair was capped at one (where `cdr_lengths` is an ordered list of CDR lengths); and the permitted proportion of sequences from a single structural cluster was capped at 30%.

#### 7.6.2 CDR length distributions used to guide antibody design

**Supplementary Table 5.**
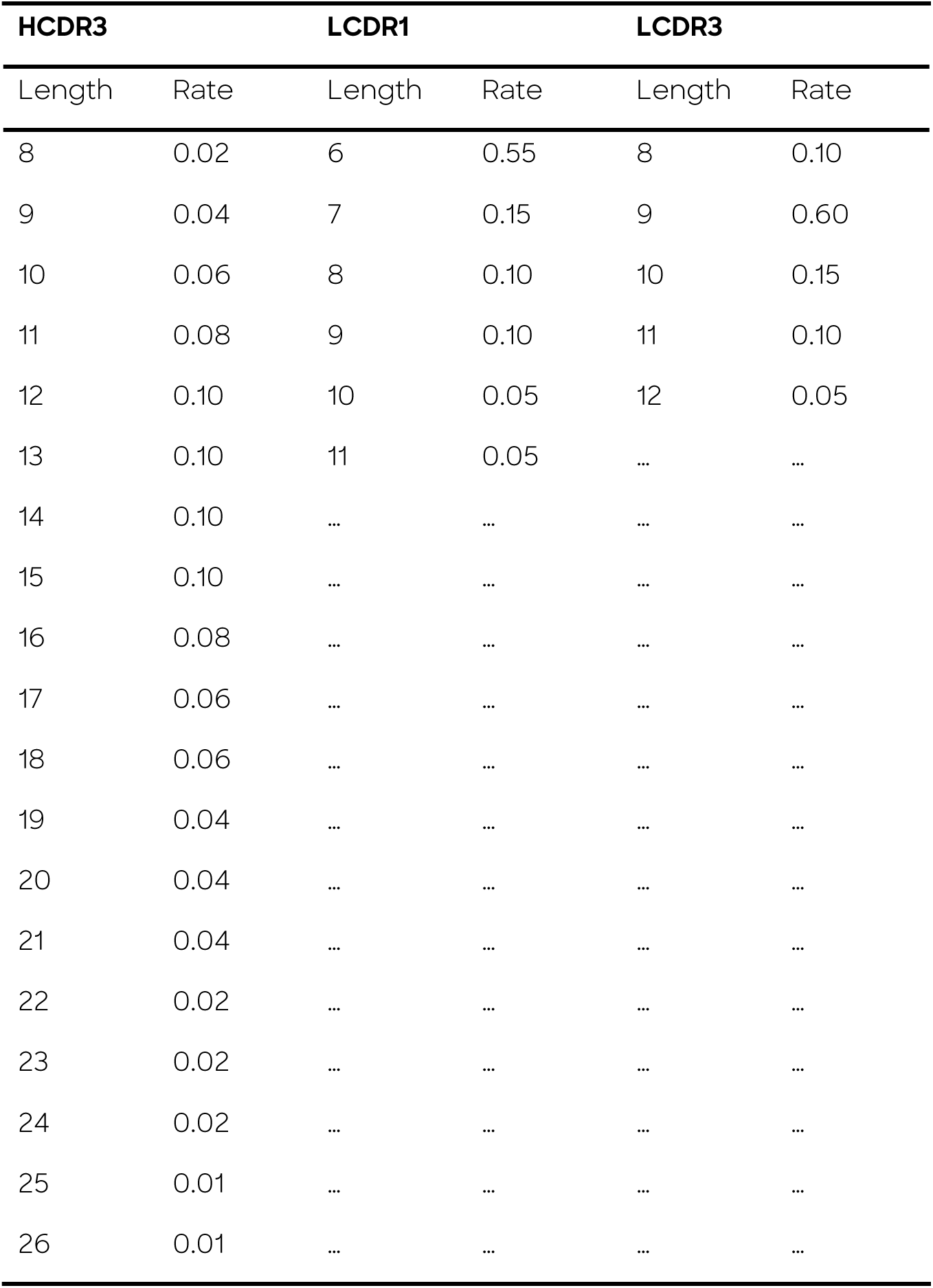
Normalized distributions of HCDR3, LCDR1, and LCDR3 lengths from AbData. Lengths for individual CDRs were sampled independently from these distributions. Length = CDR length; Rate = normalized rate of occurrence in AbData.

#### 7.6.3 Metrics used to Score AbsciGen Designs

**Supplementary Table 6.** List of metrics used to evaluate Origin-1 generated structures and sequences.

| **Metric** | **Description** | **Search Stage(s)** |
| --- | --- | --- |
| **Antibody-Aligned ipTM Score** | A custom asymmetric ipTM score that merges all antibody chains and all antigen chains so that only antibody-antigen interactions are considered. | Wide Structure Search, Deep Structure Search, Sequence Search |
| **AbsciBind ipTM Score** | Interface-predicted template modeling (ipTM) score from the AbsciBind protocol. | Deep Structure Search, Sequence Search |
| **AbsciBind Score** | Average of Antibody-Aligned ipTM Score and AbsciBind ipTM Score. | Sequence Search |
| **Intersection Score** | A design's minimum rank when comparing its Antibody-Aligned ipTM Score and AbsciBind ipTM scores, defined as: $\min(\text{rank}(\text{complexes, "AbsciBind ipTM Score"}), \text{rank}(\text{complexes, "Antibody-Aligned ipTM Score"}))$. | Deep Structure Search, Sequence Search |
| **Ligand RMSD (L-RMSD)** | The substrate-aligned root mean square deviation of atomic positions for the ligand between two complexes | Deep Structure Search, Sequence Search |
| **Antigen-aligned conserved residue RMSD** | The antigen-aligned root mean square deviation of residue positions which are structurally conserved across antibody frameworks. These positions are Chothia [64] scheme IDs 36-39 and 89-92 on the heavy chain and scheme IDs 35-38 and 85-88 on the light chain. Complexes are aligned on the antigen via the Kabsch algorithm [60]. RMSD is then calculated across these conserved positions. | Deep Structure Search, Sequence Search |
| **IgDesign2 pseudo-perplexity** | The perplexity of the sequence under the IgDesign2 logits. | Sequence Search |

#### 7.6.4 Sequence liabilities considered in antibody design

**Supplementary Table 7.** List of sequence liabilities considered while designing Origin-1 libraries for *in vitro* experimentation.

| Sequence Liability | Category |
| --- | --- |
| N-Glycosylation - The motif NXS or NXT with X any residue other than proline | Critical |
| 3-W - Five consecutive tryptophan residues in a single CDR. | Critical |
| 4-G - Four consecutive glycine residues in a single CDR. | Critical |
| 5-Y - Five consecutive tyrosine residues in a single CDR. | Critical |
| 5-S - Five consecutive serine residues in a single CDR. | Critical |
| 3-G - Three consecutive glycine residues in a single CDR. | Other (4 max) |
| 3-S - Three consecutive serine residues in a single CDR. | Other (13 max) |
| 4-S - Four consecutive serine residues in a single CDR. | Other (3 max) |

### 7.7 Lead Optimization

#### Variant Scoring

We computed the masked marginal (MM) likelihood of a mutant sequence as a fitness score for the protein language models, as this approach has been shown in benchmarking tasks to perform well with a low computational requirement, especially for single-mutants [56]. We calculated the difference between AbsciBind Scores associated with the parent relative to the mutant and used this difference as the fitness score from the AbsciBind protocol.

#### Variant Selection

Mutable positions on both heavy and light chain included the CDRs, structure-inferred paratope residues (defined as those within 5 Å of the antigen using the AbsciBind predicted structure). Additional mutable positions are included in FWR2 and FWR3, excluding selected conserved motifs. All possible single-mutants were generated for the defined mutable positions and scored with the ESM ensemble [55–57], AbLang2 [58], and the AbsciBind protocol as fitness models, along with the Therapeutic Antibody Profiler (TAP) [65] and BioPhi [47] to assess *in silico* developability and humanness. Mutant sequences that introduce chemical liabilities (e.g. N-glycosylation motifs) were also excluded.

#### Library Creation for First Round of Optimization

We selected 94 single-mutant variants per *de novo* binder. To generate these libraries, we first included an alanine scan of all structurally inferred paratope residues, all human germline reversions, and any positions with improved fitness by both AbsciBind and a sequence model (either of ESM-ensemble, or AbLang2). For the ESM ensemble, improved fitness was defined as a median score of > 0.0 across all component models. We subsequently selected the highest-scoring (by AbsciBind Score) structural mutants, allowing up to four mutants per CDR/paratope position and two mutants per mutable framework position. The remaining library budget was filled with sequence mutants (by ESM ensemble only) allowing no more than two mutants per CDR/paratope position and one mutant per framework position. Fitness thresholds and positional budgets were adjusted per design as necessary to fit the library budget, maintain balance across categories, and ensure positional diversity. **Supplementary Table 8** shows the categorical breakdown of variants selected by method in each library. We measure the binding affinity of library variants by SPR using the protocol described in Supplement §7.8.2.

**Supplementary Table 8.**
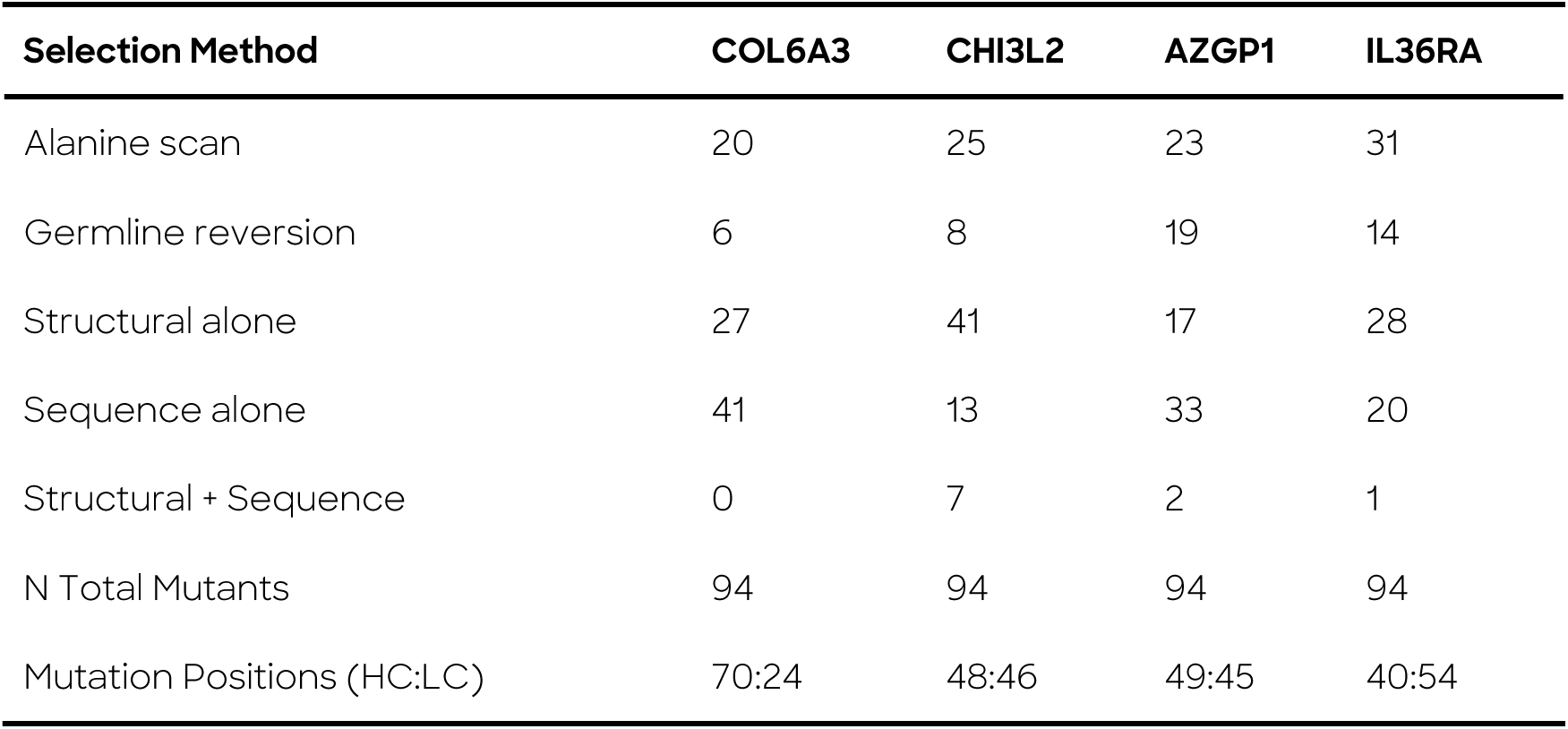
Count of single-mutants ordered by selection method for the binders identified for further lead optimization. Each 96 well plate includes the parent sequence in the order and a negative binding control mAb is added during processing for SPR. HC = Heavy Chain; LC = Light Chain.

| Selection Method | COL6A3 | CHI3L2 | AZGP1 | IL36RA |
| --- | --- | --- | --- | --- |
| Alanine scan | 20 | 25 | 23 | 31 |
| Germline reversion | 6 | 8 | 19 | 14 |
| Structural alone | 27 | 41 | 17 | 28 |
| Sequence alone | 41 | 13 | 33 | 20 |
| Structural + Sequence | 0 | 7 | 2 | 1 |
| N Total Mutants | 94 | 94 | 94 | 94 |
| Mutation Positions (HC:LC) | 70:24 | 48:46 | 49:45 | 40:54 |

#### Library Creation for Second Round of Optimization

Binding data were collected on the first-round libraries and used to identify mutants that outperformed the parental sequence on affinity. Those improved variants were then combined into higher-order mutation variants and re-scored with AbsciBind. An additive model of binding affinity (e.g. the predicted improvement in binding affinity is simply the sum of log affinity improvement of the individual mutants that make up a combination) was considered as well. Library selection balanced favoring designs with the fewest mutations but high predicted improvements in binding affinity with designs that increased AbsciBind score despite containing many mutations. 91 higher-order mutants and two controls (the original *de novo* binder and the best single mutant from the first round of lead optimization) were ordered for the second-round libraries.

### 7.8 *In vitro* Methodology

#### 7.8.1 Antibody Production

Up to 95 monoclonal antibodies per target were produced by GenScript (Piscataway, NJ, USA) using a Chinese hamster ovary (CHO) cell-based expression system at a 1mL culture volume scale. Antibodies were purified from culture supernatants utilizing protein A magnetic beads and supplied in a buffer containing sodium acetate, 0.2M L-arginine, at pH 5.5. Final antibody concentrations ranged from 2.6mg/mL to 0.05 mg/mL. Endotoxin levels were confirmed to be below 0.020 EU/mg for all samples. For hit validation studies, selected antibody sequences were expressed as Fabs by GenScript using a CHO-based system at a 30mL culture volume scale and as mAbs by WuXi Biologics (Wuxi, Jiangsu, China) using a CHO-based system at a 20mL culture volume scale. For mAbs produced by WuXi Biologics, purification was performed from culture supernatants via protein A affinity chromatography, and antibodies were formulated in sodium acetate buffer containing 20 mM histidine, 150 mM NaCl, at pH 5.5. In certain cases, formulations were further supplemented with 150 mM arginine and 60 mM succinic acid. Antibody concentrations ranged from 10 mg/mL to 0.15 mg/mL, and endotoxin levels were maintained below 0.020 EU/mg for all samples.

#### 7.8.2 Surface Plasmon Resonance (SPR)

Binding by SPR was assessed using LSA^XT^ instruments (Carterra, Salt Lake City, UT).

##### Primary Hit Screening

Binding between designed antibodies and their intended targets was assessed by SPR. Antigens were buffer-exchanged using Zeba Desalting Spin Columns (ThermoFisher, Cat. No. 89890) and transferred into 96 deep-well plates at a starting concentration of 2µM, then serially diluted 4-fold for 6 steps in 1x HBSTE-BSA assay buffer (10 mM HEPES pH 7.4, 150mM NaCl, 3mM EDTA, 0.05% Tween-20 + 0.5g/L BSA). Target-designed mAbs and four framework controls were immobilized onto a SAHC30M chip (Carterra, Cat. No. 4294) coated with 20µg/mL CaptureSelect Biotinylated Anti-IgG Human Fc antibody (ThermoFisher, 7103262100). Samples were immobilized for ten minutes in 1x HBSTE assay buffer (10 mM HEPES pH 7.4, 150 mM NaCl, 3 mM EDTA, 0.05% Tween-20) on the chip, followed by a five-minute injection of antigen, and ten-minute injection of 1x HBSTE-BSA assay buffer to measure rate of association (k_on_) and dissociation (k_off_) between antigen and antibody. Each antibody-antigen pair was run in duplicate, with antibodies re-immobilized on the sensor chip surface between prints with a 2 x 120 second regeneration injection of 10mM Glycine HCl pH 2.0 (Carterra, Cat. No. 3640). Kinetics analysis software (v2.0, Carterra) was used to analyze datasets.

Sensorgrams with signal output of Response Units (RU) < 5 were automatically excluded, and further exclusions were made manually for sensorgrams with signal < 10 RU. For any sensorgram to be further evaluated, signal for the 2 µM concentration must have been > 10 RU with signal (> 5 RU) in at least the second-highest concentration (500 nM). Sensorgrams were then visually inspected for curvature, convergence and dissociation according to evaluation criteria as part of standard analysis. Sensorgrams depicting high degrees of linearity in both the association and dissociation phases, as well as characteristic slow k_on_ and k_off_ were determined to be non-specific. By contrast, true hit sensorgrams displayed curvature in both the association and dissociation phases, with faster k_on_ and k_off_. To be considered a true hit, a design must have also bound specifically to the target it was designed for and show a lack of binding to off-targets.

##### Off Target Screening

Antibody hits were tested against two unrelated targets, TL1A and PRLR, to assess target specificity. Antigens were assessed against a dilution series starting at 1 uM and serially diluted 4-fold in 1X HBSTE + BSA assay buffer for three dilution points. Sensorgrams with signal < 10 RU were considered non-binding.

##### Lead Optimization Screening

For lead optimization libraries, antibody and antigen concentrations were adjusted to bring signal in range for affinity measurement. For the first round of lead optimization, antibodies designed against COL6A3 were immobilized at 0.5 µg/mL and assessed against a dilution series of COL6A3 starting at 2 µM and serially diluted 3-fold in 1x HBSTE-BSA assay buffer (10 mM HEPES pH 7.4, 150 mM NaCl, 3 mM EDTA, 0.05% Tween-20 + 0.5 g/L BSA) for six dilution points. Antibodies designed against AZGP1, CHI3L2 and IL36RA were immobilized at 1 µg/mL and assessed against a dilution series of their respective antigens starting at 2 µM and serially diluted 2-fold in 1x HBSTE-BSA assay buffer for six dilution points.

Antibodies designed against COL6A3 were immobilized at 0.5 µg/mL and assessed against a dilution series of COL6A3 starting at 300 nM and serially diluted 2-fold in 1X HBSTE-BSA assay buffer for eight dilution points. Antibodies designed against IL36RA were immobilized at 0.1 µg/mL and assessed against human (333 nM, 3-fold, 6 points) and mouse IL36RA (667 nM, 3-fold, 7 points) in 1x HBSTE-BSA assay buffer.

During kinetic analysis of the COL6A3 and AZGP1 antibody designs, affinity was calculated after cropping sensorgrams to 375 seconds and 600 seconds, respectively, due to biphasic binding responses. For lead optimization libraries, data were fit to a 1:1 Langmuir model to calculate affinity.

**Supplementary Figure 18.**
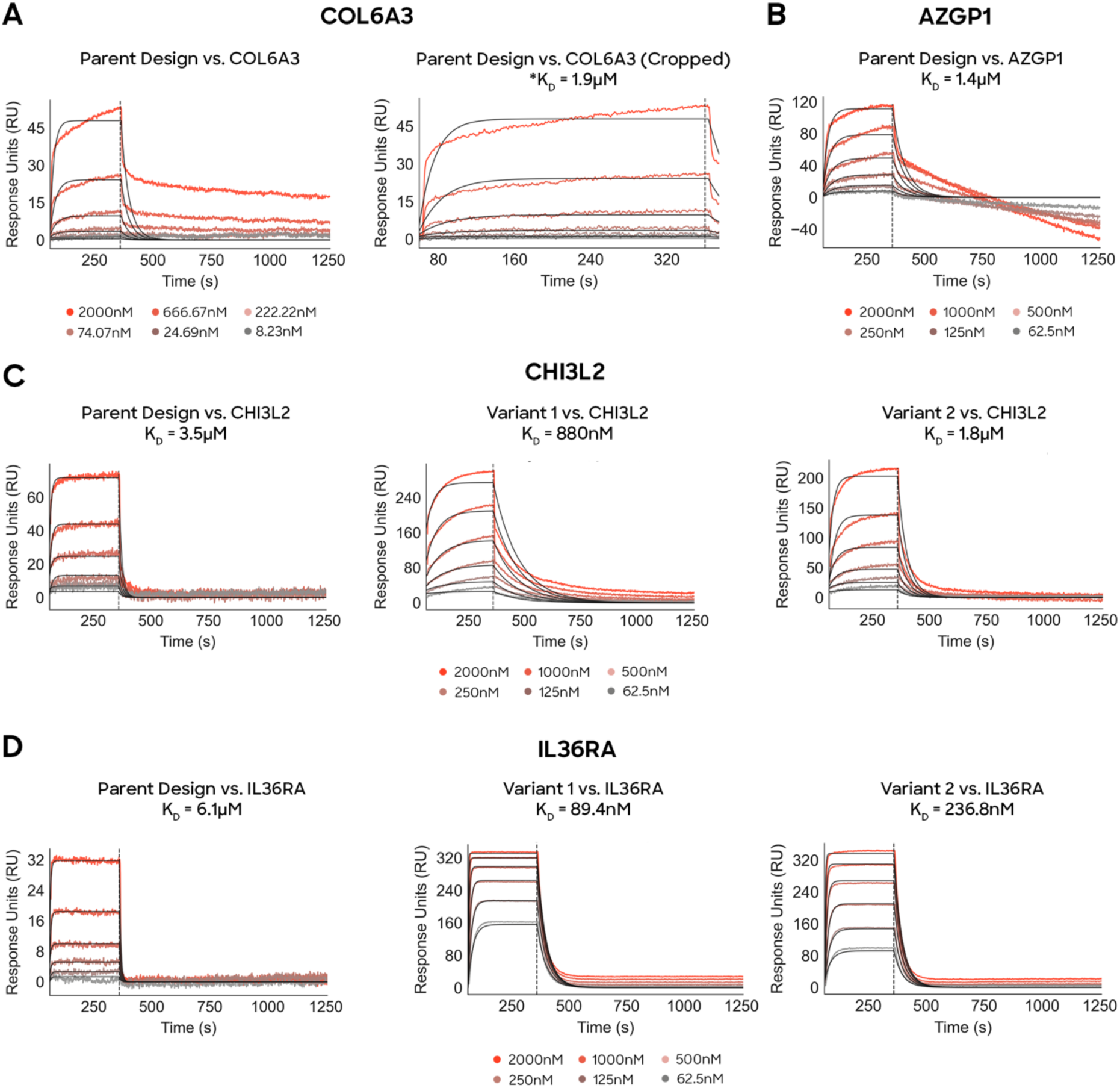
Per-target SPR sensorgrams including measured binding affinities for Origin-1 parent and optimized variant designs. (A) Left: Parent design against COL6A3 binds to COL6A3. Right: Binding affinity is measured from cropped SPR sensorgram reflecting binding of COL6A3 Parent Design against COL6A3. * indicates that binding affinity is computed from cropped sensorgram (B) Parent design against AZGP1 binds to AZGP1. (C) Left: Parent design against CHI3L2 binds with micromolar affinity. One round of AI-based affinity maturation leveraging AbsciBind improves binding affinity by approximately 4X (Middle) and 2X (Right). (D) Left: Parent design against IL36RA binds to IL36RA with micromolar affinity. One round of AI-based affinity maturation leveraging AbsciBind improves binding affinity by approximately 68X (Middle) and 26X (Right).

#### 7.8.3 Biolayer Interferometry (BLI)

A BLI kinetics assay was used to confirm binding results that were observed upstream by SPR. The Gator® label-free bioanalysis system, which includes the Gator® Prime instrument, biosensor probes, and a computer with integrated software, was used to measure the binding kinetics between antibodies and antigens. All steps in the instrument were performed at 25 °C and an orbital shaking speed of 1,000 RPM. All reagents were formulated in 1x HBSTE-BSA (1x 10 mM HEPES pH 7.4, 150 mM NaCl, 3 mM EDTA, 0.05% Tween 20, 0.5 mg/mL BSA) assay buffer. All sensors were rehydrated in buffer for a minimum of ten minutes before beginning the assay. Prior to each kinetic measurement, the Anti-Human Fab (Gator; Cat. No. 160013) biosensor probes were dipped into the assay buffer for 60 seconds to establish a baseline. For kinetics measurement, 5 µg/mL of each antibody was immobilized onto the biosensor probes for 120 seconds. Following antibody immobilization, the biosensor probes were dipped into the assay buffer for 60 seconds to assess baseline drift and evaluate the antibody loading level. Subsequently, the probes were exposed to serially diluted antigen solutions (ranging from 2000 nM to 31.25 nM in 2-fold dilutions) for five minutes to monitor real-time association kinetics. This was followed by a ten-minute dissociation phase, during which the probes were transferred to antigen-free assay buffer (HBSTE-BSA) to assess the rate of antibody-antigen complex dissociation from the biosensor surface. Kinetic data were analyzed using the GatorOne analysis software (v2.17.7, Gator Bio). Quality of fit was assessed by using the value of *R*^-^ > 0.95 with manual inspection of sensorgram curvature. Kinetics sensorgrams were plotted in GraphPad Prism 10.0.

#### 7.8.4 Antibody-Antigen Complexation

Antibody binders were reformatted as Fabs and combined with respective antigen at a 1.3:1 antigen:Fab molar ratio and purified by Size-Exclusion Chromatography using a Superdex200 Increase 10/300 column (Cytiva). Chromatograms were normalized and plotted against Fab/antigen alone to compare differences in retention volume.

We show gels related to complexation chromatograms in Supplementary Figure 19.

**Supplementary Figure 19.**
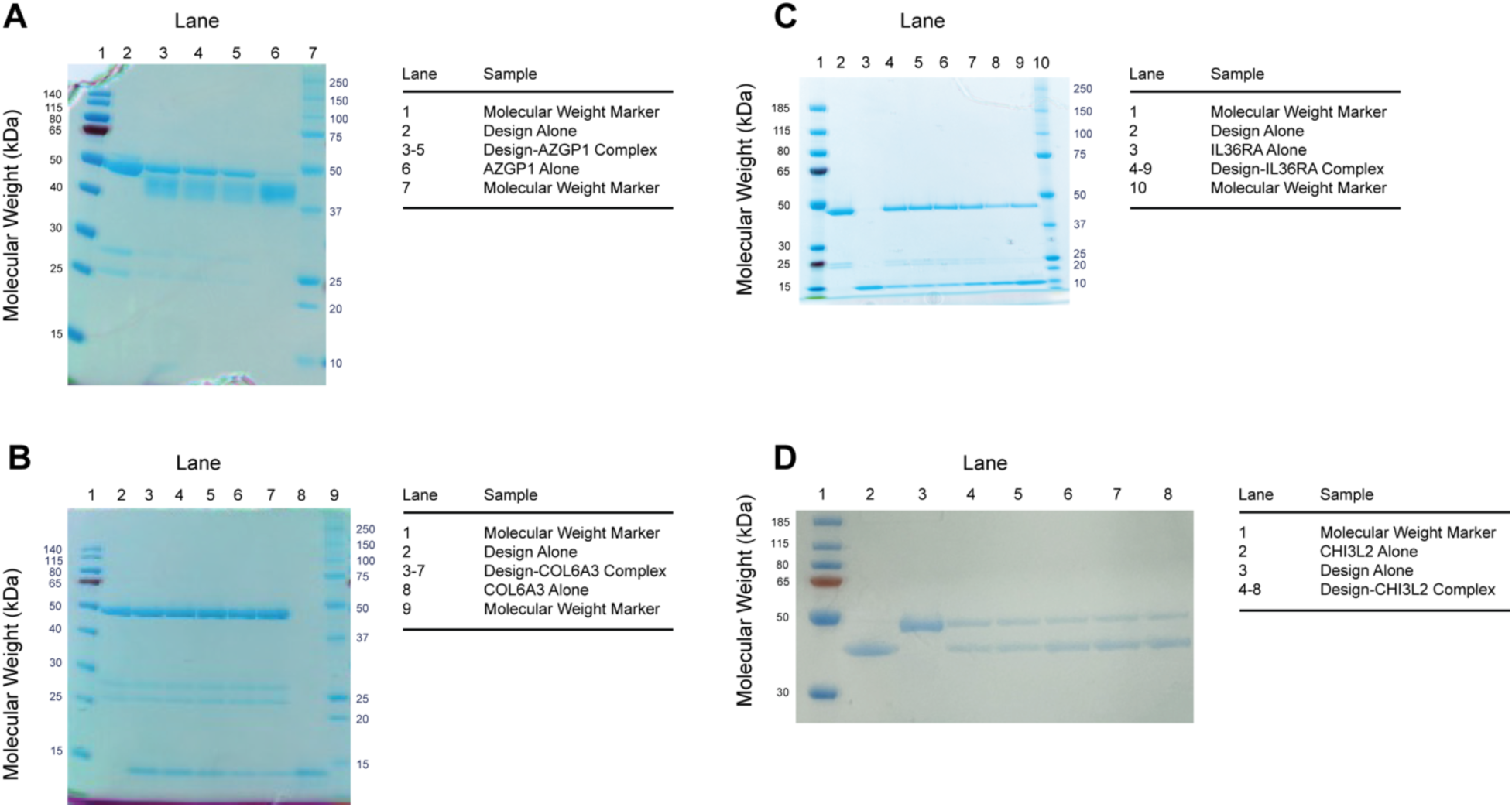
SDS-PAGE gels for validated antigen-Fab combinations. (A) SDS-PAGE gel with lanes corresponding to AZGP1 alone, Fab alone, and selected AZGP1-Fab complex fractions from chromatogram seen in **Figure 10**. (B) SDS-PAGE gel with lanes corresponding to COL6A3 alone, Fab alone, and selected COL6A3-Fab complex fractions from chromatogram seen in **Figure 9**. (C) SDS-PAGE gel with lanes corresponding to IL36RA alone, Fab alone, and selected IL36RA-Fab complex fractions from chromatogram seen in **Figure 12**. (D) SDS-PAGE gel with lanes corresponding to CHI3L2 alone, Fab alone, and selected CHI3L2-Fab complex fractions from chromatogram seen in **Figure 11**.

#### 7.8.5 Developability Assessments

##### Antibody Quality Assessment

Antibody quality assessments were performed by SEC, non-reduced CGE (NR-CGE) or Microchip-CGE (MCGE), and intact mass spectrometry. Concentration of mAbs was determined by A280 using the SoloVPE instrument (CTech^TM^) and each antibody’s calculated extinction coefficient [66]. Aggregation determination by SEC was performed using HPLC 1260 Infinity II (Agilent) with mobile phase 1X PBS, pH 7.4 and separation was done on a 30 cm TSKgel UP-SW2000 (Tosoh) column. Fragmentation evaluation by NR-CGE was performed using CESI 8000 Plus (Sciex) with a high-speed setup method for separation. Alternatively, for high-throughput analysis, fragmentation was performed by MCGE using a LabChip GX Touch HT (Revity) and the protein express reagent kit (Revity). Confirmation of identity was done by intact mass spectrometry using a reversed phase HPLC 1290 Infinity II (Agilent) connected to a TripleTOF 6600+ MS System (Sciex). Intact mass data analysis was performed using PMI-Byos software v4.5-53 (Protein Metrics). All assays contained a system suitability check in each run or plate as determined by the performance of trastuzumab (SEC, NR-CGE, MCGE), BSAA standard (concentration of mAbs) or NIST mAb (intact mass).

##### Antibody Developability Assays

Particle size and thermal stability of the antibodies were performed by dynamic light scattering (DLS) and nano differential scanning fluorimetry (nanoDSF) using Prometheus Panta instrument (NanoTemper). DLS measurements were acquired at 25°C under high sensitivity mode. Thermal unfolding profiles were subsequently recorded from 25°C to 90°C with a temperature ramp of 0.5°C per minute. Data was processed and analyzed using Prometheus Panta software version 1.1.

Antibody self-association, polyreactivity, and hydrophobicity were analyzed using affinity capture self-interaction nanoparticle spectroscopy (AC-SINS), ELISA-based assays, and hydrophobic interaction chromatography (HIC), respectively. AC-SINS was performed according to the method reported before [67]. Polyreactivity was assessed as anti-DNA and anti-insulin ELISAs according to procedure described previously [68]. Data reporting for polyreactivity was performed in a novel manner to account for plate-to-plate variability and the four dilution levels used in the study. Molecules were run at 0.08, 0.4, 2.0 and 10.0 µg/mL against immobilized DNA or Insulin, with the resulting absorbance values divided by the value of the blank (buffer) for a score. This score was then normalized for each dilution level using the maximum value on the plate (rescored 11) and blank value on the plate (rescored 1) to reduce plate to plate variability affecting scoring. These normalized scores were then added for all four dilution levels for a Range Normalized Summation (RNS) reported as the total polyreactivity result. HIC analyses were performed according to a procedure detailed in Jain et al. [69]. HIC reporting was done as relative retention time compared to trastuzumab (Sample retention time/trastuzumab retention time) All the assays were evaluated for system suitability checks in each run or plate as determined by the performance of a negative control, trastuzumab for all assays, and at least one positive control such as Infliximab for AC-SINS, Bococizumab and Briakinumab for polyreactivity, or BSA and Insulin for HIC. For DLS and nanoDSF, standard particles solution and lysozyme standard were used for system suitability checks in each run, respectively.

#### 7.8.6 Cryogenic Electron Microscopy (Cryo-EM)

##### Data Acquisition

For the AZGP1-Design complex, fractions purified as described above were combined and concentrated down to 1 mg/mL before snap freezing. For the COL6A3-Design complex, fractions purified as described above were combined at a 1:1:1 molar ratio with modified variants of an anti-Kappa VHH (Q5V, Q113K, Q116P, referred as Nanodaptor) and NabFab (S123E, Q199R, referred as Kappabulk). Relevant fractions were pooled and concentrated down to 4.5mg/mL before snap freezing. For the IL36Ra-Design complex, purified fractions were combined at a 1:1:1 molar ratio with modified anti-Kappa VHH and NabFab proteins. Relevant fractions were pooled and concentrated down to 9.4mg/mL before snap freezing.

For cryo-EM grid preparation, 2.5µL of purified complexes at concentrations of 0.4mg/ml to 0.6mg/ml were applied to glow-discharged Cu 300 mesh holey carbon grids (Quantifoil R1.2/1.3). The grids were blotted and plunge-frozen in liquid ethane using a Vitrobot Mark IV (ThermoFisher Scientific).

For the AZGP1-Design complex dataset, 9435 movies were collected on a Titan Krios transmission electron microscope operated at 300kV and equipped with a Falcon4i direct electron detection detector. Data were acquired at a nominal magnification of 130,000, corresponding to a calibrated pixel size of 0.932 Å. Each movie consisted of 40 frames resulting in a total accumulated electron dose of approximately 49 e^-^/Å^2^.

For the COL6A3-Design complex dataset, 9502 movies were collected on a Titan Krios transmission electron microscope operated at 300 kV and equipped with a K3 direct electron detection detector. Data were acquired at a nominal magnification of 105,000, corresponding to a calibrated pixel size of 0.824 Å. Each movie consisted of 40 frames resulting in a total accumulated electron dose of approximately 48 e^-^/Å^2^.

For the IL36RA-Design complex dataset, 9000 movies were collected on a Titan Krios transmission electron microscope operated at 300 kV and equipped with a Falcon4i direct electron detection detector. Data were acquired at a nominal magnification of 130,000, corresponding to a calibrated pixel size of 0.932 Å. Each movie consisted of 40 frames resulting in a total accumulated electron dose of approximately 50 e^-^/Å^2^

##### Image Processing and 3D Reconstruction

Movie stacks were motion-corrected and dose-fractionated at the micrograph level using MotionCor2 [70] to correct for beam-induced motion and stage drift. Non–dose-weighted micrographs were used for contrast transfer function (CTF) estimation with CTFFIND [71]. Subsequent data processing was performed using RELION 4.0 [72].

For the AZGP1-Design complex, 5.8 million particles were auto-picked and subject to multiple rounds of 2D classification. Following 2D classification, approximately 2.0 million particles were selected for ab initio reconstruction. From five resulting 3D classes, one class comprising 435,000 particles was subjected to 3D refinement, CTF refinement, Bayesian polishing, and post processing. The final reconstruction reached a global resolution of 3.1 Å, as determined by the gold-standard Fourier shell correlation (FSC) 0.143 criterion (**Supplementary Figure 20**).

For the COL6A3-Design-Nanodaptor-Kappabulk complex, approximately 4.1 million particles were auto-picked and subject to several rounds of 2D classification. Following 2D classification, approximately 1.0 million particles were used for ab initio reconstruction. From five 3D classes, one class comprising 494,000 particles was selected for 3D refinement, CTF refinement, Bayesian polishing, and post-processing, yielding a reconstruction with a global resolution of 3.0 Å (FSC 0.143 criterion). To further improve map quality at the binding interface, a focused local refinement on the COL6A3–Fab region was performed, resulting in a local resolution of 2.9 Å (**Supplementary Figure 21**).

For the IL36RA-Design-Nanodaptor-Kappabulk complex, approximately 3.2 million particles were auto-picked and subject to several rounds of 2D classification. Following 2D classification, approximately 1.0 million particles were used for ab initio reconstruction. From five 3D classes, one class comprising 566,000 particles was selected for 3D refinement, CTF refinement, Bayesian polishing, and post-processing, yielding a reconstruction with a global resolution of 3.1 Å (FSC 0.143 criterion). To further improve map quality at the binding interface, a focused local refinement on the IL36RA–Fab region was performed, resulting in a local resolution of 3.1 Å (**Supplementary Figure 22**).

##### Model Building and Refinement

The starting model for COL6A3, AZGP1 and IL36RA were derived from PDB entries 1KTH, 1T7V and 4POJ respectively. Initial atomic models of the corresponding complexes were generated by autobuilding directly from the Fab sequences using CryFold, followed by manual building of missing/incorrect regions in Coot. Final models were completed by iterative rounds of manual model adjustment in Coot and automated real-space refinement using PHENIX.

Structure visualizations in this work were made possible by PyMOL [75] and Mol* [76].

**Supplementary Figure 20.**
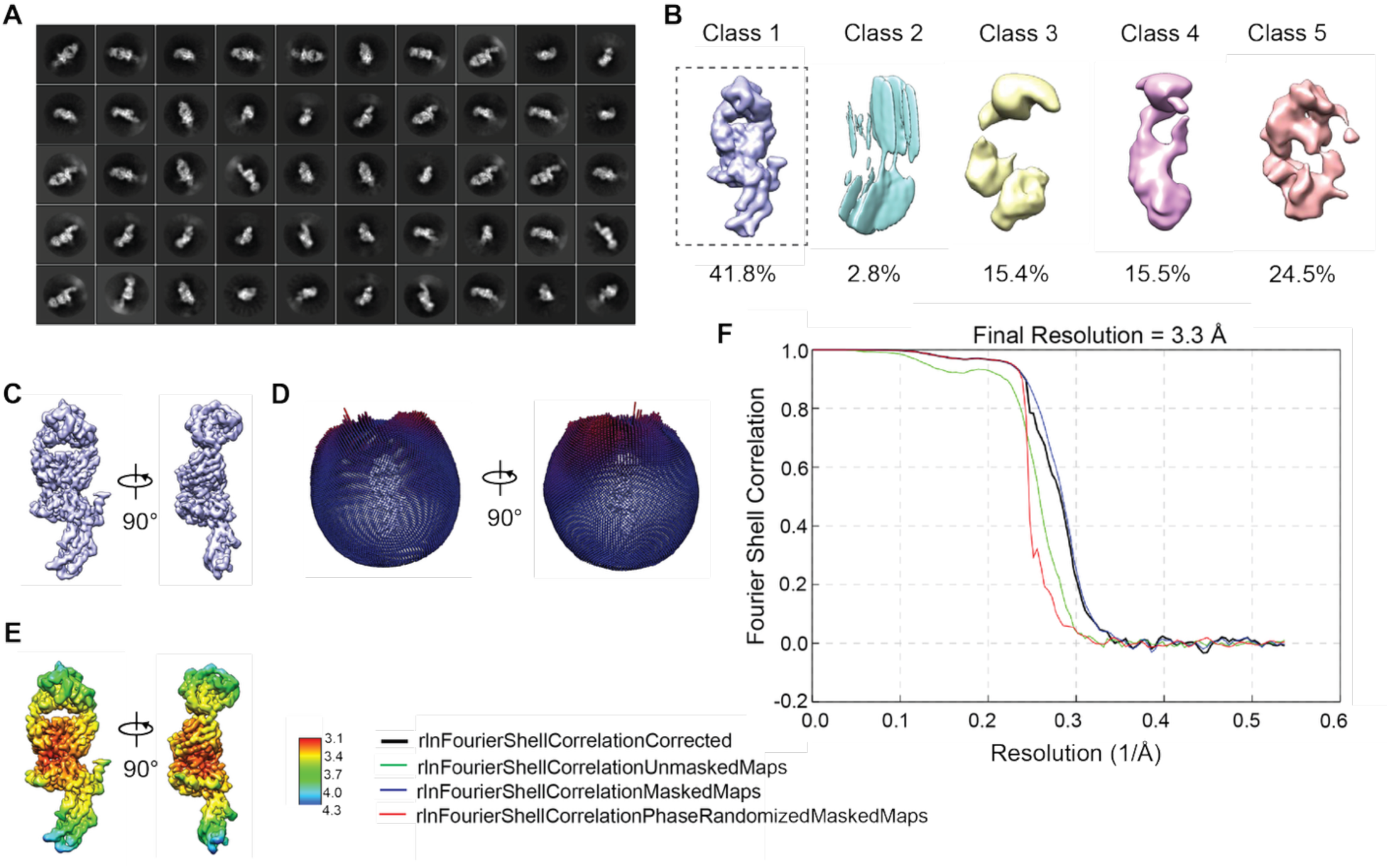
CryoEM processing workflow of AZGP1-Design structure. (A, B) Representative 2D class averages and 3D classes. (C-E) Global 3D reconstruction, angular distribution of particle orientations, local resolution estimations and sharpened map. (F) Gold-standard FSC plots for the AZGP1-Design complex.

**Supplementary Figure 21.**
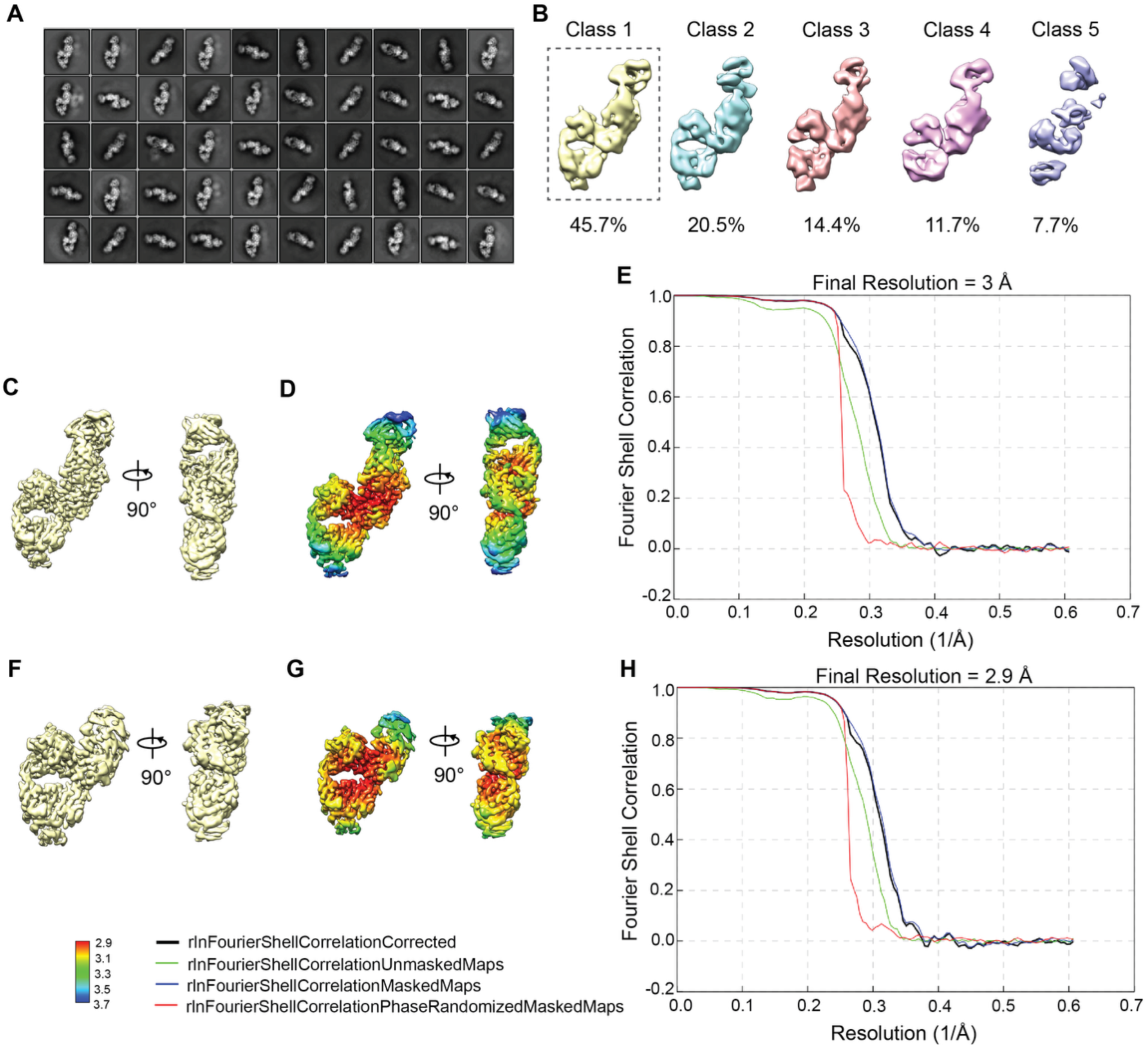
CryoEM processing workflow of COL6A3-Design-Anti-Kappa VHH-NabFab structure. (A, B) Representative 2D class averages and 3D classes. (C-E) 3D reconstruction, local resolution estimations and gold-standard Fourier shell correlation (FSC) plots for the COL6A3 complex global map. (F-H) Locally refined 3D reconstruction map, local resolution estimations and gold-standard FSC plots for the COL6A3-Design region.

**Supplementary Figure 22.**
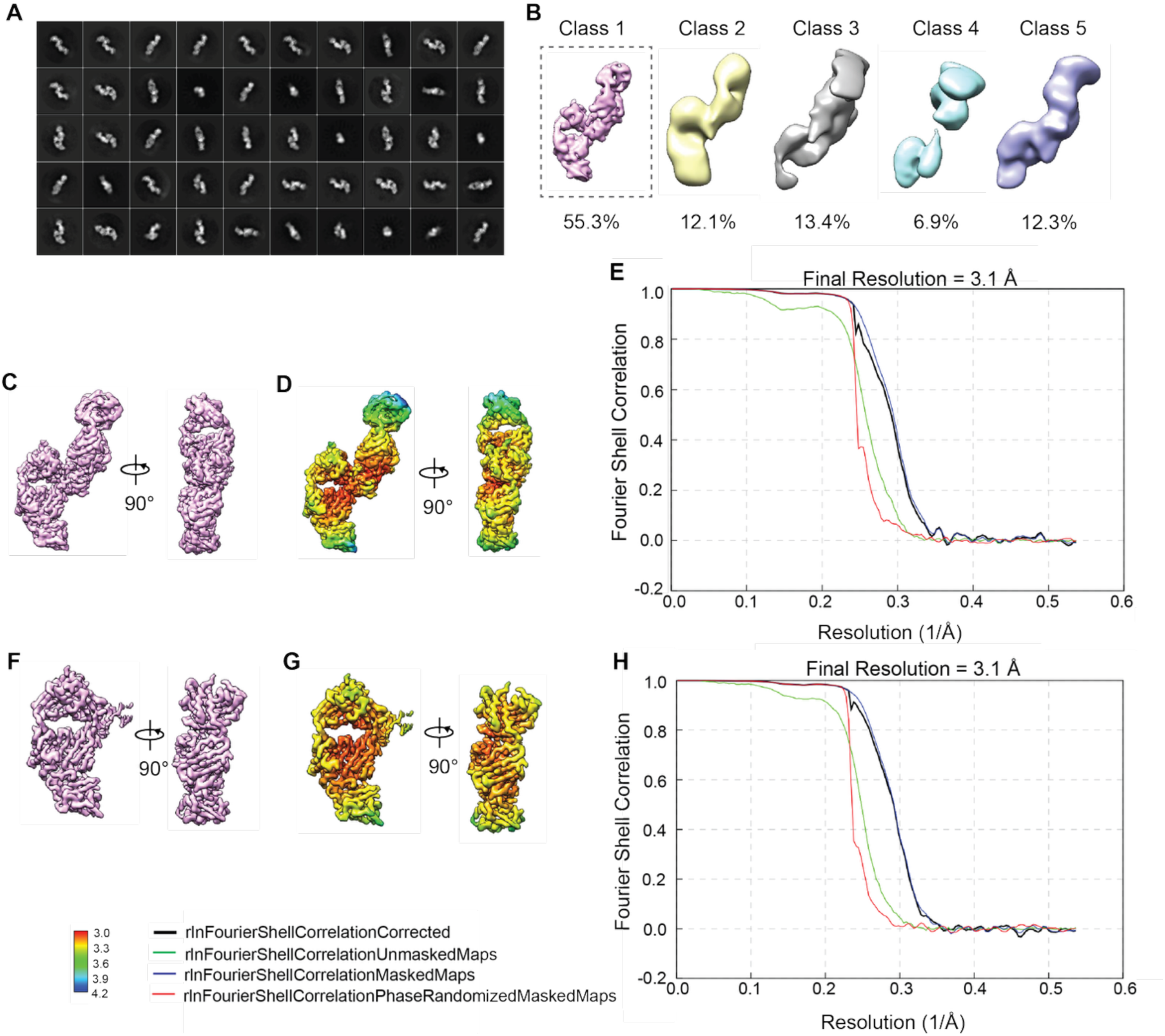
CryoEM processing workflow of IL36RA-Design-Anti-Kappa VHH-NabFab structure. (A, B) Representative 2D class averages and 3D classes. (C-E) 3D reconstruction, local resolution estimations and gold-standard Fourier shell correlation (FSC) plots for the IL36RA complex global map. (F-H) Locally refined 3D reconstruction map, local resolution estimations and gold-standard FSC plots for the IL36RA-Design region.

#### 7.8.7 Functional Assessment

##### Testing Functional Activity of Identified IL36RA Binders in Human Cells

The IL36 cytokine axis is a tightly regulated inflammatory signaling pathway in which the agonist ligands IL36α, IL36β, and IL36γ trigger IL36 receptor activation and downstream MAPK and NF-κB signaling [49]. In physiological settings, this response is restrained by IL36RA, an endogenous receptor antagonist that competes with agonist ligands and suppresses signaling output.

To functionally characterize antibodies directed against IL36RA, we used a reporter-based system in which IL36 receptor signaling is converted into a quantitative, secreted readout. In HEKBlueIL36 cells, activation of AP1 and NF-κB drives expression of secreted embryonic alkaline phosphatase (SEAP), enabling pathway activity to be monitored directly in cell culture supernatants using a simple colorimetric assay. In the present study, IL36γ is used to stimulate the pathway, IL36RA is added to impose pharmacologic inhibition, and test antibodies are evaluated for their ability to neutralize IL36RA activity and thereby restore IL36γ-dependent signaling, as measured by SEAP production.

HEK-Blue-IL36 cells were seeded into 96-well tissue-culture plates at a density of 25,000 cells/well in the presence of IL36γ (2 pM), IL36RA (50 nM), and increasing concentrations of test antibody. Test antibodies were prepared as threefold serial dilutions spanning 1 µM to 1 nM.

Cells were incubated for 24 h at 37°C, 5% CO₂, under humidified conditions. Following incubation, 20 µL of conditioned medium was transferred and mixed with 180 µL of QuantiBlue (InvivoGen) working solution in a clear 96-well plate and incubated at room temperature for 60 min. Absorbance was measured at 620 nm using a SpectraMax i3x plate reader. Raw data were processed in Excel, and dose–response curves were fitted by nonlinear regression using Prism 10 (GraphPad).

##### Testing Functional Activity of Identified IL36RA Binders in Mouse Cells

CT-26.WT murine colorectal carcinoma cells secrete CCL2 (also known as MIP-1) in response to IL36β stimulation. Engagement of the IL36 receptor by its ligands (IL36α, IL36β, and IL36γ) activates the MAPK and NFκB signaling pathways. IL36RA acts as a receptor antagonist, inhibiting IL36 signaling.

We developed an assay that determines whether an antibody targeting IL36RA can restore IL36 signaling (blocked by IL36RA) by quantifying CCL2 secretion into the culture medium.

In brief, CT-26.WT cells were seeded in 96-well plates at 150,000 cells/well. Cells were cultured with 10 nM mIL36β and 300 nM mIL36Ra in the presence of serially diluted test antibody (three-fold dilutions from 2 µM to 34 pM). Plates were incubated for 24 hours at 37°C in 5% CO₂ and high humidity.

Following incubation, 50 µl of conditioned medium from each well was collected for quantification of CCL2 using a commercial ELISA kit (R&D Systems), according to the manufacturer’s instructions. Absorbance at 450 nm was measured with a SpectraMax i3x plate reader. Data were processed in Microsoft Excel and analyzed by non-linear regression using Prism 10 (GraphPad).

#### 7.8.8 CHI3L2 Antigen Generation and Validation

SPR, BLI, and aSEC complexation experiments were initially run with CHI3L2 from R&D Systems. To further validate binding, we produced CHI3L2 internally and re-ran SPR and aSEC experiments for our *de novo* hit. We found that SPR results were consistent between both sources of CHI3L2. The data reported in **Figure 11** show SPR and aSEC against internally produced antigen and BLI against antigen from R&D Systems. Below we describe how CHI3L2 was internally produced.

CHI3L2 (aa26-390) was cloned into gWIZ containing an N-terminal Murine IgG heavy chain signal peptide and a C-terminal 6xHis tag (Twist Biosciences). Expi293F cells were transfected at a density of 3×10^6^ cells/mL using 1 ug/mL of plasmid DNA, PEIStar transfection reagent (Bio-Techne) at a 5.3:1 PEI:DNA ratio, and an in-house enhancer formulation containing Glucose (46 mM), Valproic Acid (5 mM), and Sodium Propionate (6.9 mM). After 6 days the supernatant was harvested and batch bound with Ni-NTA resin (Cytiva) overnight before purification. Samples were concentrated, polished using Size Exclusion Chromatography in 50 mM HEPES, pH 7.4 + 150 mM Sodium Chloride, and snap frozen for downstream experiments.

##### Comparing Commercially Purchased and Internally Produced CHI3L2 by SPR

Antibody binders for CHI3L2 were immobilized at 1 ug/mL and assessed against a dilution series of commercially purchased CHI3L2 (R&D Systems 5112-CH-050) and internally produced CHI3L2 starting at 1 µM and serially diluted 3-fold in 1X HBSTE-BSA assay buffer for six dilution points. Data were fit to a 1:1 Langmuir model to calculate affinity.

